# The effects of caloric restriction on adipose tissue and metabolic health are sex- and age-dependent

**DOI:** 10.1101/2022.02.20.481222

**Authors:** Karla J. Suchacki, Ben J. Thomas, Yoshiko Matsumoto Ikushima, Kuan-Chan Chen, Claire Fyfe, Adriana A.S. Tavares, Richard J. Sulston, Andrea Lovdel, Holly J. Woodward, Xuan Han, Domenico Mattiucci, Eleanor J. Brain, Carlos J. Alcaide-Corral, Hiroshi Kobayashi, Gillian A. Gray, Phillip D. Whitfield, Roland H. Stimson, Nicholas M. Morton, Alexandra M. Johnstone, William P. Cawthorn

## Abstract

Caloric restriction (CR) is a nutritional intervention that reduces the risk of age-related diseases in numerous species, including humans. CR’s metabolic effects, including decreased fat mass and improved insulin sensitivity, play an important role in its broader health benefits. However, the extent and basis of sex differences in CR’s health benefits are unknown. We found that 30% CR in young (3-month-old) male mice decreased fat mass and improved glucose tolerance and insulin sensitivity, whereas these effects were blunted or absent in young female mice. Females’ resistance to fat and weight loss was associated with decreased lipolysis, lower systemic energy expenditure and fatty acid oxidation, and increased postprandial lipogenesis compared to males. Positron emission tomography-computed tomography (PET/CT) with ^18^F-fluorodeoxyglucose (^18^F-FDG) showed that peripheral glucose uptake was comparable between sexes. Instead, the sex differences in glucose homeostasis were associated with altered hepatic ceramide content and substrate metabolism: compared to CR males, CR females had lower TCA cycle activity but higher blood ketone concentrations, a marker of hepatic acetyl-CoA content. This suggests that males use hepatic acetyl-CoA for the TCA cycle whereas in females it accumulates, thereby stimulating gluconeogenesis and limiting hypoglycaemia during CR. In aged mice (18-months old), when females are anoestrus, CR decreased fat mass and improved glucose homeostasis to a similar extent in both sexes. Finally, in a cohort of overweight and obese humans CR-induced fat loss was also sex- and age-dependent: younger females (<45 years) resisted fat loss compared to younger males while in older subjects (>45 years) this sex difference was absent. Collectively, these studies identify age-dependent sex differences in the metabolic effects of CR and highlight adipose tissue, the liver and oestrogen as key determinants of CR’s metabolic benefits. These findings have important implications for understanding the interplay between diet and health and for maximising the benefits of CR in humans.

**HIGHLIGHTS:**

- Caloric restriction (CR) decreases fat mass and improves glucose homeostasis in young male mice, but young females resist these effects.
- CR females resist lipolysis, decrease energy expenditure and increase postprandial lipogenesis more than CR males, explaining how females resist fat loss.
- Sex differences in glucose homeostasis are associated with altered hepatic metabolism and gluconeogenesis, without marked differences in peripheral glucose uptake.
- CR’s effects on fat loss and glucose homeostasis are comparable in aged male and female mice, implicating oestrogen as the driver of the sexually dimorphic effects in young mice.
- In humans, females resist CR-induced fat loss in an age-dependent manner, further supporting the role of oestrogen in the sexually dimorphic effects of CR.

## INTRODUCTION

Caloric restriction (CR) is a therapeutic nutritional intervention involving a sustained decrease in calorie intake whilst maintaining adequate nutrition. CR extends lifespan and reduces the risk of age-related diseases in numerous species, ranging from yeast to primates (1–3). CR can have detrimental effects, including bone loss (4) and increased susceptibility to infections (5), and therefore it may not be suitable for all individuals; however, trials of CR in humans show numerous health benefits, including the prevention of cardiovascular disease, hypertension, obesity, type 2 diabetes, chronic inflammation, and risk of certain cancers (6). Thus, the ability of CR to promote healthy ageing is now also recognised in humans. In addition to these health benefits, many effects of CR reflect evolutionary adaptations that confer a survival advantage during periods of food scarcity (5). Establishing the extent and basis of CR’s effects may thereby reveal new knowledge of healthy ageing and the interplay between food, nutrition, and health.

A key contributor to the health benefits of CR is its impact on metabolic function. Ageing is characterised by hepatic insulin resistance, hyperinsulinaemia and excessive accumulation of white adipose tissue (WAT), particularly visceral WAT (7–9). The latter is coupled with adipose dysfunction, whereby WAT becomes unable to meet the demands for safe lipid storage. This results in ectopic lipid accumulation in the liver and other tissues, contributing to insulin resistance and metabolic dysregulation (7, 9). CR counteracts these effects, decreasing WAT mass, increasing fatty acid (FA) oxidation, preventing ectopic lipid deposition and enhancing insulin sensitivity and glucose tolerance (5). Notably, studies in rodents show that removal of visceral WAT, independent of CR, is sufficient to prevent insulin resistance, improve glucose tolerance and increase lifespan (10–12). Visceral WAT loss in humans is also strongly associated with CR’s metabolic benefits (13). Thus, the ability of CR to decrease visceral WAT, and the resultant improvements in hepatic insulin sensitivity, are likely central to CR’s effects on metabolic function and healthy ageing.

This importance of adiposity and metabolic function raises the possibility of sex differences in the CR response. Indeed, it is now well established that metabolic homeostasis and adipose biology differ substantially between males and females (14–17); however, most preclinical rodent studies have used males only (18), suggesting that much of the CR literature may have overlooked sex as a potential determinant of CR’s effects. Nevertheless, some clinical and preclinical CR studies from our lab and others have identified sexually dimorphic responses (19–22), whereby males lose fat mass to a greater extent than females. Oestrogens underlie many metabolic sex differences (14, 23), suggesting that oestrogen may contribute to females’ resistance to fat loss during CR. However, the extent and basis of sexual dimorphism in CR’s metabolic effects remains to be firmly established.

Herein, we first systematically review the CR literature to establish the degree to which sex has been overlooked a determinant of the CR response. To further elucidate the extent and basis of sex differences, we studied CR in male and female mice and humans at ages where oestrogen is physiologically active or absent. Together, our findings show that the CR field has largely overlooked sex differences and reveal that both mice and humans display age-dependent sexual dimorphism in the metabolic effects of CR.

## RESULTS

### Sex is routinely overlooked as a biological variable in the CR literature

One review of the recent CR literature found that most rodent studies use males only, with females typically used only to address female-specific experimental questions (19); however, the extent of this bias in earlier CR studies, and whether it applies to human CR research, has not been assessed. To address these issues, we first systematically quantified the use of males and females, and the consideration of sex as a biological variable, in mouse and human CR studies (Fig. 1, S1). We focused on research published since 2003, when the European Commission first highlighted the importance of addressing sex as a biological variable (24). We also excluded studies that necessarily focused on a single sex, such as those addressing effects of CR on female reproductive function, leaving only studies in which there is no scientific rationale for ignoring potential sexual dimorphism. We found that male-only studies predominated for mice, accounting for ∼64% of all mouse CR studies since 2003 (Fig. 1A, S1B). This is consistent with previous analyses of the more-recent CR literature (19). Fewer than 20% of studies used females only, while around 7.3% combined males and females, suggesting an assumption that sex would not influence experimental outcomes. In contrast, ∼60% of human studies combined males and females, while ∼23% used only females and ∼12% only males (Fig. 1C, S1B). Strikingly, by the end of 2021, studies that included both sexes and analysed data with sex as a variable were in a minority for both mice (∼3.4%) and humans (∼4.5%) (Fig. 1A, 1C). Moreover, the proportion of studies using each sex or combination of sexes has remained relatively constant since 2003 (Fig. 1A, 1C). Thus, efforts to increase the consideration of sex in experimental design, as promoted by the European Commission (24), Canadian Institutes of Health Research (CIHR) and National Institutes of Health (NIH) (25), appear to have had little impact on the field of CR research.

**Figure 1.**
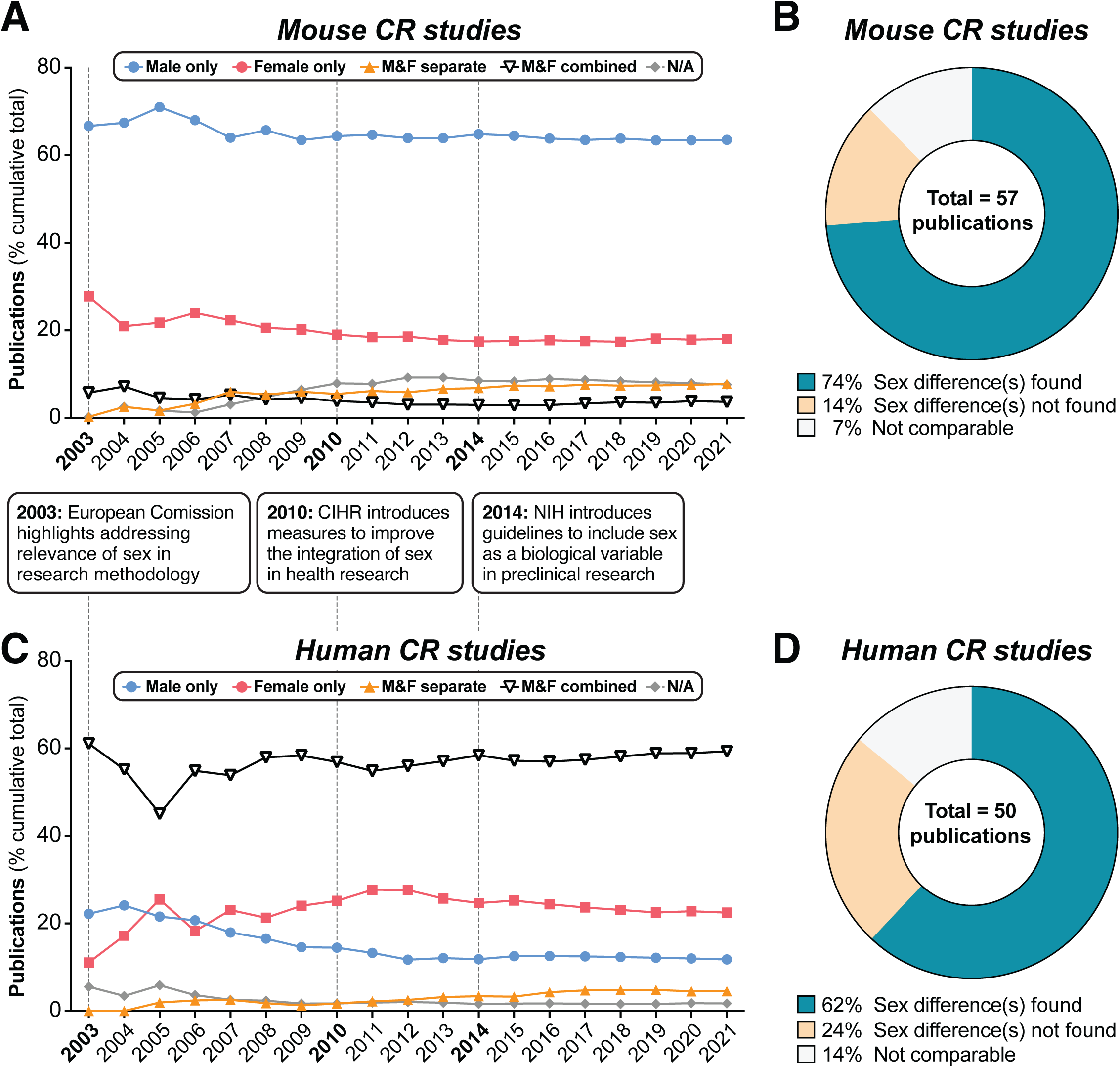
Summary of sex differences in mouse and human CR research. PubMed was searched using MeSH terms to identify research articles that studied caloric restriction *in vivo* in mice (**A,B**) or humans (**C,D**), published between 2003 and 2021. Search results were then classified into the following groups depending on the sexes included in each study: ‘Male only’ = male subjects used exclusively; ‘Female only’ = female subjects used exclusively; ‘Male & Female separate’ = male and female subjects used with data reported respective to each sex, allowing sex differences to be addressed; ‘Male & female combined’ = male and female subjects used with data from both sexes combined either in part or in full; ‘N/A’ = no sex data available. **(A,C)** Cumulative publications for studies within each group. The boxes between (A) and (C) highlight the dates of funders’ policies highlighting the importance of sex as a biological variable. (**B,D**) Pie charts of those studies in the ‘Male & Female separate’ group that considered sex a biological variable, and the proportion of these that did or did not detect sex differences. Source data are provided as a Source Data file. See also Supplementary Figure 1.

We next focused on the minority of studies that did consider sex in their experiments (“M&F separate”) and that included analysis of metabolic parameters. We found that results in ∼74% of mouse studies (Fig. 1B) and ∼62% of human studies (Fig. 1D) indicated a sex difference in the CR response. Thus, sex differences have been described in the CR literature, but the continuing dearth of studies that include both sexes suggests that this issue continues to be overlooked in the CR field.

### Females resist CR-induced weight loss and fat loss

To explore sex differences in the CR response, we assessed the effects of 30% CR, implemented from 9-15 weeks of age, in C57BL/6J and C57BL/6N mice. As expected, CR decreased body mass in both males and females (Fig. 2A) but this effect was greater in males, with ANOVA confirming a significant sex-diet interaction. This sex difference in response to CR is particularly clear when body mass is presented relative to baseline, pre-CR levels (Fig. 2B).

**Figure 2.**
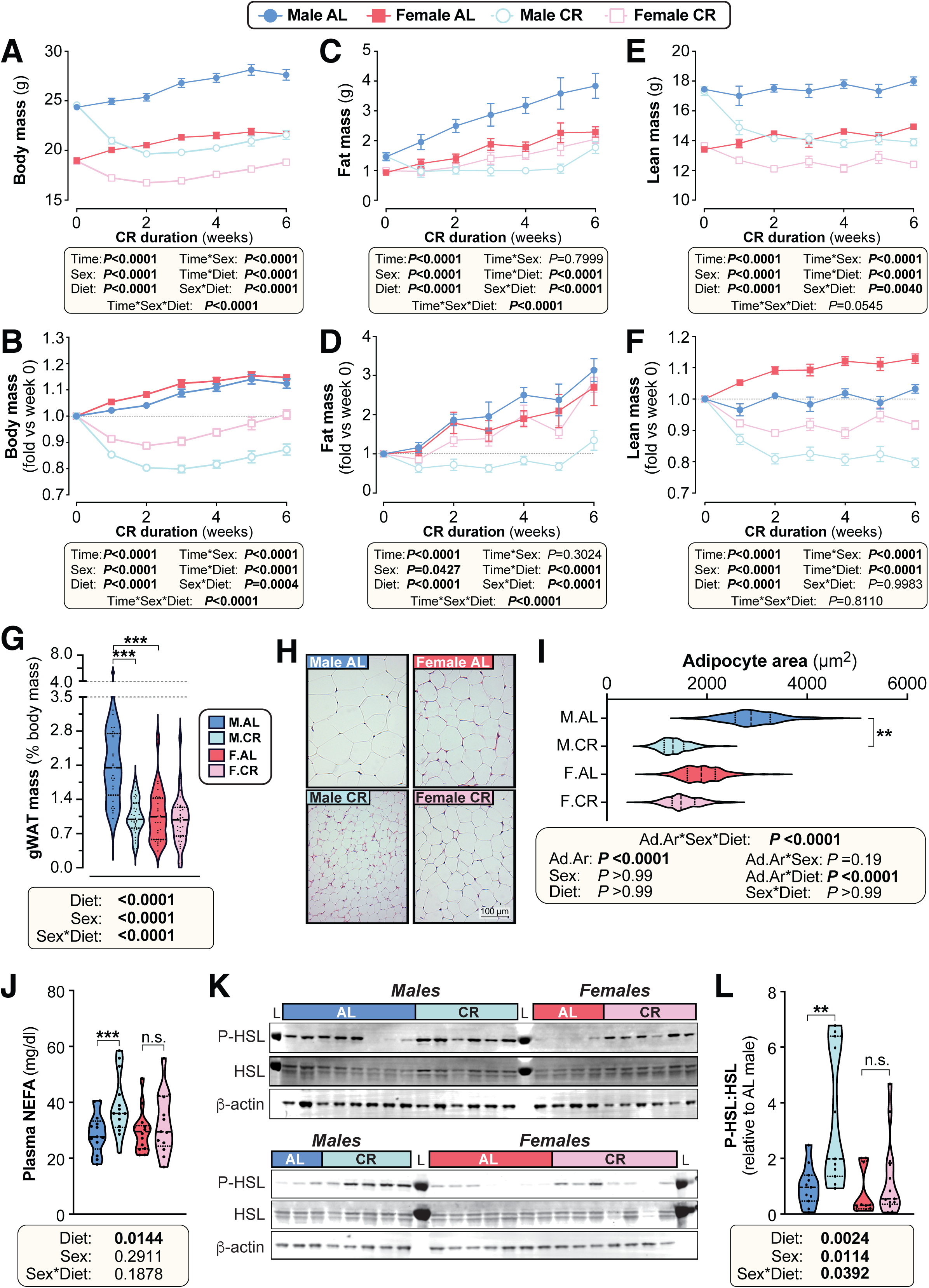
Female mice resist weight loss, fat loss and lipolysis during CR. Male and female mice on a C57BL/6NCrl or C57BL/6J background were fed *ad libitum* (AL) or a 30% CR diet from 9-15 weeks of age (0-6 weeks of CR). **(A-F)** Each week mice were weighed (A,B) and body composition was determined by TD-NMR (C-F). Body mass, fat mass and lean mass are shown as absolute masses (A,C,E) or fold-change relative to baseline (B,D,F). **(G)** The mass of gWAT (gonadal WAT) was recorded at necropsy and is shown as % body mass. **(H,I)** Micrographs of H&E-stained sections of gWAT (H) were used for histomorphometric analysis of adipocyte area (I); in (H), scale bar = 100 µm. **(J)** Plasma was sampled at 15 weeks of age and the concentration of non-esterified fatty acids (NEFA) was assayed. **(K,L)** At 10 weeks of age, during the period of maximal weight and fat loss, a separate cohort of mice were culled and iWAT was collected for analysis of the indicated proteins by Western blotting. Fluorescent Western blots (K) were quantified using LICOR software (L); L = protein ladder. Data in (H) and (K) show representative micrographs (H) and immunoblots (K). Data in (A-F) are shown as mean ±SEM of the following numbers of mice per group: *male AL*, n=42; *female AL,* n=43; *male CR,* n=44; *female CR,* n=52. Data in (G), (J) and (L) are shown as violin plots of the following numbers of mice per group: *male AL,* n=29(G), 13 (J) or 11 (L); *female AL,* n=28(G), 14 (J) or 10 (L); *male CR,* n=33 (G), 14 (J) or 11 (L); *female CR,* n=34 (G) or 13 (J,L). Data in (H-I) are shown as representative micrographs or violin plots from 5 (*male AL)* or 6 (*female AL, male CR, female CR)* mice per group. For (A-F), significant effects of diet, sex and/or time, and interactions thereof, were determined by 3-way ANOVA or a mixed-effects model. For (G), (J) and (L), significant effects of diet and/or sex were determined by 2-way ANOVA with Tukey’s or Šídák’s multiple comparisons tests. For (I), significant effects of diet and/or sex on adipocyte area (Ad.Ar) were determined using a mixed-effects model, while significant differences in median Ad.Ar between AL and CR mice were determined 2-way ANOVA with Šídák’s multiple comparisons test. *P values* from ANOVA or mixed models are shown beneath the graphs, as indicated. For (G), (I), (J) and (L), significant differences between comparable groups are indicated by * (*P*<0.05), ** (*P*<0.01) or *** (*P*<0.001). Source data are provided as a Source Data file. See also Supplementary Figures 2-4, 7 and 14.

To determine how fat and lean mass contribute to these diet and sex effects, body composition was assessed weekly using time-domain nuclear magnetic resonance (TD-NMR; Fig. 2C-F, S2A-B). CR decreased fat and lean mass in males, whereas females maintained fat mass and lost only lean mass (Fig. 2C-F). A significant sex-diet interaction was apparent for absolute fat mass (Fig. 2C) and for fat mass relative to baseline (Fig. 2D); the latter showed that CR decreased fat mass in males, whereas AL males and AL or CR females increased fat mass to a similar extent over the six-week duration (Fig. 2D). For lean mass, a significant sex-diet interaction occurred for absolute mass, with losses being greater in males than in females (Fig. 2E). However, when compared to baseline lean mass, the CR vs AL effect was similar between the sexes, in part because AL females continued to increase lean mass over time (Fig. 2F).

Given that these body composition changes coincide with changes in overall body mass, we also assessed fat and lean mass as % body mass. Males preferentially lost fat mass and preserved lean mass in response to CR, with % lean mass being greater in CR vs AL-fed males (Fig. S2A-B). In contrast, diet did not alter % fat or % lean mass in females, indicating that changes in body composition were proportionate to the overall changes in body mass (Fig. S2A-B).

To determine how these changes relate to regional adiposity, we measured adipose depot masses after 6 weeks of AL or CR diet (Fig. 2G, S2C, S3A). We found that in males but not females CR decreased the absolute mass of gonadal (gWAT), inguinal (iWAT), mesenteric (mWAT) and perirenal (pWAT) WAT depots, as well as brown adipose tissue (BAT; Fig. S2C). CR also tended to decrease the absolute mass of pericardial WAT (pcWAT) in males only (Fig. S2C). To determine if these changes were proportionate to changes in overall body mass, we further analysed the mass of each adipose depot as % body mass (Fig. 2G, S3A). This showed that the significant sex-diet interaction persisted for gWAT, iWAT, mWAT and pWAT. CR did not affect % BAT mass in either sex, though % BAT mass was significantly greater in males than in females (Fig. S3A). Thus, both in absolute terms and as % body mass, CR decreases WAT mass in males but not in females.

We next analysed the masses of other tissues to determine if this sex-dependent effect of CR is unique to WAT. CR significantly decreased the absolute mass of the liver, pancreas, kidneys, gastrocnemius muscle (gastroc), heart, spleen and thymus (Fig S3B). Significant sex-diet effects were detected for the liver and kidney, with CR causing greater decreases in males than in females; however, for each of the other tissues the CR effect was similar between the sexes (Fig. S3B). In contrast to these effects on absolute mass, CR did not significantly affect the relative masses of each of these tissues, nor did sex influence the CR effect (Fig. S3C). This indicates that the absolute masses of these tissues decreased in proportion to the changes in overall body mass. One notable exception is the adrenal glands, the mass of which was increased with CR to a greater extent in males than in females (Fig. S3B-C). Together, these data show that the sex differences in CR-induced loss of body mass and fat mass are driven primarily by decreased WAT mass in males, which females robustly resist.

### Females resist adipocyte hypotrophy and lipolysis during CR

Differences in WAT mass can be driven by changes in adipocyte size and/or adipocyte number. Thus, given the marked sex differences in the effect of CR on WAT mass, we next investigated this effect at the level of adipocyte size (Fig. 2H-I, Fig. S4A-B). CR significantly decreased average adipocyte area in males but not in females, both for gWAT (Fig. 2H-I) and for mWAT (Fig. S4A-B). This suggests that females resist lipolysis during CR. Consistent with this, CR increased plasma NEFA concentrations in males but not in females (Fig. 2J). To further assess adipocyte lipolysis, we analysed phosphorylation of hormone-sensitive lipase (HSL) in iWAT. CR stimulated HSL phosphorylation in males but not in females (Fig. 2K-L, Fig. S4C). Moreover, across both diets HSL phosphorylation was lower in females whereas total HSL was increased by CR in females only (Fig. 2K-L, Fig. S4C). Together, these observations show that females resist lipolysis during CR.

### Females suppress energy expenditure and increase postprandial lipogenesis more than males during CR

We next investigated if altered energy expenditure also contributes to the sex differences in the CR response. To do so, we used indirect calorimetry to analyse mice during week 1 and week 3 of CR, corresponding to periods of weight loss and weight maintenance, respectively (Fig. 2A). This revealed that CR decreased total energy expenditure in both sexes, with greater decreases occurring during week 3 compared to week 1, and during nighttime compared to daytime (Fig. 3A-B). Notably, during week 1, CR females had lower daytime, nighttime, and total energy expenditure than CR males (Fig. 3A-B), likely explaining females’ resistance to weight loss and fat loss during the first week of CR (Fig. 2A-D). The daytime diet differences disappeared when normalised to lean body mass (Fig. S5A), suggesting that they are driven primarily by the loss of lean mass and a consequent reduction in basal metabolic rate. In contrast, CR still decreased in nighttime and total energy expenditure even when normalised to lean body mass (Fig. S5A). The relationship between lean mass and total energy expenditure (*P,* Slope) did not differ among the groups; however, the intercepts of the best-fit lines (*P,* Intercept) did differ significantly, both during week 1 (Fig. 3C) and week 3 (Fig. S5B). Thus, for a given lean body mass, CR-fed males and females had significantly lower energy expenditure than their AL-fed counterparts (Fig. 3C, S5B). In contrast to these decreases in energy expenditure, CR *increased* total and daytime physical activity in both sexes, with CR females having higher activity than CR males during week 1 (Fig. S5C). Together, these data show that CR decreases energy expenditure more in females than in males, despite increasing physical activity, and that factors beyond decreased lean mass contribute to this CR effect.

**Figure 3.**
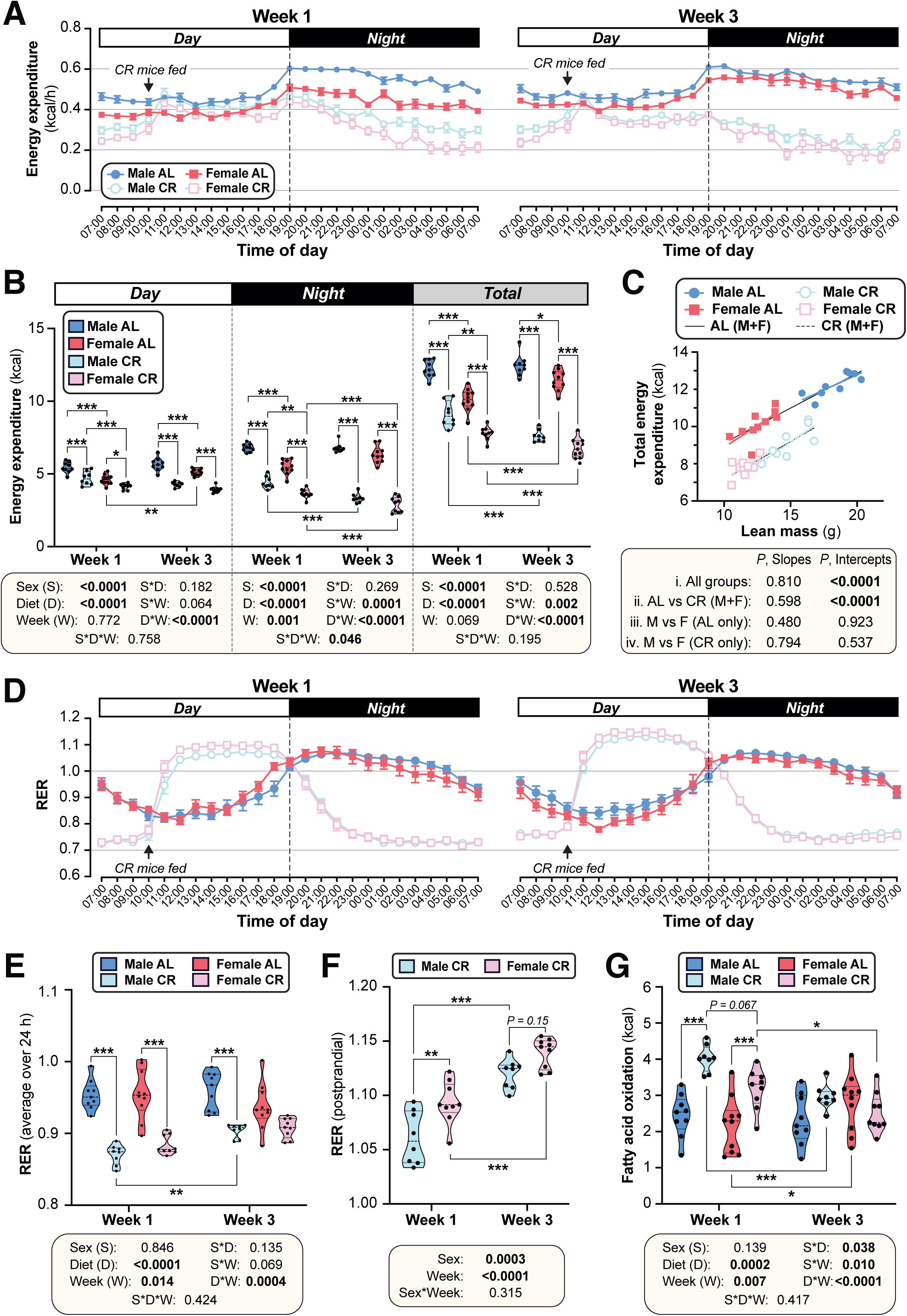
CR decreases energy expenditure and stimulates postprandial lipogenesis more in female than in male mice. Male and female mice were fed AL or CR diets, as described for Figure 2. In week 1 and week 3 after beginning AL or CR diets, mice were housed for four days in Promethion CORE System cages for indirect calorimetry. Energy expenditure (A-C) and respiratory exchange ratio (RER; D-F) was recorded every minute throughout the 4 days. **(A,D)** Average energy expenditure (A) (kcal) or RER (D) per hour over the 24 h light (Day) and dark (Night) periods, based on the average for days 2-4 of Promethion housing, for Week 1 (left) and Week 3 (right). **(B)** Overall energy expenditure (kcal) during the day, night, or day + night (Total) for Week 1 and Week 3. **(C)** Linear regression of lean mass vs total energy expenditure (kcal/24 h) during Week 1. **(E)** Average total RER (day + night) in Weeks 1 and 3. **(F)** Average RER during the postprandial period, from 12.00-17.00, for CR mice. **(G)** Absolute FA oxidation was determined based on energy expenditure and RER as described (26). Data are from 10 (*female AL*), 9 (*female CR, male AL*) or 8 (*male CR)* mice per group. In (A) and (D), data are shown as mean ± SEM. In (B) and (E-G), data are shown as violin plots overlaid with individual data points; within each time period (day, night, or total), significant effects of sex, diet, week, and interactions thereof, were determined by 3-way (B,E,G) or 2-way ANOVA (F), with *P* values shown beneath each graph. Statistically significant differences between comparable groups were further assessed by Šídák’s (B,E,G) or Tukey’s (F) multiple comparisons tests and are indicated by * (*P*<0.05), ** (*P*<0.01) or *** (*P*<0.001). For linear regression in (C), ANCOVA was used to test if the relationship between lean mass and total energy expenditure differs significantly across all of the individual diet-sex groups (*i. All mice*); between AL and CR mice, irrespective of sex (*ii. AL vs CR (M+F)*); and between males and females fed AL diet (*iii*) or CR diet (*iv*) only. ANCOVA *P* values for differences in slope and intercept are reported beneath the graph. See also Supplementary Figures 5 and 6.

We next analysed the respiratory exchange ratio (RER) to assess how diet and sex influence carbohydrate and lipid oxidation, and if these effects differ as CR progresses. In AL mice, RER peaked during the night, when most food consumption and physical activity occurs; in contrast, RER for CR mice peaked from 12.00-17.00 in the daytime, following provision of the daily ration of CR diet (Fig. 3D-E; S6A). The effects of CR on average daytime, night-time or total RER did not differ between the sexes (Fig. 3E, S6A); however, the dynamic changes in RER during these periods, and the strong influence of feeding and fasting on RER in the CR mice, make such average RER measurements difficult to interpret. Thus, we further investigated if RER in the postprandial and/or fasted states differs between CR males and females. Postprandial RER, calculated as the average RER from 12.00-17.00, exceeded 1 in both sexes (Fig. 3D, F), indicating that CR mice use dietary carbohydrates for FA synthesis (26). Notably, this postprandial RER was greater in females than in males, particularly during week 1 (Fig. 3F). Fasting RER, calculated as the average from 04.00-09.00, was below 0.8 for both sexes, indicating preferential oxidation of lipids rather than carbohydrates during this fasted state (Fig. S6B). Fasting RER was lower during week 1 than during week 3 of CR but, unlike for postprandial RER, no sex differences were observed (Fig. S6B). This suggests that relative lipid oxidation is greater in the first week of CR and is similar in males and females. However, given that females have lower absolute energy expenditure, we hypothesised that they would also have lower absolute FA oxidation (26). As shown in Fig. 3G and S6C, CR increased absolute FA oxidation in both sexes during week 1, but not during week 3. Across both timepoints there was a significant sex-diet interaction, with CR increasing FA oxidation more in males than females (Fig. 3G). Together, these data show that females have greater postprandial FA synthesis and lower absolute FA oxidation than males, particularly during the first week of CR. This highlights further mechanisms through which females maintain fat mass during CR.

### Females resist CR-induced improvements in glucose homeostasis

We next investigated if other metabolic effects of CR also differ between the sexes. We found that CR decreased blood glucose to a greater extent in males than in females (Fig. 4A). Consistent with this, oral glucose tolerance tests (OGTT) revealed that CR improved glucose tolerance in both sexes, but this effect was greater in males (Fig. 4B). These diet and sex effects were reflected by the OGTT total area under the curve (tAUC), calculated against 0 mM blood glucose; however, they were not apparent for the incremental AUC (iAUC), calculated against fasting blood glucose for each mouse (Fig. 4C). This suggests that differences in fasting glucose are the main driver of CR-induced improvements in glucose tolerance and the sex differences therein.

**Figure 4.**
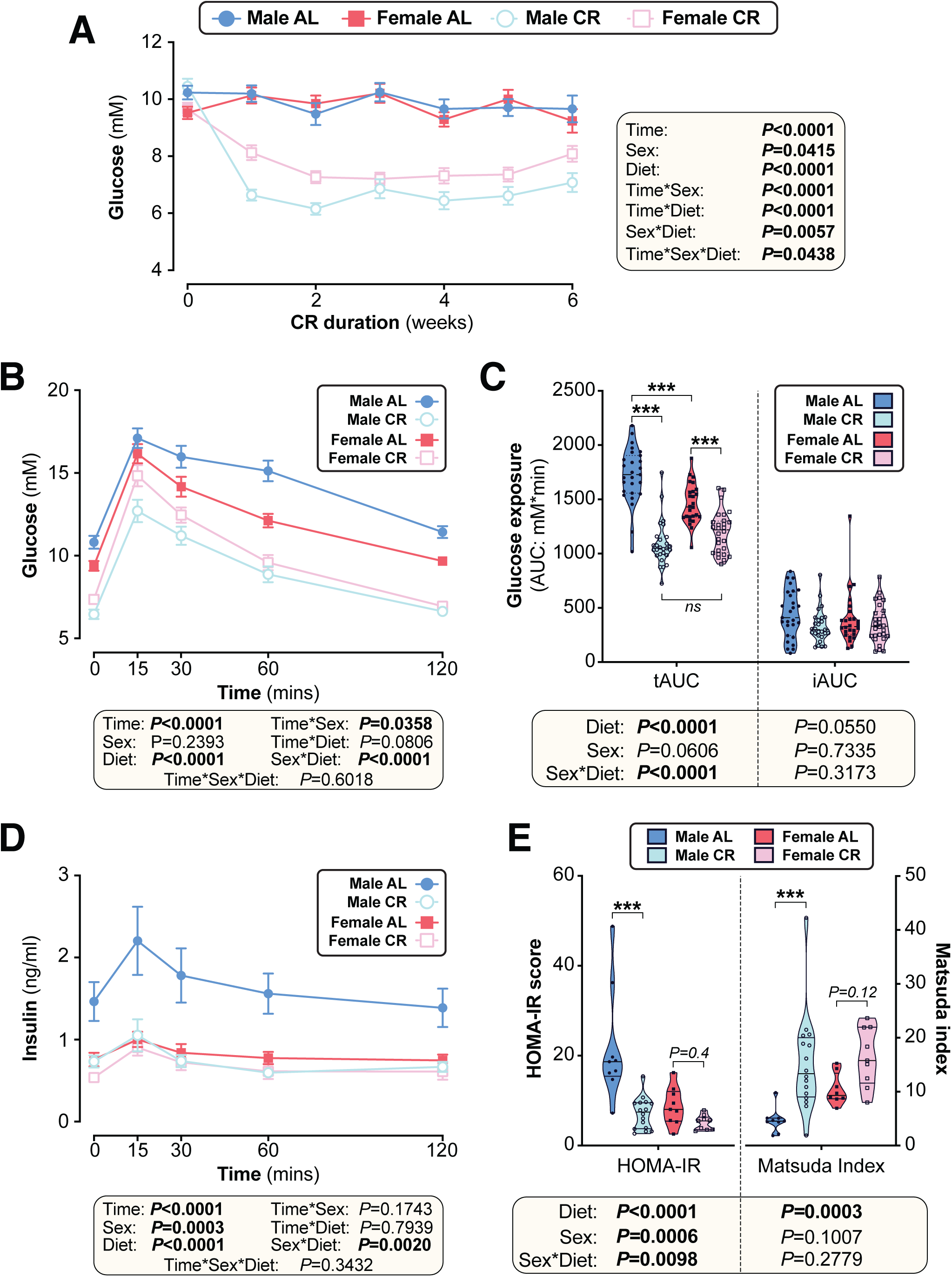
The effects of CR on glucose homeostasis differ between young male and female mice. Male and female C57BL6/NCrl mice were fed AL or CR diet from 9-15 weeks of age, as described for Figure 2. **(A)** Random-fed blood glucose was recorded each week. **(B-D)** At 13 weeks of age, mice underwent an oral glucose tolerance test (OGTT). (B) Blood glucose readings during the OGTT. (C) Area under the curve (AUC) during the OGTT was determined relative to 0 mmol/L (total AUC: tAUC) and relative to baseline (incremental AUC: iAUC). (D) Glucose-stimulated insulin secretion in mice during OGTT was assessed using an insulin ELISA. **(E)** HOMA-IR and Matsuda indices of mice calculated from glucose and insulin concentrations during the OGTT. Data in (A), (B), and (D) are presented as mean ± SEM. Data in (C) and (E) are presented as violin plots overlaid with individual data points. For each group and timepoint, the following numbers of mice were used: **(A)**: *male AL,* n=42 (Wk 0), 36 (Wk2, Wk 2), 34 (Wk 3), 29 (Wk 4), 26 (Wk 5), or 22 (Wk 6); *female AL,* n=43 (Wk 0), 28 (Wk 1), 35 (Wk 2), 31 (Wk 3), 27 (Wk 4), 26 (Wk 5), or 21 (Wk 6); *male CR,* n=44 (Wk 0), 40 (Wk 1), 38 (Wk 2), 35 (Wk 3), 31 (Wk 4), 29 (Wk 5), or 26 (Wk 6); *female CR,* n= 51 (Wk 0), 44 (Wk 1, Wk 2), 41 (Wk 4), 35 (Wk 4), 33 (Wk 5), or 27 (Wk 6). **(B)**: *male AL,* n=27 (T_0_, T_15_, T_30_, T_60_, T_120_); *female AL,* n=26 (T_15_, T_30_, T_120_), or 25 (T_0_, T_60_); *male CR,* n=27 (T_0_, T_15_, T_30_, T_60_, T_120_); *female CR,* n=28 (T_0_, T_15_, T_30_, T_60_, T_120_). **(C)**: *male AL,* n=27; *female AL,* n=26; *male CR,* n=27; *female CR,* n=28. **(D)**: *male AL,* n=9 (T_0_, T_15_, T_30_, T_60_, T_120_); *female AL,* n=11 (T_0_, T_15_), 10 (T_30_, T_60_), or 8 (T_120_); *male CR,* n=17 (T_0_), 16 (T_30_), 15 (T_60_), 13 (T_15_), or 11 (T_120_); *female CR,* n=11 (T_15_, T_30_, T_60_,), or 10 (T_0_, T_120_). **(E)**: *male AL,* n=9; *female AL,* n=9; *male CR,* n=16; *female CR,* n=9. In (A), (B), and (D), significant effects of diet, sex and/or time, and interactions thereof, were determined using a mixed-effects model. In (C) and (E), significant effects of sex, diet, and sex-diet interaction for tAUC, iAUC, HOMA-IR and Matsuda Index were assessed using 2-way ANOVA with Tukey’s multiple comparisons test. Overall *P* values for each variable, and their interactions, are shown with each graph. For (C) and (E), statistically significant differences between comparable groups are indicated by *** (*P*<0.001) or with *P* values shown. Source data are provided as a Source Data file. See also Supplementary Figure 8.

To further assess the basis for these effects on glucose tolerance, we analysed plasma insulin during the OGTT. CR decreased plasma insulin in males but not in females, largely because AL females were already relatively hypoinsulinaemic (Fig. 4D). Based on these blood glucose and insulin concentrations we calculated the homeostasis model assessment for insulin resistance (HOMA-IR) (27) and the Matsuda Index of insulin sensitivity (28). Here, a lower HOMA-IR score or a higher Matsuda index indicate increased insulin sensitivity. Across both sexes CR significantly decreased HOMA-IR and increased the Matsuda Index; however, within each sex the CR effect was significant for males only, indicating that females resist CR-induced improvements in insulin sensitivity (Fig. 4E).

### Time of CR feeding does not alter the sex differences in CR’s metabolic effects

Time of feeding can influence the metabolic effects of CR (5, 29). Therefore, to test if our feeding regime influenced the observed sex differences, we studied a cohort of mice that received their daily ration of CR diet in the evening instead of the morning (Fig. S7-8). Consistently, we found that the sex-dependent effect of CR on body mass, fat mass, lean mass and WAT mass was conserved (Fig. S7A-H). Similarly, an OGTT revealed that the diet and sex effects on glucose homeostasis persisted (Fig. S8A-B).

### Sex differences in glucose homeostasis are not explained by differential glucose uptake

The basis for the sex differences in CR-improved glucose homeostasis was next investigated. We first used positron emission tomography-computed tomography (PET/CT) with ^18^F-fluorodeoxyglucose (^18^F-FDG) to assess glucose uptake *in vivo* (Fig. 5, S9). We focused on tissues with high basal and/or insulin-stimulated glucose uptake, including the heart, brain, skeletal muscle, BAT, WAT, bone, and the bone marrow (BM) (30); the latter was also included because CR increases bone marrow adipose tissue (BMAT), another site of high basal glucose uptake (31).

**Figure 5.**
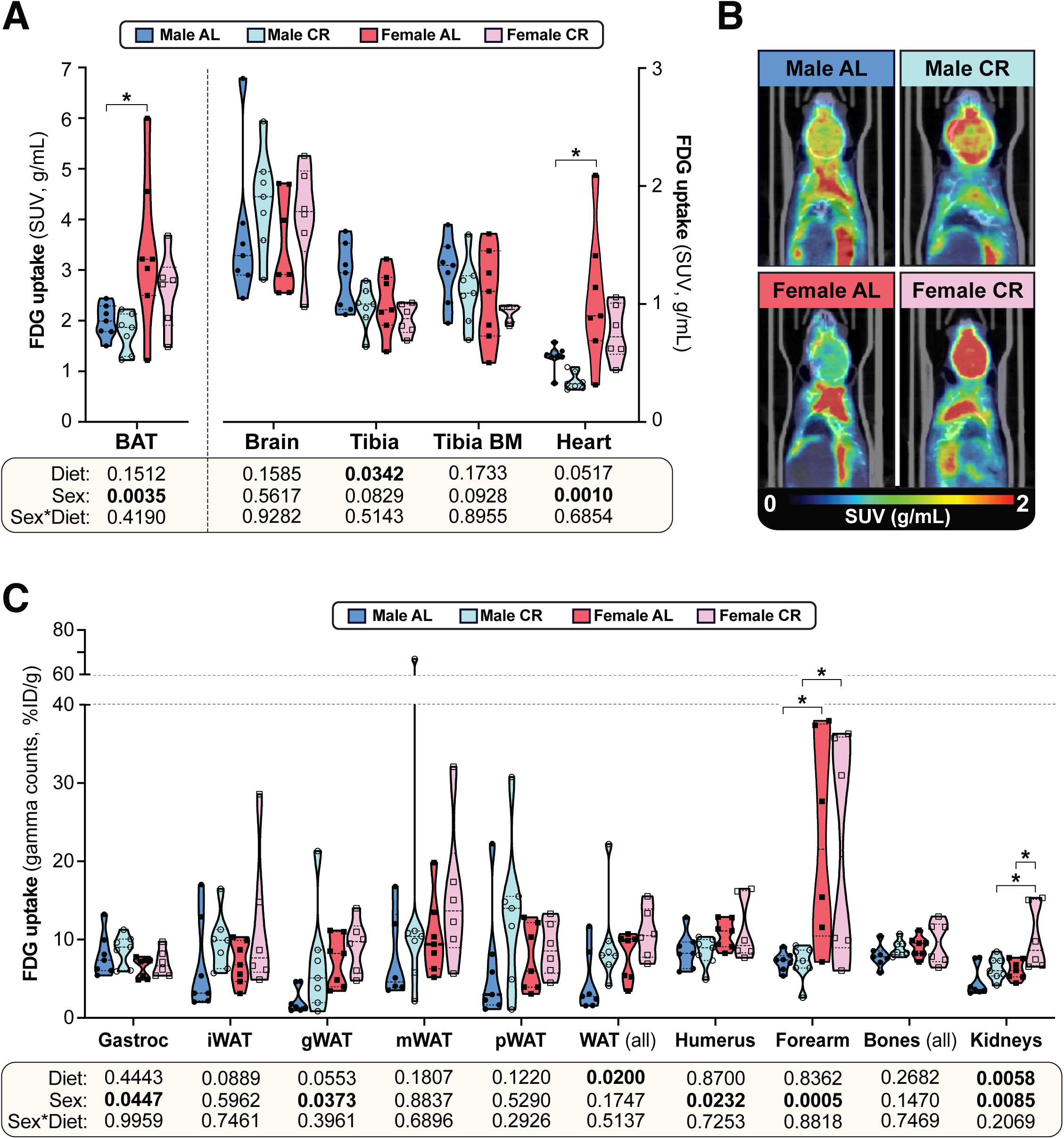
– The effects of CR on glucose uptake do not differ between male and female mice. Male and female C57BL6/NCrl mice were fed AL or CR diet from 9-15 weeks of age, as described for Figure 2. At 15 weeks of age, glucose uptake was assessed by PET/CT with ^18^F-fluorodeoxyglucose (^18^F-FDG). **(A-C)** ^18^F-FDG uptake in the indicated tissues was determined as SUVs from PET/CT scans (A) or as % injected dose per gram of tissue (%ID/g) from gamma counting of dissected whole tissues (C). **(B)** PET/CT images confirm that ^18^F-FDG uptake in interscapular BAT is greater in females than in males, regardless of diet. Data are shown as violin plots (A,C) or representative images (B) of 7 (*male AL, female AL, male CR*) or 6 (*female CR*) mice per group. For each tissue in (A) and (C), significant effects of sex, diet, and sex-diet interaction were assessed by 2-way ANOVA; overall *P* values for each variable, and their interactions, are shown beneath each graph. Significant differences between each group, as determined using Tukey’s multiple comparisons test, are indicated by * (*P* <0.05). Source data are provided as a Source Data file. See also Supplementary Figure 9.

Consistent with previous observations (31), we found that glucose uptake was highest in BAT, followed by the brain, bones, BM, the heart and skeletal muscle (Fig. 5A-B, S9A). However, CR did not stimulate glucose uptake in any of these tissues and decreased tibial glucose uptake (Fig. 5A), including in the BMAT-enriched distal tibia (Fig. S9A). Gamma counting further revealed no significant effect of CR in the gastrocnemius muscle, arm bones, liver, pancreas, spleen or thymus (Fig. 5C, S9B); however, across both sexes, CR tended to increase glucose uptake in iWAT and gWAT and this effect was significant when assessed across all WAT depots (Fig. 5C). CR also significantly increased glucose uptake in the kidneys (Fig. 5C). Several sex differences were further detected, independent of diet. Thus, females had lower glucose uptake into gastrocnemius muscle and higher glucose uptake into BAT, the heart, gWAT, the humerus, the forearm and the kidneys (Fig. 5A-C). Notably, however, sex did not significantly alter the effect of CR in any of the tissues assessed. Thus, the sexually dimorphic effects of CR on glucose homeostasis are not a result of CR increasing glucose disposal to a greater extent in males than in females.

### CR exerts sexually dimorphic effects on hepatic sphingolipid content

The lack of CR-driven differences in peripheral uptake suggested that sex differences in glucose homeostasis relate instead to effects on hepatic insulin sensitivity and glucose production. Therefore, we next tested whether altered lipid storage, implicated in sex-dependent differences in hepatic glucose production (23), might contribute to the sexually dimorphic metabolic effects of CR. Because gross hepatic triglyceride content was not affected by diet or sex (Fig. 6A), we undertook a more-detailed molecular investigation of lipid species that may underpin altered hepatic glucose regulation, focussing on sphingolipids (32, 33). Targeted lipidomics revealed that total ceramide content was significantly lower in females than in males but was not influenced by CR (Fig. 6B); however, individual ceramide species were modulated by CR in a sex-dependent manner. Thus, 22:0 and 22:1 ceramides were lower in females than in males and were decreased by CR, with this diet effect being stronger in males (Fig. 6B, S10A). Conversely, 23:0 ceramides were greater in AL females than in AL males and were increased by CR in males only (Fig. 6B); a similar pattern occurred for 14:0, 16:0, 18:0, 18:1, 20:1, 26:0 and 26:1 ceramides (Fig. S10A).

**Figure 6.**
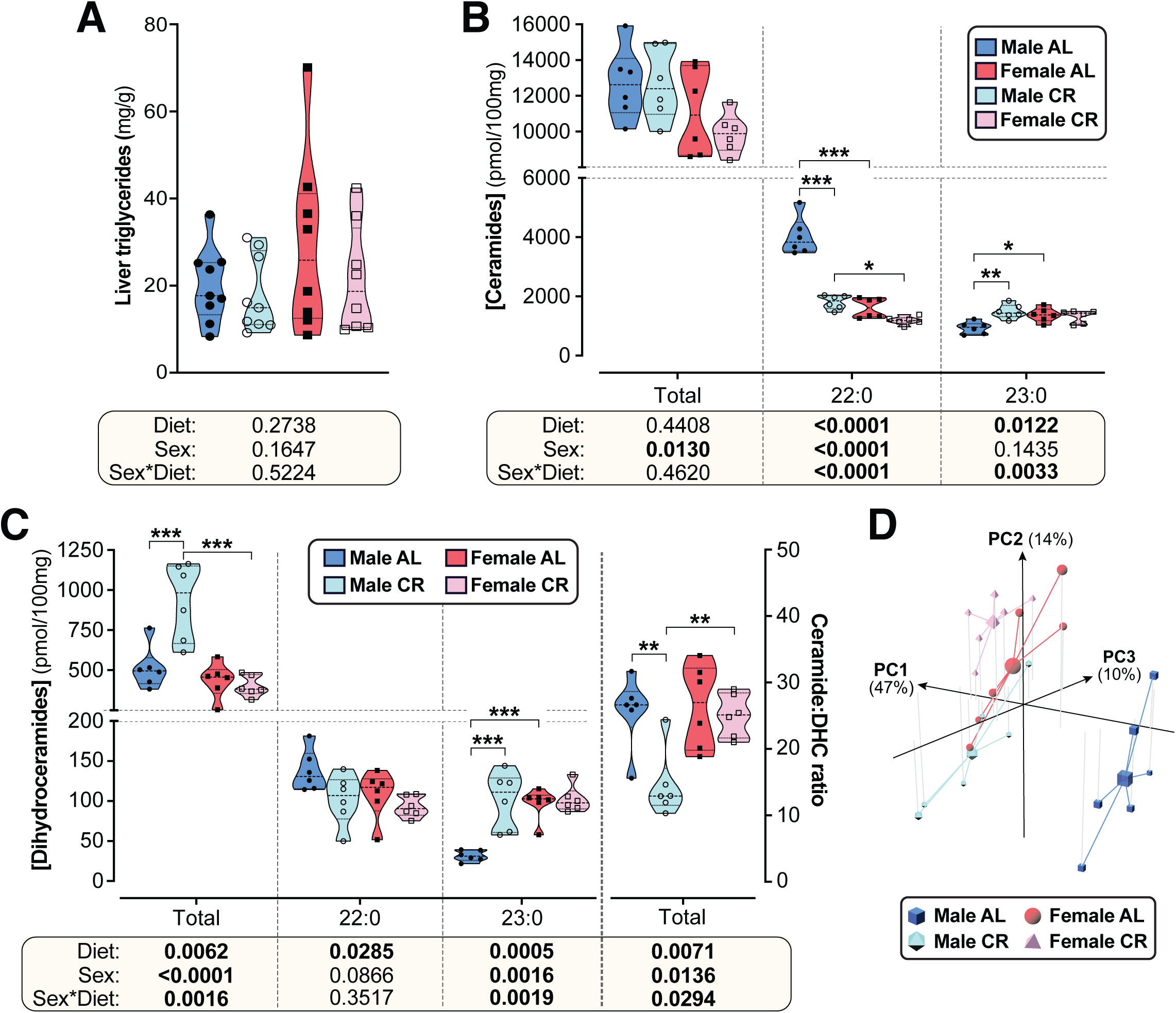
CR exerts sexually dimorphic effects on hepatic sphingolipid content. Male and female C57BL6/NCrl mice were fed AL or CR diet from 9-15 weeks of age, as described for Figure 2. Livers were sampled at necropsy (15 weeks) and used to assess triglyceride, ceramide, and dihydroceramide. **(A)** Hepatic triglyceride content. **(B,C)** LC-MS analysis of total, 22:0 and 23:0 ceramides (B), dihydroceramides (C, left side) and the ceramide:DHC ratio (C, right side). **(D)** Principal component analysis of ceramide and dihydroceramide content based on data in (B-C) and Supplementary Figure 8. PC1, PC2 and PC3 account for 47.4%, 14.1% and 10.5% of the variance, respectively. Data in (A), (B), and (C) are presented as truncated violin plots overlaid with individual data points. Data represent the following numbers of mice per group. **(A)**: *male AL,* n=9; *female AL,* n=8; *male CR,* n=9; *female CR,* n=8. **(B-D):** 6 mice per group. For (A), (B), and (C), significant effects of sex, diet, and sex-diet interaction were assessed using 2-way ANOVA; overall *P* values for each variable, and their interactions, are shown beneath each graph. Significant differences between comparable groups were assess using Tukey’s multiple comparison test and are indicated by * (*P*<0.05), ** (*P*<0.01) or *** (*P*<0.001). Source data are provided as a Source Data file. See also Supplementary Figure 10.

Ceramides are generated from dihydroceramides (DHCs) and an elevated hepatic ceramide:DHC ratio promotes metabolic dysfunction and insulin resistance (33, 34). Thus, we further analysed DHCs and the ceramide:DHC ratio as potential mediators of CR’s sexually dimorphic metabolic effects. As for total ceramides, total DHCs were lower in females than in males; however, unlike for total ceramides, CR increased total DHCs in males only (Fig. 6C). Differential effects of sex and/or diet also occurred for the specific DHC species. A common pattern was for CR to increase DHC concentrations in males only, such that these were lowest in AL males but similar in CR males, CR females and AL females; this pattern occurred for 16:0, 18:0, 18:1, 20:0, 23:0, 24:0, and 24:1 DHC species (Fig. 6C, S10B). CR increased 14:0 DHCs in both sexes, albeit more strongly in males, and decreased 22:0 DHCs across both sexes (Fig. 6C, S10B). The overall ceramide:DHC ratio was decreased by CR in males but not in females (Fig. 6C) and this also occurred for many of the individual species, including 16:0, 18:0, 20:0, 22:0, 22:1, 23:0, 24:0 and 24:1 (Fig. S10B); however, unlike for the total ceramide:DHC ratio, the ratio for each of these species was highest in the AL males and significantly lower in females, such that CR males had a similar ratio to females on either diet (Fig. S10B). Consistent with this, principal component analysis of these ceramide data identified AL males as the most distinct group, with CR males being more similar to females on either diet (Fig. 6D).

Together, these data indicate that CR suppresses ceramide synthesis to a greater extent in males than in females, highlighting a potential mechanism for CR’s sexually dimorphic effects on glucose homeostasis.

### CR exerts sexually dimorphic effects on hepatic gene expression and ketogenesis

To further investigate how hepatic function contributes to the sex differences in glucose homeostasis, we used RNA-seq to characterise the livers of AL and CR mice. Principal component analysis of this RNA-seq data identified AL males as the most transcriptionally distinct group, while CR males were more similar to AL or CR females (Fig. 7A). Volcano plots further revealed the substantial effects of sex and diet on hepatic gene expression (Fig. 7B, Fig. S11A-B): hundreds of genes were differentially expressed between males and females within each diet group (Fig. 7B, Fig. S11B) and between AL and CR mice, both within and across the two sexes (Fig. S11A-B). To assess the metabolic implications of these transcriptional differences, we first analysed the expression of transcripts that encode proteins associated with gluconeogenesis, glycolysis, or the TCA cycle. Few significant changes were detected for individual transcripts (Fig. 7B, Fig. S11A), and therefore we used gene set enrichment analysis (GSEA) to assess higher-level transcriptional changes associated with these and other relevant metabolic pathways (Fig. 7C). Gluconeogenesis-related genes were significantly increased with CR in each sex, but, against our expectations, they were more highly expressed in males than in females during CR (Fig. 7C). The greatest difference between CR males and CR females was for genes related to oxidative phosphorylation and the TCA cycle, which increased with CR in males but not in females (Fig. 7C); this was also apparent from the distribution of individual TCA-related genes in the volcano plots (Fig. 7B, Fig. S11A). Glycolysis-related genes were also increased by CR, more so in males, whereas transcripts relevant to FA β-oxidation and FA biosynthesis were increased by CR in both sexes (Fig. 7C, Fig. S11C-D). These gene expression profiles support the following model (Fig. 7D): during CR, both sexes increase FA β-oxidation to generate acetyl-CoA and this is supported by increased FA biosynthesis. Males, more than females, also increase glycolysis for acetyl-CoA production; however, males utilise the acetyl-CoA to support increased TCA cycle activity and oxidative phosphorylation, pathways that are not increased by CR in females. If so, females should have greater hepatic acetyl-CoA levels than males during CR. To test this, we measured plasma ketone concentrations, which are a biomarker for hepatic acetyl-CoA (35). Consistent with our model, plasma ketones were significantly higher in females than males, particularly during CR (Fig. 7E). Hepatic acetyl-CoA stimulates gluconeogenesis by activating pyruvate carboxylase; thus, despite transcriptional profiles suggesting increased gluconeogenesis in males (Fig. 7C), females’ higher levels of acetyl-CoA would be expected to cause greater stimulation of gluconeogenesis, thereby explaining why, compared to males, females are better at maintaining blood glucose during CR (Fig. 7D).

**Figure 7.**
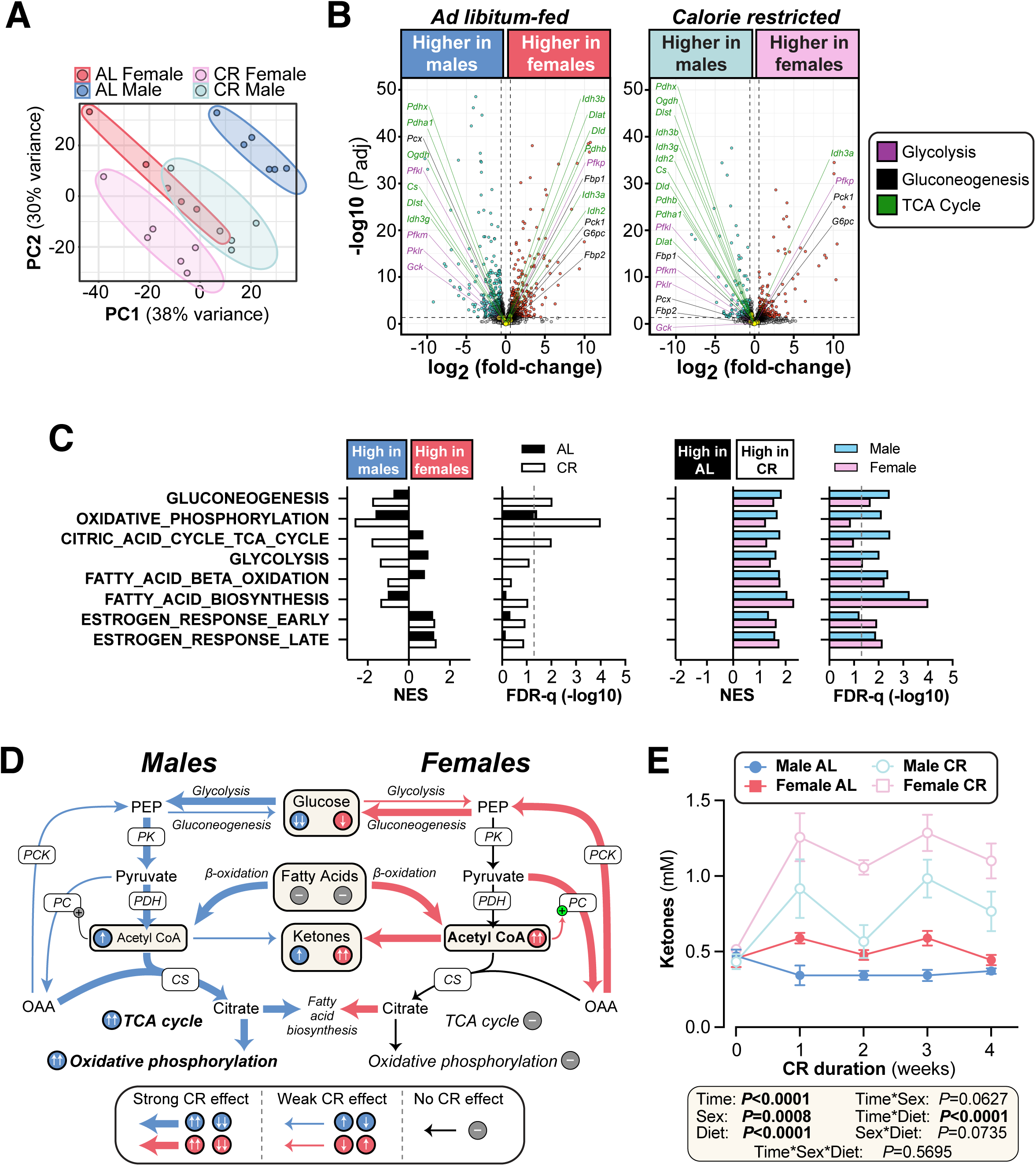
CR exerts sexually dimorphic effects on hepatic gene expression and blood ketone levels. Male and female C57BL6/NCrl mice were fed AL or CR diet from 9-13 weeks of age, as described for Figure 2. Blood ketone concentrations were assessed weekly. Livers were sampled at necropsy (13 weeks) and analysed by RNA-seq. **(A)** Principal component analysis of all samples. PC1 and PC2 account for 38% and 30% of the variance, respectively. **(B)** Volcano plots showing differentially expressed genes between AL males and AL females (left) or between CR males and CR females (right). Transcripts encoding key proteins in gluconeogenesis, glycolysis, or the TCA cycle are shown on each graph. The names of genes with positive fold-change values are shown on the right side and those with negative fold-change values are shown on the left side of the volcano plots; none of these genes had absolute fold-change > 1.5 and *P*adj < 0.05. **(C)** GSEA results for gene sets indicative of relevant pathways. Significant sex differences (within AL or CR diets) are shown on the left; significant diet differences (within males or females) are shown on the right; NES = normalised enrichment score. **(D)** Schema showing the proposed sex differences in liver metabolism during CR, based on the RNA-seq data. PK, pyruvate kinase; PDH, pyruvate dehydrogenase; CS, citrate synthase; PC, pyruvate carboxylase; PCK, phosphoenolpyruvate carboxykinase. **(E)** Blood ketone concentrations. Data represent the following numbers of mice per group: **(A-C) –** *male AL,* n=6; *female AL,* n=5; *male CR,* n=5; *female CR,* n=6. **(E) –** *male AL,* n=7; *female AL,* n=9; *male CR,* n=6; *female CR,* n=7. For (A) and (B), R script are provided as a Source Data file. For (C), gene set collection files are prepared with the 8 gene sets shown for GSEA. For (E), significant effects of sex, diet, and sex-diet interaction were assessed using 3-way ANOVA; overall *P* values for each variable, and their interactions, are shown beneath each graph. Source data are provided as a Source Data file. See also Supplementary Figure 11.

### Sex differences in CR’s metabolic effects are absent when CR is initiated in aged mice

Another intriguing finding from our RNA-seq data is that CR significantly stimulates oestrogen response pathways (Fig. 7C). Thus, one possibility is that young females resist many of CR’s metabolic effects because their endogenous oestrogens cause them to remain relatively metabolically healthy, even on an AL diet (36, 37). To begin testing if endogenous oestrogens influence the CR response, we next investigated the effects of CR initiated in 18-month-old mice, when females are anoestrous (38). Unlike in young mice, CR decreased body mass and fat mass to a similar extent in aged males and females, while the decreases in lean mass were greater in aged females (Fig. 8A-C). Consistent with this, CR’s effects on the masses of individual WAT depots did not differ between the sexes (Fig. 8D, S12A). Among non-adipose tissues CR significantly decreased pancreas mass only in aged females, but there were no sex differences in the effects of CR on the masses of other tissues (Fig. S12B-C). Thus, unlike in young mice, aged males and females show a similar effect of CR on overall adiposity and this is driven predominantly by loss of WAT.

**Figure 8.**
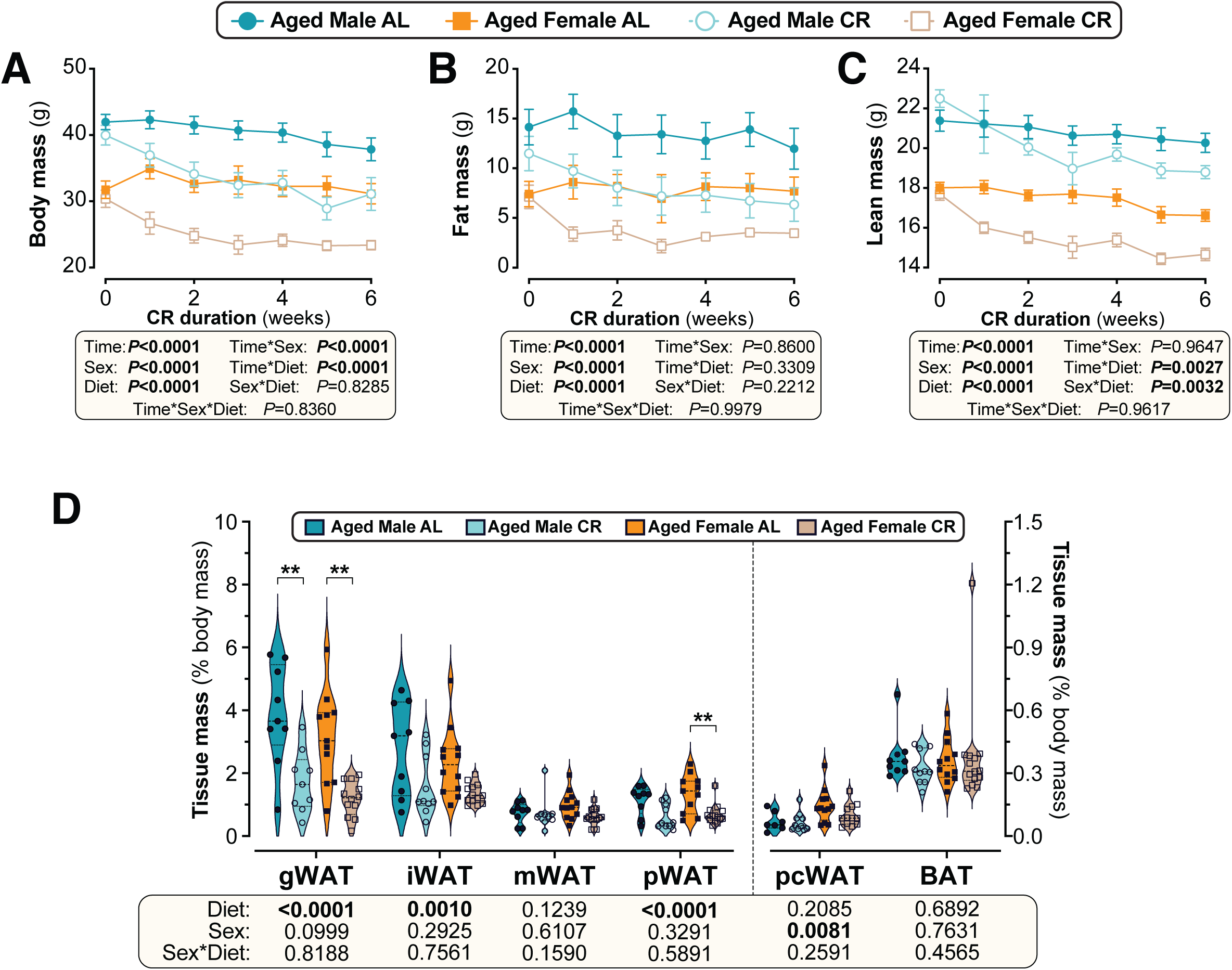
Sex differences in CR-induced weight loss and fat loss are absent when CR is initiated in aged mice. Male and female C57BL6/NCrl mice were single housed and fed AL or a CR diet (70% of daily AL intake) from 78-84 weeks of age. **(A-F)** Each week mice were weighed (A,B) and body composition was determined by TD-NMR (C-F). Body mass, fat mass and lean mass are shown as absolute masses (A,C,E) or fold-change relative to baseline (B,D,F). **(G)** Masses of gWAT, iWAT, mWAT, pWAT, pcWAT and BAT were recorded at necropsy and are shown as % body mass. Data in (A-F) are shown as mean ±SEM. Data in (G) are shown as violin plots overlaid with individual data points. For each group and timepoint, the following numbers of mice were used: **(A,B)**: *male AL*, n=9 (Wk 0, Wk 2, Wk3, Wk 4), 8 (Wk 1), or 6 (Wk 5, Wk 6); *female AL,* n=13 (Wk 0, Wk 2), 12 (Wk 4, Wk 5), 11 (Wk 6), 9 (Wk 1, or 8 (Wk 3); *male CR,* n=10 (Wk 0, Wk 2-4), 9 (Wk 1), 7 (Wk 5), or 6 (Wk 6); *female CR,* n=14 (Wk 0, Wk 4, Wk 5), 13 (Wk 2, Wk 6), 10 (Wk 3), or 9 (Wk 1). **(C-F)**: *male AL,* n=9 (Wk 0, Wk 2), 8 (Wk 3, Wk 4, Wk 6), or 7 (Wk 1, Wk 5); *female AL,* n=13 (Wk 0, Wk 2), 9 (Wk 4, Wk 6), 8 (Wk 5), 6 (Wk 1), or 5 (Wk 3); *male CR,* n=10 (Wk 0, Wk 2), 9 (Wk3-6), or 8 (Wk 1); *female CR,* n=14 (Wk 1, Wk 2), 11 (Wk 4-6), 7 (Wk 3), or 6 (Wk 1). **(G)**: *male AL,* n=9; *female AL,* n=12 (iWAT, mWAT) or 11 (gWAT, pWAT); *male CR,* n=10 (iWAT, pWAT), or 9 (gWAT, mWAT); *female CR,* n=14 (mWAT, pWAT), 13 (iWAT), or 12 (gWAT). For (A-F), significant effects of diet, sex and/or time, and interactions thereof, were determined by 3-way ANOVA or a mixed-effects model. For (G), significant effects of sex, diet, and sex-diet interaction were assessed using 2-way ANOVA with Tukey’s multiple comparisons test. Overall *P* values for each variable, and their interactions, are shown beneath each graph. For (G), significant differences between comparable groups are indicated by ** (*P*<0.01) or *** (*P*<0.001). Source data are provided as a Source Data file. See also Supplementary Figures 12 and 14.

We then investigated CR’s effects on glucose homeostasis and insulin sensitivity. As shown in Figure 9A, in aged mice the ability of CR to decrease fasting blood glucose was not significantly influenced by sex. CR improved glucose tolerance to a similar extent in aged males and females, with tAUC but not iAUC being significantly decreased with CR (Fig. 9B-C). The latter indicates that, as for young mice, CR improves glucose tolerance in aged mice primarily by decreasing fasting glucose. Despite no sex differences in glucose tolerance, both sex and diet decreased glucose-stimulated insulin secretion, with this being greatest in aged AL males and decreased by CR more so in males than in females (Fig. 9D). Interestingly, a sex-diet effect was observed for HOMA-IR but not for the Matsuda Index (Fig. 9E); this is similar to the effects observed in young mice (Fig. 4E) and suggests that in aged mice there remain some sex differences in the relationship between plasma insulin and gluconeogenesis in the fasted state. However, overall, these studies demonstrate that sex differences in the metabolic effects of CR are highly age dependent, with CR being equally effective in aged males and females whereas young females resist these aspects of the CR response.

**Figure 9.**
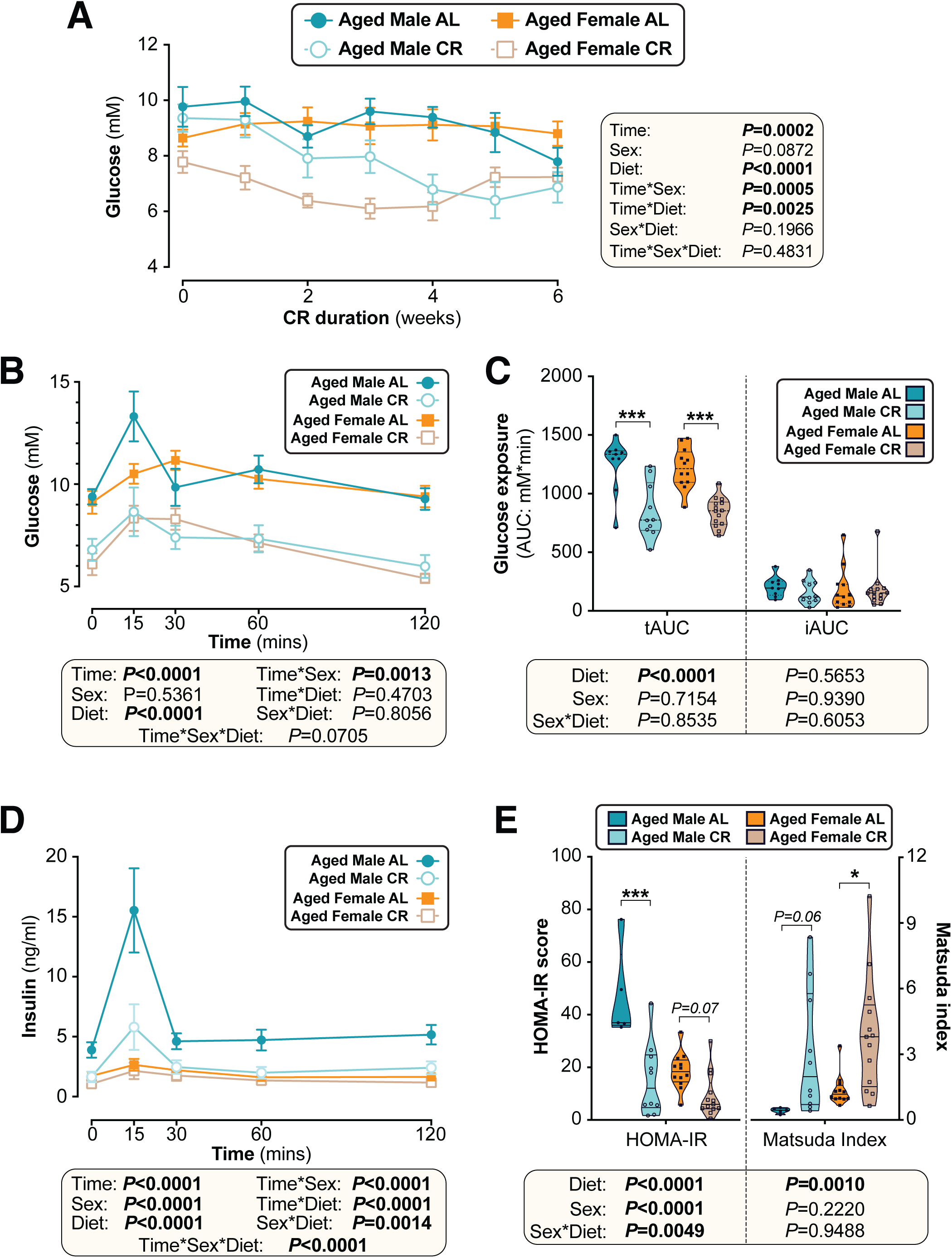
Sex differences in CR’s effects on glucose homeostasis are largely absent when CR is initiated in aged mice. Male and female C57BL6/NCrl mice were fed AL or CR diet from 78-84 weeks of age, as described for Figure 7. **(A)** Random-fed blood glucose was recorded each week. **(B-D)** At 82 weeks of age mice underwent OGTTs. Blood glucose was recorded at each timepoint (B) and the tAUC and iAUC was determined (C). (D) Glucose-stimulated insulin secretion in mice during OGTT was assessed using an insulin ELISA. **(E)** HOMA-IR and Matsuda indices of mice calculated from glucose and insulin concentrations during the OGTT. Data in (A), (B), and (D) are presented as mean ± SEM. Data in (C) and (E) are presented as violin plots overlaid with individual data points. For each group and timepoint the following numbers of mice were used: **(A)**: *male AL,* n=9 (Wk 0, Wk 2-4, Wk 6), 8 (Wk 1), or 5 (Wk 5); *female AL,* n=12 (Wk 0, Wk2, Wk4), 8 (Wk 1, Wk 3), or 11 (Wk 5, Wk 6); *male CR,* n=10 (Wk 0, Wk 2-4, Wk 6), 9 (Wk 1), or 7 (Wk 5); *female CR,* n= 14 (Wk 0, Wk 2, Wk 4-6), 9 (Wk 1) or 10 (Wk 3). **(B-E)**: *male AL,* n=9 (B-C) or 5 (D-E); *female AL,* n=12; *male CR,* n=10; *female CR,* n=14. In (A), (B), and (D), significant effects of diet, sex and/or time, and interactions thereof, were determined using a mixed-effects model or 3-way ANOVA. In (C) and (E), significant effects of sex, diet, and sex-diet interaction for tAUC, iAUC, HOMA-IR and Matsuda Index were assessed using 2-way ANOVA with Tukey’s multiple comparisons test. Overall *P* values for each variable, and their interactions, are shown with each graph. For (C) and (E), statistically significant differences between comparable groups are indicated by * (*P*<0.05) or *** (*P*<0.001), or with *P* values shown. Source data are provided as a Source Data file.

### In humans CR-induced fat loss is sex- and age-dependent

Finally, we investigated if the metabolic effects of CR in humans are also sex- and age-dependent. To do so we retrospectively analysed data from forty-two overweight and obese men and women who participated in a weight loss study involving a 4-week CR intervention. As in young mice, body mass and fat-free (lean) mass decreased more so in males than in females, with males also showing a trend for greater fold-decreases in total fat mass (Fig. 10A-F). When adjusted for age (Fig. 10G), fold-change in body mass differed significantly between males and females (*P,* Intercept = 0.025). This demonstrates that overall weight loss is greater in males than females, independent of age. In contrast, the decreases in fat mass and fat-free mass were sex- and age-dependent. Thus, fat loss in younger individuals was greater in males than females but, with age, fat loss increased in females but diminished in males (Fig. 10H) (*P,* Slope = 0.007). Conversely, the loss of fat-free mass in younger individuals was greater in females than in males and this relationship reversed in older individuals (Fig. 10I). As for fat mass, the relationship between age and loss of fat-free mass differed significantly between the sexes (*P,* Slope = 0.0004). In contrast, the sex differences in loss of fat mass or fat-free mass were unrelated to fat mass or BMI at baseline (Fig. S13).

**Figure 10.**
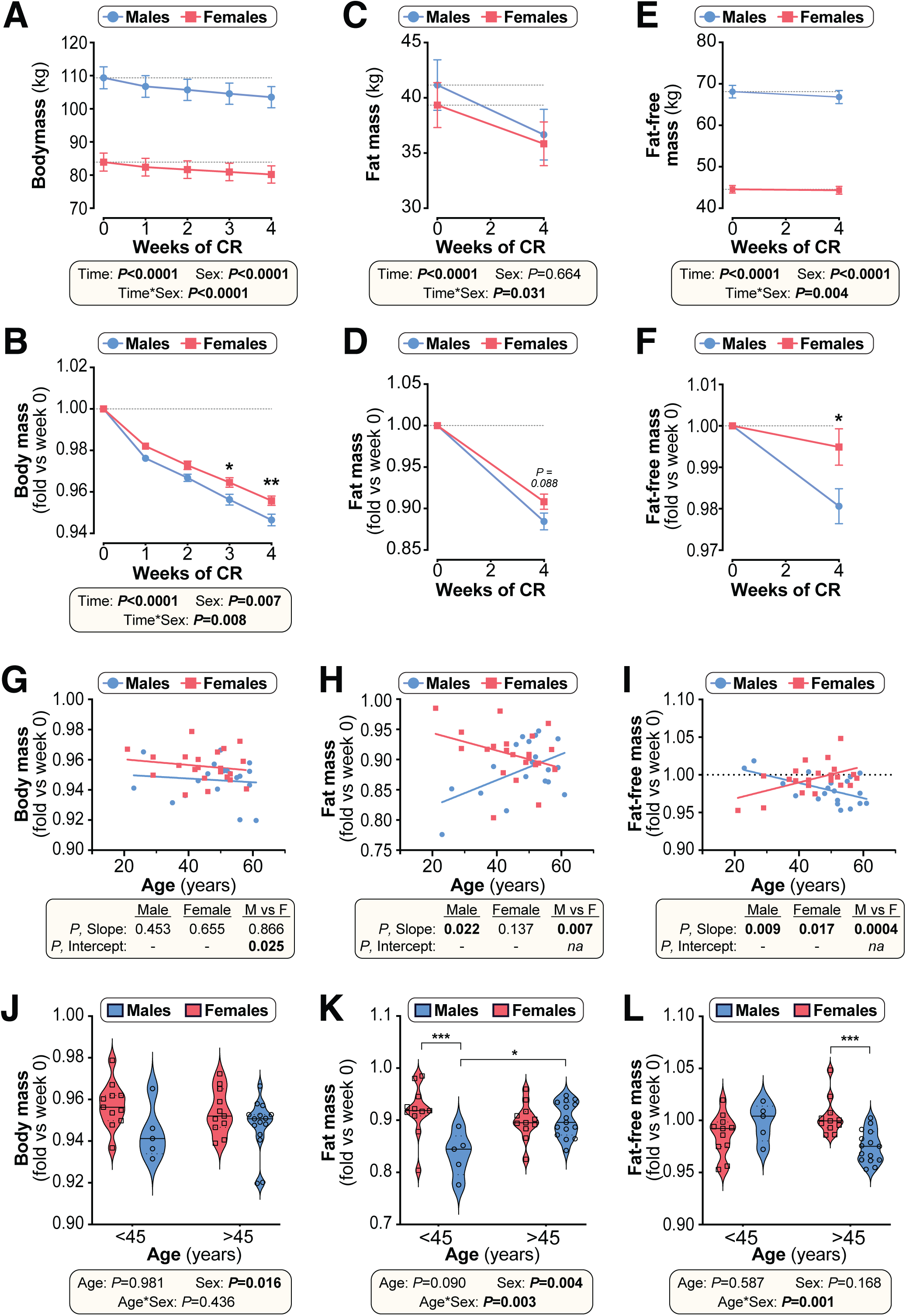
Effects of CR on body mass and body composition in humans are sex- and age-dependent. Twenty male and twenty-two female volunteers participated in a weight loss study involving a 4-week dietary intervention. **(A-F)** Body mass was recorded each week (A,B). Fat mass (FM) and fat-free mass (FFM) were measured by air-displacement whole-body plethysmography at weeks 0 and 4 (C-F). Body mass, fat mass and fat-free mass are shown as absolute masses (A,C,E) or fold-change relative to baseline (B,D,F). Data are presented as mean ± SEM. For (A-C) and (E), significant effects of time, sex, and time*sex interaction were assessed using 2-way ANOVA. In (B), (D) and (F), significant differences between males and females at each time point were determined by Šídák’s multiple comparisons test (B) or unpaired T-test (D,F) and are indicated by * (*P*<0.05) or ** (*P*<0.01). **(G-I)** Simple linear regression of age vs fold-change (*week 4 vs week 0)* in body mass (G), fat mass (H) and fat-free mass (I). For each sex, significant associations between age and outcome (fold-change) are indicated beneath each graph as ‘*P,* Slope’. ANCOVA was further used to test if the age-outcome relationship differs significantly between males and females. ANCOVA results are reported beneath each graph as ‘*P,* Slope’ and ‘*P,* Intercept’ for males vs females (M vs F). In (G), similar slopes but different intercepts show that sex significantly influences weight loss, but the influence of age does not differ between the sexes. In (H,I) the slopes differ significantly, indicating that the age-outcome relationship differs between the sexes. **(J-L)** Fold-change (week 4 vs week 0) in body mass (J), fat mass (K) and fat-free mass (L) for males vs females separated into younger (<45 years) and older (>45 years) groups. Data are presented as violin plots overlaid with individual data points. Significant effects of age, sex, and age*sex interaction were assessed using 2-way ANOVA with Tukey’s multiple comparisons test. Overall *P* values for each variable, and their interactions, are shown beneath each graph. Significant differences between comparable groups are indicated by * (*P*<0.05) or *** (*P*<0.001). Source data are provided as a Source Data file. See also Table 3 and Supplementary Figures 13-15.

These age-sex interactions were particularly apparent when the cohort was divided into younger and older groups (<45 vs >45 years of age), selected based on the age when females’ oestrogen levels begin to decline (39). Total weight loss was greater in males than females regardless of age (Fig. 10J), but only in the younger age group did males show greater fat loss than females (Fig. 10K). Conversely, males’ greater loss of fat-free mass occurred only in the older age group (Fig. 10L). Notably, a similar pattern of age-dependent fat loss was apparent for CR in young vs aged mice (Fig. S14). Thus, our data demonstrate that the ability of CR to decrease body mass, fat mass and lean mass is sex- and age-dependent in both mice and humans.

## DISCUSSION

Herein we aimed to determine the extent and basis of sex differences in the metabolic effects of CR. We reveal that 96.6% (mouse) and 95.7% (human) of published CR research overlooks potential sex differences, with most mouse studies using males only whereas most human studies combine data from males and females. We show that, in young mice, CR decreases fat mass and improves glucose homeostasis to a greater extent in males than in females, whereas these sex differences are blunted or absent when CR is initiated in old mice. Young CR females likely resist weight loss and fat loss by having lower energy expenditure, lipolysis and FA oxidation, and greater postprandial lipogenesis, than young CR males, The effects of CR on hepatic gene expression and sphingolipid content also differ between young male and female mice, highlighting the liver as a key mediator of CR’s sexually dimorphic metabolic effects. Finally, we reveal that humans also exhibit age-dependent sex differences in the CR response, with younger age groups showing greater fat loss in males than females whereas, with increasing age, CR-induced fat loss becomes similar between the sexes. Together, these data identify sex and age as factors that have a major influence on the metabolic health outcomes of CR.

### Age-dependent sex differences in CR’s metabolic benefits

Our observation that young females resist overall weight loss during CR is consistent with some previous findings in mice (21, 22, 40), monkeys (3), and humans (41–46). Young females’ resistance to CR-induced fat loss has also been reported in mice (21, 22, 47–49) and humans (20, 43, 50–52); however, fewer studies have directly assessed sex differences in the effect of CR on glucose homeostasis and insulin sensitivity. Many studies have included males and females but did not sex-stratify their analyses, thereby overlooking the potential impact of sex on these important metabolic outcomes (13, 53). Nevertheless, CR in monkeys and humans has been found to improve glucose homeostasis and insulin sensitivity more so in males than in females (3, 44), consistent with our findings in young mice.

Our observations fundamentally extend this previous research by demonstrating, for the first time, that these sex differences are age dependent. The novelty of this finding highlights that relatively few studies have investigated CR when initiated in aged subjects. In 2004, Dhahbi *et al* showed that CR’s health benefits persist even when initiated in aged mice (54), and subsequent research has compared the metabolic effects of CR when implemented in young vs aged animals (55, 56). Unfortunately, these studies did not include females, an oversight that has persisted in the CR field (Fig. 1, S1). Other notable research has comprehensively compared aged males and females, but only in the context of lifelong CR that was initiated in younger mice (57). Two recent studies have also highlighted age-dependent sex differences in the effects of methionine restriction or time-restricted feeding, but these dietary interventions were limited to young vs middle-aged mice (36, 37). The most relevant previous research is perhaps from Brandhorst *et al,* who observed significant visceral fat loss in female mice fed a ‘fasting-mimicking diet’ from 16-20.5 months of age (58). While this is consistent with our observations in aged females (Fig. 8, S12), it is limited by having not assessed glucose homeostasis nor including male counterparts for direct comparison. Thus, our study is the first to compare CR when initiated in young or old males and females and to thereby identify age-dependent sex differences in CR’s metabolic effects.

Our findings have several implications for interpreting previous CR research. Although most studies show that young females resist overall weight loss and fat loss during CR, such sex differences have not been universally observed. For example, in young female rats some studies report that CR decreases body mass and/or fat mass to a similar extent in both sexes (59–61), while others find fat loss to be even greater in females than in males (62, 63). CR-induced fat loss has also been noted in young female mice (64–67), albeit in studies that lack males for direct comparison. Several human studies also show no differences in total fat loss with CR, even in younger women who are unlikely to be menopausal (44, 46, 68). What are the reasons for these inconsistent results, and can they further inform the basis for sex differences in the CR response?

Considering our present findings, one possibility is that, even in younger subjects, age differences might have influenced the outcomes of these CR studies. Indeed, both Janssen *et al* and Das *et al* studied women who, while pre-menopausal, were significantly younger than their male counterparts (46, 68). Given that males’ fat loss decreases as they get older (Figs. 10H,K), this disparity could have undermined the ability of these studies to detect sex differences in fat loss. A related, broader issue is that much human CR research has studied females who are over 45 years old (41, 43, 44, 50, 52, 69–71). This is important because it is the age at which oestrogen levels typically begin to decline (39) and after which we demonstrate no sex differences in fat loss (Fig. 10K). Thus, our findings underscore the importance of properly age-matching males and females, and ideally confirming menopausal status, if the response to CR is to be reliably assessed.

While age is clearly an important variable, many animal studies still find no resistance to fat loss in young females (64–67), even when appropriately age-matched to male counterparts (59–63). Therefore, other factors must also be influencing CR’s sexually dimorphic effects. One possibility is that males show a greater CR response because they are usually heavier and fatter than females at the onset of CR, and thus have more fat available to lose. Consistent with this ‘baseline adiposity’ hypothesis, during CR in rats Hill *et al* found total fat loss to be greater in young females than young males and, unlike in mice, pre-CR adiposity was greater in the females (62, 63). Moreover, we show that sex differences in baseline adiposity become less pronounced with ageing, such that % fat mass is similar in aged male and female mice (Fig. 8D). Therefore, this hypothesis could explain why CR decreases fat mass in the fatter, aged females, whereas young, leaner females resist this effect. However, three key points argue against this hypothesis. Firstly, we show that young females’ resistance to fat loss still occurs in obese and overweight humans, among whom % fat mass is greater in females than in males, irrespective of age (Fig. S15). This is consistent with numerous previous studies of overweight or obese humans, in which CR causes more weight loss and/or fat loss in males than females (20, 41–45, 50, 52, 69). Secondly, in our human study we show that the sex differences in loss of fat or fat-free mass are unrelated to baseline adiposity or BMI (Fig. S13). Finally, in young, non-obese humans, sex differences in fat loss persist even after controlling for variation in baseline fat and body mass (46). Thus, sex differences in the CR response are a not simply a consequence of baseline differences in these parameters.

A third possibility is that young females’ resistance to weight and fat loss can be overcome if the extent and/or duration of CR is great enough. Our CR protocol involves a 30% decrease in daily caloric intake over six weeks. In contrast, those studies reporting fat loss in young females typically used CR of a greater extent and/or duration. For example, Hill *et al* studied 60% CR for 11 weeks (62, 63) while Valle *et al* used 40% CR for 14 weeks (60, 61). Others have tested CR of a similar extent to our studies (20-30%) but for longer durations, ranging from ten weeks to six months (59, 64, 66). It is now well established that the CR response is dose and duration dependent (5, 22, 48, 57, 72, 73), with the former having been systematically investigated in male mice (74–77). Thus, a key goal for future studies will be to determine how sex and age influence these dose- and duration-dependent effects.

A final important consideration is the distinction between total versus regional adiposity. Indeed, the CALERIE trial finds no sex differences in the loss of total fat mass but does show that, compared to men, women resist loss of trunk fat during CR (46). This is consistent with our data showing that fat loss in young male mice is greatest for the visceral depots (Figs. 2G, S2C, S3A, S7G), and with previous observations for CR in obese humans (20, 50, 52, 69). Some CR studies may therefore have failed to detect sex differences because they measured only total adiposity, missing out on possible site-specific differences (44, 78). Thus, any studies seeking to fully understand sex differences in CR’s metabolic effects, whether in preclinical models or in humans, should always aim to assess adiposity on both whole-body and regional levels.

### Basis for young females’ resistance to weight loss and fat loss

Our studies also shed light on the mechanistic basis for these sex differences. Previous murine CR studies show that females decrease total energy expenditure more than males, possibly resulting from greater suppression of BAT activity (21, 60, 61). Consistent with this, we find that CR suppresses energy expenditure more in young females than in young males, at least during the first week of CR (Fig. 3A-B). In contrast, CR males and females have similar energy expenditure at week 3 of CR (Fig. 3A-B), similar to our previous findings at week 4 of CR (79). Consistent with this, BAT activity is similar between AL and CR mice after 6 weeks of CR (Fig. 5). Therefore, if using indirect calorimetry to understand the CR response, it is important to ensure that the calorimetry analyses coincide with timepoints at which the relevant CR responses occur.

Another advance over our previous calorimetry studies (79) is that, by using the higher-sensitivity Promethion system, we were able to measure RER and thereby identify sex differences in lipid metabolism. We find that, compared to CR males, CR females have greater postprandial lipogenesis but lower FA oxidation and lipolysis. This confirms previous observations in male or female mice only (26, 65) but extends these earlier findings by demonstrating key sex differences in these effects. It is also like the situation in humans, in which CR enhances WAT lipolysis in both sexes and females resist this effect (43). Together, these differences in lipid metabolism likely explain why young females resist fat loss during CR. Whether ageing alters these sex differences in lipid metabolism remains to be determined.

A final consideration is how time of CR feeding influences energy homeostasis. In our morning-fed mice, CR decreases energy expenditure more at nighttime than in the day. In contrast, Dionne *et al* found that in nighttime-fed mice CR decreases energy expenditure in the day but not at night (48). However, both we and Dionne *et al* show that CR decreases 24-hour energy expenditure, demonstrating that this overall CR effect occurs irrespective of time of feeding.

### Importance of visceral fat loss for CR’s metabolic effects

These effects on fat mass and lipid metabolism likely contribute to young females’ resistance to improved glucose homeostasis during CR. Human studies have shown that inefficient WAT lipolysis is associated with impaired glucose metabolism in women (80) and that, during CR, decreased fat mass is linked to the improvements in insulin sensitivity (13). Similarly, studies of CR or lipectomy in animal models suggest that visceral fat loss contributes to CR’s broader metabolic benefits (7, 10–13). However, no previous research has investigated if this relationship between fat loss and insulin sensitisation differs between the sexes. Our work therefore supports and extends these previous studies by demonstrating that CR-induced fat loss coincides with improved glucose homeostasis and insulin sensitivity in males and aged females, whereas young females’ resist these metabolic effects.

### The liver as a key driver of sex differences in glucose tolerance

Several of our findings highlight the importance of the liver as a key mediator of CR’s sexually dimorphic metabolic effects. Firstly, our PET/CT data show that CR improves glucose tolerance without any marked increases in peripheral glucose uptake (Fig. 5). This includes the BMAT-rich distal tibia, suggesting that, although BMAT has high basal glucose uptake (31), increased BMAT glucose uptake is not required for improved glucose tolerance during CR. Moreover, although CR does increase glucose uptake in WAT and the kidneys, this effect does not differ between males and females and therefore is unlikely to explain the sex differences in glucose tolerance (Fig. 5B). Together our data are consistent with human studies showing that, in the fasted state, CR does not affect overall glucose disposal (81, 82); however, CR enhances insulin-stimulated glucose uptake (83) and it is possible that this effect differs between the sexes. Our second relevant finding is that, during an OGTT, CR decreases the tAUC but not the iAUC, highlighting decreases in fasting glucose, rather than insulin-stimulated glucose disposal, as the main driver of the improvements in glucose tolerance. Thirdly, we show that CR’s effects on hepatic gene expression differ between young male and female mice (Fig. 7, S11). These transcriptional differences suggest that, during CR, both sexes increase hepatic FA oxidation, but only males increase TCA cycle activity and oxidative phosphorylation (Fig. 7B-C). Strikingly, females have higher plasma ketone concentrations, a marker of hepatic acetyl-CoA content (35). Based on these data, our model is that CR males utilise hepatic acetyl-CoA to support the TCA cycle and ATP generation via oxidative phosphorylation; in contrast, in CR females the hepatic acetyl-CoA accumulates, thereby stimulating pyruvate carboxylase and promoting gluconeogenesis (Fig. 7D). Consequently, females are better at resisting hypoglycaemia during CR. As discussed below, future studies using metabolic tracers or related approaches will be important to further test these conclusions.

How is loss of visceral fat linked with these hepatic effects? Visceral adiposity is strongly associated with increased hepatic fat storage (84), and decreased liver fat coincides with improved insulin sensitivity during CR, both in rats and humans (85, 86). Moreover, young female mice resist the decreases in hepatic triglycerides following short-term fasting (23) and prolonged time-restricted feeding (40). Therefore, we were surprised to find that CR does not affect hepatic triglyceride content in young mice of either sex (Fig. 6A). This may be because our young AL controls are lean and healthy, and therefore already have livers that are not excessively fatty. In contrast, many of the previous studies investigated CR in animals or humans that are older and/or with type 2 diabetes (40, 85, 86), thereby increasing the baseline/control hepatic fat content. This would not explain why our findings differ to those of Della Torre et al, who studied fasting in young, healthy mice (23); however, it is perhaps unsurprising that our 6-week CR elicits different effects to those induced during shorter-term fasting.

Our above observations are important because they show that CR can exert metabolic benefits without decreasing hepatic triglyceride content. In contrast, we find that effects on sphingolipids may play a role, with CR altering hepatic sphingolipid content more so in young males than in young females. Here, the effect of diet and/or sex depends on the sphingolipid species concerned (Fig. 6; S10). This is reminiscent of the findings of Norheim *et al*, who report sex differences in hepatic ceramide content in mice fed obesogenic diets (87). They further showed that the relationship between ceramides, body fat and insulin sensitivity also differs between males and females, focussing on 16:0 ceramide (C16:0). Indeed, ceramides’ influence on insulin sensitivity depends on the ceramide species. For example, hepatic increases in C16:0 have been implicated with the development of obesity-associated insulin resistance (33, 88–90). This is seemingly at odds with our finding that CR increases hepatic C16:0 in young males but not young females (Fig. S10). Thus, unlike in obesity, increased hepatic C16:0 during CR appears to be associated with improvements in insulin sensitivity.

The rate of conversion of ceramides to DHCs may be a root cause of insulin resistance (91, 92). This underscores the concept that the ratio between each ceramide and DHC species may influence metabolic function more than the absolute values of either (34). More research is needed to determine how specific ceramide and DHC species, and the ratios thereof, impact sex differences in the health benefits of CR. Despite these remaining questions, our work extends previous observations by revealing that CR’s hepatic effects are sexually dimorphic and that this contributes to the sex differences in glucose tolerance and insulin sensitivity.

### Oestrogen as a modulator of the CR response

In many cases young females’ resistance to the metabolic effects of CR is associated with them already being relatively metabolically healthy on an AL diet. For example, young AL females have lower visceral fat mass and are more glucose tolerant than AL males (Figs. 2G, 4B-C), and are similar to CR males and females in terms of fasting insulin and HOMA-IR (Fig. 4D-E). We also show that, for most sphingolipid species, the hepatic ceramide:DHC ratio is similar between AL females and CR mice of either sex (Fig. S10C), while our RNA-seq data show that CR males are more similar to AL females than AL males (Fig. 7A). These patterns suggest that the mechanisms through which CR decreases visceral adiposity, improves glucose tolerance and modulates sphingolipid metabolism are already at least partially active in young females, even on an AL diet.

This observation helps to clarify potential mechanisms for CR’s age-dependent sex differences. Sexual dimorphism in metabolic function typically results from sex hormones, with genetic factors and/or developmental programming also playing a role. For example, X-chromosome-linked genetic factors influence sex differences in adiposity independently of sex hormones or other secondary sexual characteristics (93, 94), while the developmental and neonatal environment causes sexually dimorphic metabolic programming (95). However, it is unlikely that these developmental or genetic factors explain the sex differences we observe. Developmental programming is unlikely because the early life environment of our mice was the same for both sexes and was devoid of any adverse or stressful stimuli. Similarly, if X-linked genetic factors were responsible then the sex differences would be expected to persist in aged females, but they mostly do not. This leaves sex steroids as the likely mediator. Consistent with this, oestrogen is now well known to influence metabolic function and the fasting response (14, 23), and many actions of oestrogens overlap with those central to the effects of CR (96–99). Indeed, our RNA-seq data show that CR activates oestrogen response pathways in males and females (Fig. 7C), further suggesting commonalities in the effects of oestrogens and CR. Herein, we further show that the sex differences in CR’s metabolic effects are absent or greatly diminished when CR is initiated at 18 months of age, when C57BL/6 females are typically anoestrus (38, 39). Similarly, CR does decrease fat mass in young ovariectomised females (100), while oestrogen treatment protects male mice from obesity-induced insulin resistance (101, 102). Therefore, we propose that young females resist the metabolic effects of CR because, through endogenous oestrogen action, they already have high activation of pathways that mediate CR’s metabolic effects.

### Limitations and further considerations

Our study represents a significant advance in knowledge by revealing that age and sex have a substantial influence over the metabolic benefits of CR. This highlights the need for preclinical and human CR research to address these variables as key determinants of the CR response, a consideration that we show has been lacking in most CR research. We show that these sex differences occur on a systemic, physiological level and that they extend to cellular and molecular effects within adipose tissue and the liver, observations that identify potential mechanisms for the age-dependent sex differences; however, a general limitation of our study is that these mechanisms remain to be directly addressed. Specific limitations are as follows.

Firstly, we used a feeding regimen common in rodent CR studies, with mice receiving a single ration of diet in the morning. We found that the sex differences persisted when the daily ration was given in the evening, demonstrating that the sexual dimorphism is not specific to a morning-fed regimen. However, such single-ration protocols invoke extensive daily fasting, with mice typically consuming all of the food within ∼2 hours (103, 104). Therefore, they include elements of both CR and of intermittent fasting. Recent studies have shown that such prolonged daily fasting contributes to the metabolic benefits and lifespan extension associated with CR, at least in male mice (103, 105, 106). Thus, it will be interesting to determine if these beneficial effects of prolonged fasting, and not just CR *per se,* also differ between males and females.

Secondly, regarding insulin sensitivity and glucose homeostasis, it would be informative to use hyperinsulinaemic-euglycaemic clamps and labelled metabolic tracers to comprehensively assess systemic and hepatic insulin sensitivity and determine if, as we propose, the differences in glucose homeostasis relate primarily to effects on gluconeogenesis and hepatic acetyl-CoA metabolism.

Thirdly, the role of oestrogen remains to be directly tested. This could be done in mice through ovariectomy studies and/or manipulation of ER*α* activity. We have also tried to determine oestradiol concentrations in our mouse plasma samples using mass spectrometry, a method required for reliable assessment of sex steroid concentrations (107). However, despite extensive mass spectrometry optimisation efforts, we have not yet been able to reliably measure oestradiol using the relatively small plasma volumes available. Related to this, in our human studies we were unable to confirm the menopausal status of the female participants. Other limitations of our human studies are that they were done only in overweight or obese individuals; included relatively few younger participants; and lacked analysis of regional adiposity (e.g. visceral versus subcutaneous adipose tissue). These limitations reflect that our human study was not originally designed to directly test the influence of age and sex, with the data instead analysed retrospectively to address this. Future human research must therefore prioritise assessment of menopausal status and regional adiposity, both in lean and overweight/obese subjects, if the outcomes of CR are to be fully interpreted.

A fourth limitation regards the underlying molecular mechanisms. Pathways implicated in mediating CR’s effects include activation of AMPK, SIRT, IGF-1 and FGF21, modulation of mitochondrial function and inhibition of mTOR (5, 19). Thus, genetic, pharmacological and/or other approaches to manipulate these candidate pathways could be used to further dissect the molecular mechanisms underlying CR’s age-dependent sexually dimorphic metabolic effects. Such approaches are a promising basis for future research.

Beyond these specific limitations, a broader question is whether other effects of CR also differ between males and females in this age-dependent manner. Some research suggests that CR’s effects on cardiovascular function and healthy ageing differ between males and females (108–110); however, given the dearth of CR studies that directly assess sex differences (Fig. 1, S1) there remains a critical need to determine how age and sex influence the many other outcomes of CR. This is important not only for understanding fundamental biology, but also if we are to translate the benefits of CR, or CR mimetics, to improve human health. For example, if younger women resist many of CR’s health benefits, should any CR-based intervention focus only on older women, who may be more responsive?

Finally, while it will be useful to understand the extent and basis for these sex differences, it is also important to consider their broader purpose. Adipose tissue profoundly influences female reproductive function, with adipose-secreted factors acting within the central nervous system to indicate if the body’s energy stores are sufficient to support reproduction (111). The hormone leptin is the key adipose-derived factor that impacts the hypothalamic-pituitary-gonadal axis: if fat mass is too low, for example in anorexia nervosa or lipodystrophy, then leptin levels are also insufficient, resulting in decreased gonadotropin secretion and impaired reproductive capacity. Normal menses resumes in anorexic individuals who regain weight and fat mass, or in lipodystrophic individuals following leptin treatment (111, 112). Notably, we previously demonstrated that female mice resist hypoleptinaemia during CR (22). Thus, we speculate that females resist fat loss during CR to preserve reproductive function. Given the tight interplay between adipose tissue and systemic metabolic homeostasis, such resistance to fat loss then has consequences for CR’s other metabolic effects. This evolutionary pressure to overcome periods of food scarcity would thereby have driven the divergence of the CR response in males and females, with implications for current efforts to exploit CR therapeutically to improve human health.

## MATERIALS AND METHODS

### Animals

Studies in C57BL/6NCrl and C57BL/6JCrl mice were approved by the University of Edinburgh Animal Welfare and Ethical Review Board and were done under project licenses granted by the UK Home Office. Mice were originally obtained from Charles River (UK) and then bred and aged in-house. They were housed on a 12 h light/dark cycle in a specific-pathogen-free facility with free access to water and food, as indicated. Differences in mouse cohorts are summarised in Table 2, along with the groups being compared and the experimental unit. The exact number of mice used is stated in the figure legends. Sample sizes for each cohort were determined by power calculations using G*Power software, with effect sizes based on pilot studies and/or previously published data for fat loss and improved glucose tolerance (the primary outcomes being tested) (22, 47); to increase power, data for young mice are reported across multiple cohorts from separate studies, each of which revealed the same sex differences in CR’s metabolic effects. Mice were randomly assigned to each diet. To minimise potential confounders, such as the order of treatments or measurements, mice were treated or analysed based on the order of their ID number, a variable randomly distributed across the experimental variables (diet and sex). Blinding was not possible during the *in vivo* experiments because the experimenter(s) can see, based on presence or absence of diet, which mice are AL and which are CR; however, data analysis was done in a blinded manner. Mice were excluded from the final analysis only if there were confounding technical issues or pathologies discovered at necropsy. Ten mice from the young study were excluded from the analysis of OGTT and necropsy tissue masses because of a technical issue during gavaging for the OGTT, which compromised their subsequent food intake. One mouse was excluded from PET/CT data analysis because it accidentally had two doses of FDG administered. One aged mouse was excluded because of developing confounding liver tumours. For some cohorts, specific tissues were not weighed at necropsy and therefore these mice have no data for these readouts. For further details, please see the Source Data file.

**Table 1.**
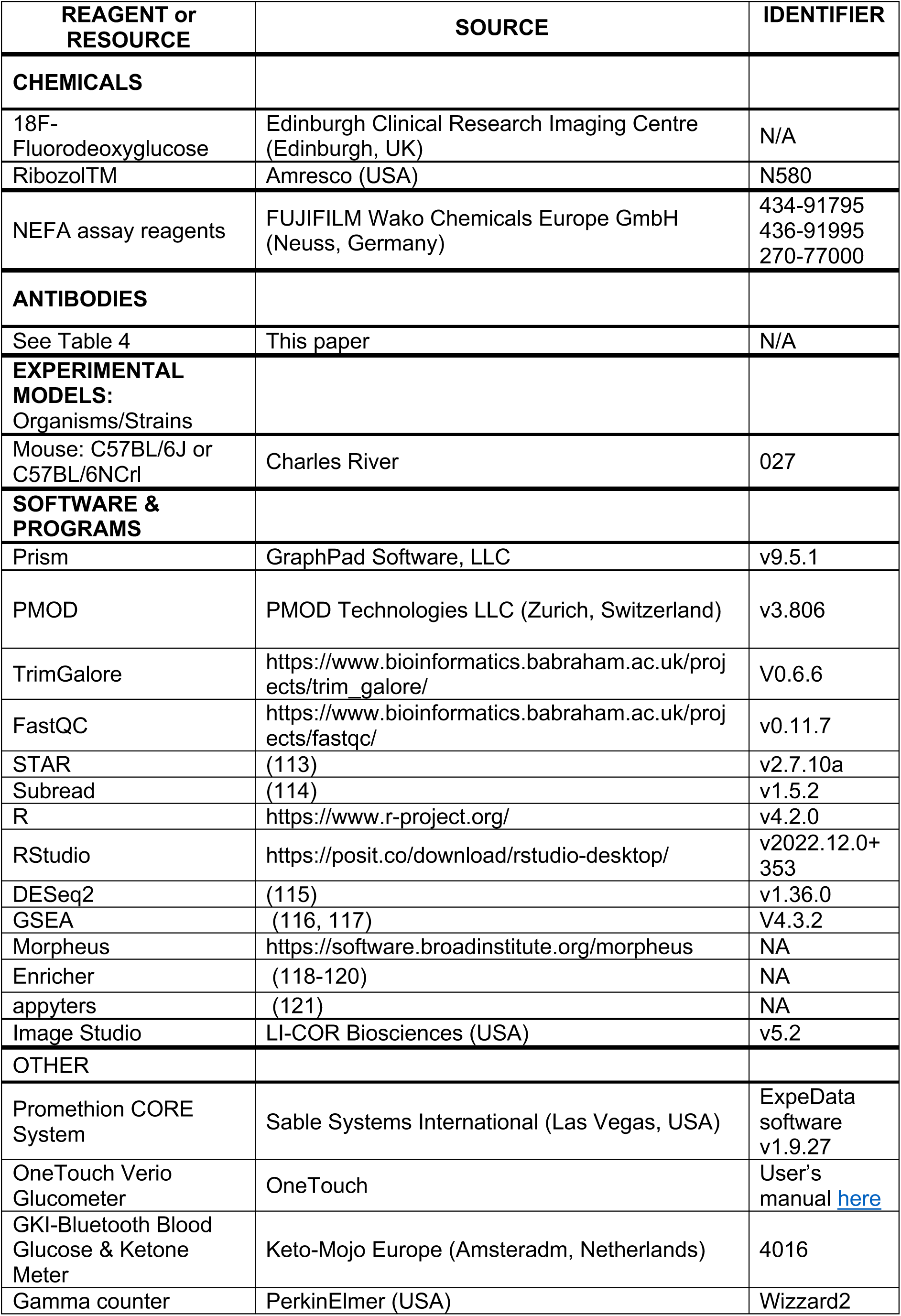
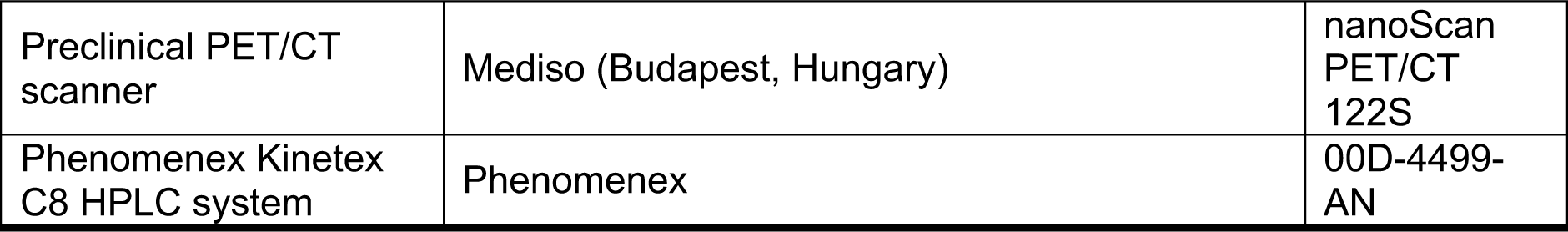
Key Resources

**Table 2.**
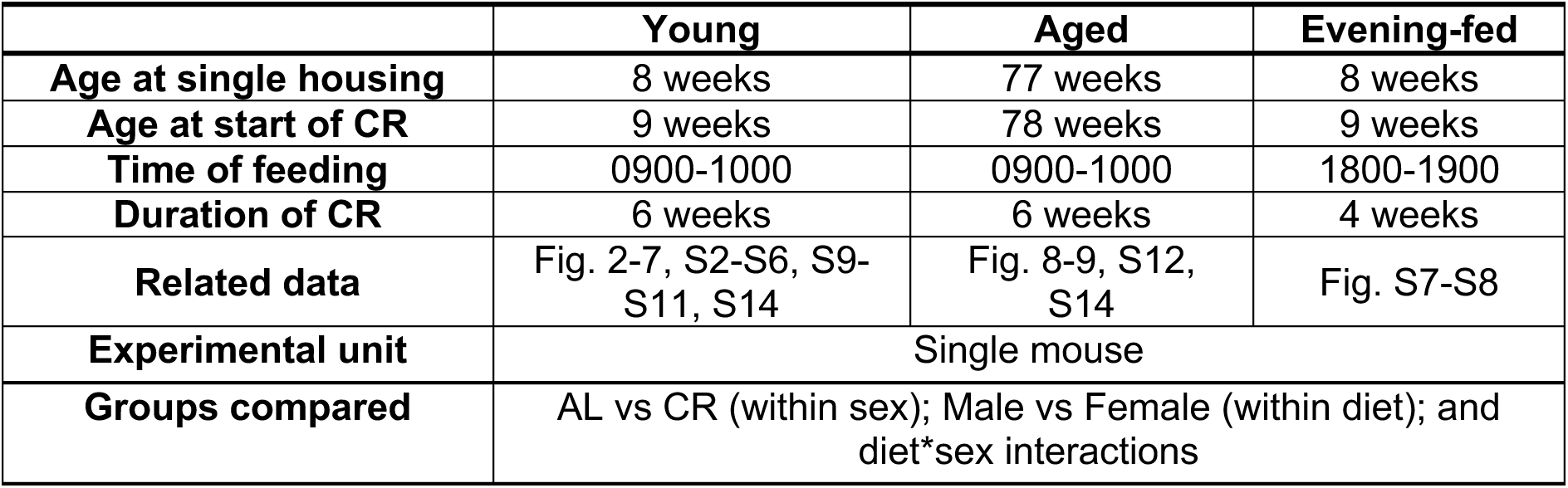
Summary of CR protocol in each group of mice. Because mice are singly housed, each mouse represents an independent experimental unit.

|  | Young | Aged | Evening-fed |
| --- | --- | --- | --- |
| **Age at single housing** | 8 weeks | 77 weeks | 8 weeks |
| **Age at start of CR** | 9 weeks | 78 weeks | 9 weeks |
| **Time of feeding** | 0900-1000 | 0900-1000 | 1800-1900 |
| **Duration of CR** | 6 weeks | 6 weeks | 4 weeks |
| **Related data** | Fig. 2-7, S2-S6, S9-S11, S14 | Fig. 8-9, S12, S14 | Fig. S7-S8 |
| **Experimental unit** | Single mouse |  |  |
| **Groups compared** | AL vs CR (within sex); Male vs Female (within diet); and diet*sex interactions |  |  |

### Mouse CR studies

Before assigning to AL or CR diets, mice were housed individually between 8-9 weeks of age (for young mice) or 77-78 weeks of age (for old mice). During this time, mice were fed a control diet (Research Diets, D12450B) *ad libitum* and daily food intake was determined by weighing the food remaining in each cage each day. Following one week of single housing, all mice were randomly assigned to control diet (Research Diets D12450B, provided AL) or the CR diet (Research Diets D10012703, administered at 70% of the average daily AL diet consumption). This CR diet is formulated for iso-nutrition at 70% CR, ensuring that malnutrition is avoided. The evening-fed CR mice (Fig. S5-S6) were fed daily between 1800-1900, while all other CR mice were fed daily between 0900-1000. We find that CR mice consume their entire ration of diet within 2-3 hours of feeding, consistent with other CR studies (103, 104) For the first two weeks of CR mice were monitored and weighed daily to ensure they did not lose more than 30% of their initial body mass. For the following 4 weeks of CR, body mass was recorded weekly. Mice were assessed non-invasively for body fat, lean mass, and free fluid weekly from weeks 0 to 6 using TD-NMR (Minispec LF90II; Bruker Optics, Billerica, MA, USA). Blood ketone concentrations were measured from weeks 0 to 4 using a blood ketone meter (Keto-Mojo Europe, Amsterdam, Netherlands). At necropsy, mice were humanely sacrificed via cervical dislocation and decapitation. Tissues were harvested, weighed, and stored either in 4% formalin or on dry ice. Tissues in formalin were fixed at 4°C for 2-3 days before being washed and stored in DPBS (Life Technologies). Tissues on dry ice were stored at −80°C prior to downstream analysis.

### Human weight loss study – Participants

This study was done before pre-registration was a requirement for clinical trials. Herein, we retrospectively analysed the study outcomes to investigate if sex and/or age influence the CR response. Forty-five overweight and obese males and females; age 21-61y, Body Mass Index (BMI) 26-42 kg/m^2^, were recruited by newspaper advertisement to participate in a weight loss (WL) dietary intervention study. Thus, the participants were non-randomly selected individuals who were sufficiently motivated to actively respond to the request for volunteers. During recruitment, participants underwent a medical examination and their general practitioners were contacted to confirm medical suitability to participate in the study. Written, informed consent was obtained, and the study was reviewed by the NHS North of Scotland Research Ethics Service (ethics number 09/S0801/80). Participants were informed that the study was funded by an external commercial sponsor but the company was not named, and all packaging of the food range was removed before distribution. Three participants withdrew for personal reasons therefore data for *n* = 42 is presented throughout; participant characteristics are summarised in Table 3. No data points were excluded from the final analyses.

**Table 3.**
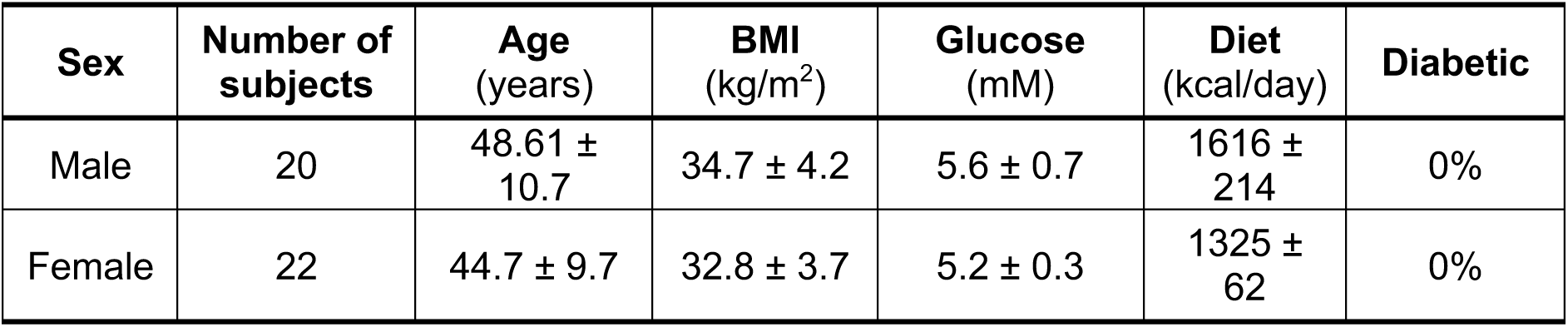
Human subject characteristics at baseline. Age, BMI, glucose and daily caloric intake (Diet) are mean ± SD. *ND* = not determined.

| Sex | Number of subjects | Age (years) | BMI (kg/m<sup>2</sup>) | Glucose (mM) | Diet (kcal/day) | Diabetic |
| --- | --- | --- | --- | --- | --- | --- |
| Male | 20 | 48.61 ± 10.7 | 34.7 ± 4.2 | 5.6 ± 0.7 | 1616 ± 214 | 0% |
| Female | 22 | 44.7 ± 9.7 | 32.8 ± 3.7 | 5.2 ± 0.3 | 1325 ± 62 | 0% |

### Human weight loss study – Protocol

Participants were asked to attend the Human Nutrition Unit (HNU) at the Rowett Institute of Nutrition and Health (Aberdeen, UK) twice weekly to pick up food supplies. Participants had their basal energy requirements determined and each participant was then fed an individualised diet with a caloric content equivalent to 100% of their resting metabolic rate (Table 3). This approach was taken to standardise the diet to account for individual energy requirements and energy restriction. All required food, drinks and snacks were provided and the study dietician was present to answer any questions. Participants were provided with a diet guide whereby women were encouraged to consume no more than 6.3 MJ/d (1,500 kcal) and men 8.4 MJ/d (2,000 kcal). They were encouraged to consume 3 meals a day (breakfast, lunch and dinner) with a portion of protein at each meal, and 5 portions of fruit and vegetables, 2 portions of fats, 3 portions of milk and dairy and 4 portions of wholegrain bread, potatoes and cereals each day. Participants completed a descriptive food diary every day to record what and when they ate. Participants ate *ad libitum*, choosing when and what to eat from their supplies. They completed a detailed 7d weighed dietary record during week 4 of the WL phase of the study, to compare eating habits with a baseline 7d weighed record they completed before the study started. Energy and nutrient intakes were calculated using the Windiets programme (Univation Ltd; The Robert Gordon University, Aberdeen, UK).

### Human weight loss study – Measurement of body composition

Before and after WL, at weeks 0 and 4 respectively, measurements of body composition were conducted under standardized conditions as described previously (122), with participants instructed to fast overnight (at least 10 h). Height was measured at the beginning of the study to the nearest 0.5 cm using a stadiometer (Holtain Ltd, Crymych, Dyfed, Wales). The body weight of the participants was measured at screening, during food provision days and on all test days, wearing a previously weighed dressing gown, to the nearest 100 g on a digital scale (DIGI DS-410, C.M.S. Weighing Equipment Ltd, London, UK). Fat mass (FM) and fat free mass (FFM) were measured with the use of air displacement whole-body plethysmography (BodPod® Gold – Body Composition System, Cosmed, Italy) (123).

### Indirect calorimetry

Male and female AL and CR mice were housed individually in Promethion CORE System cages (Sable Systems International) from 9-10 or 12-13 weeks of age, corresponding to weeks 0-1 (“week 1”) or 2-3 (“week 3”) after beginning AL or CR feeding. At each timepoint, mice entered the cages around 11 am on day 1 and were housed for three nights (four days in total). Data herein are from the 48 h period between 7am on day 2 and 7am on day 4, to allow habituation to the new cages during day 1. Before and after Promethion housing, each mouse was weighed and body composition determined by TD-NMR. Measurements of energy expenditure, oxygen consumption, carbon dioxide production, and physical activity were analysed using ExpeData software according to the manufacturer’s instructions. FA oxidation was determined as described previously (26).

### Oral glucose tolerance tests (OGTT)

Mice were given a half portion of food the night (6pm) prior to the metabolic challenge. At 12:00 pm on the day of the OGTT a basal blood sample was collected by venesection of the lateral tail vein into EDTA-coated capillary tubes (Microvette 16.444, Sarstedt, Leicester, UK) and the basal glucose levels was measured using a glucometer (OneTouch Verio IQ, LifeScan Inc., Zug, Switzerland). D-glucose (G8270, Sigma, Poole, UK) was then administered at 2 mg per g of body mass by oral gavage of a 25% (w/v) solution. At 15, 30, 60 and 120 minutes post-gavage, blood glucose was measured by a glucometer and tail vein blood was sampled in EDTA tubes, as for the basal (0 min) timepoint. Blood samples were kept on ice prior to isolation of plasma. Insulin was then measured in plasma samples by ELISA, following the manufacturer’s instructions (cat. #90080, ChrystalChem, Chicago, IL, USA). Insulin was converted from ng/ml to μU/mL by multiplying the ng/mL value by 28.7174 (based on mouse insulin molar mass of 5803.69). Homeostatic Model Assessment for Insulin Resistance (HOMA-IR) and Matsuda index were calculated as previously described (27, 28)).

### PET/CT scanning

Immediately before and 15- and 60-min post ^18^F-FDG intraperitoneal injection, blood glucose was measured by tail venesection and blood was sampled directly into EDTA-microtubes (Sarstedt, Leicester, UK). At 60-minutes post ^18^F-FDG injection, mice were anaesthetised and ^18^F-FDG distribution was assessed by PET/CT. Following the PET/CT scan, mice were sacrificed by overdose of anaesthetic (isoflurane), death was confirmed by cervical dislocation and tissues of interest were then dissected in a lead-shielded area. Half of each of the dissected tissues were snap frozen on dry ice or placed into 10% formalin. ^18^F-FDG uptake was then determined using a gamma counter (PerkinElmer) as previously described (31)

### PET/CT analysis

PET/CT images were reconstructed and data analysed using PMOD version 3.806 (PMOD, Zurich, Switzerland). SUVs were calculated for regions of interest, namely BAT, iWAT, gWAT; heart; bone tissue (without BM) from tibiae, femurs, and humeri; To distinguish bone tissue from BM, a calibration curve was generated using HU obtained from the acquisition of a CT tissue equivalent material (TEM) phantom (CIRS, model 091) and mouse CT scans, as previously described (31).

### Histology and Adipocyte size quantification

Dissected mouse tissue was fixed for 48 h and paraffin embedded by the histology core at The University of Edinburgh’s Shared University Research Facilities (SuRF). Paraffin-embedded tissue sections were then sectioned at 100 µm intervals using a Leica RM2125 RTS microtome and collected onto 76 x 26 mm StarFrost 624 slides (VWR, UK). The slides were baked at 37°C overnight before Haematoxylin and Eosin (H&E) staining. Quantification of adipocyte number and size was performed on H&E-stained adipose tissue using the ImageJ plug-in Adiposoft, an automated software for the analysis of adipose tissue cellularity in histological sections (124).

### NEFA analysis in plasma samples

NEFA standard solution (Wako, 270-77000) was serially diluted to concentrations of 2, 1.5, 1, 0.5, 0.25 and 0.125 mM. The standard dilutions, blank sample and plasma (2 μL) were pipetted into separate duplicate wells of a 96-well plate. Reagent 1 (200 μL) (Wako, 434-91795) was added to each well and the plate incubated at 37°C for 5 min. Absorbance of each well was measured at 550 and 660 nm. Reagent 2 (100 μL) (Wako, 436-91995) was then added to each well and the plate was incubated again at 37°C for 5 min and absorbance measured at 550 and 660 nm. The 660 nm reading was subtracted from the 550 reading from both measurements and the mean blank absorbance subtracted from the mean duplicate absorbance from each sample. NEFA concentrations were then determined by regression analysis.

### Protein isolation and western blotting

Frozen WAT samples were lysed in preheated SDS lysis buffer (1% sodium dodecyl sulfate (SDS), 0.06 M Tris pH 6.8, 12.7 mM ETDA, 1 mM PMSF, 1 mM sodium orthovanadate). After tissue disruption, the lysate was boiled at 95 °C on a dry heat block for 10 min. Boiled lysate was centrifuged at 16,000×*g* and the cleared liquid fraction below the lipid layer and above the pellet was removed to a new tube for downstream analyses. Protein concentration was quantified using the BCA protein assay (Pierce 23225). For SDS polyacrylamide gel electrophoresis, lysates were diluted to equal protein concentration in lysis buffer plus 4X SDS loading buffer (4% SDS, 240mM Tris-HCL ph6.8, 40% glycerol, 0.276%(w/v) beta-mercaptoethanol, 0.05% Bromophenol blue). Dithiothreitol was added to the lysates (25 mM final concentration). Samples were boiled for 5 min, cooled on ice for 1 min, vortexed, and equal amounts of protein (13 µg per lane) separated on gradient polyacrylamide gels (Bio-Rad, Hercules, CA, USA). Samples were then transferred to Immobilon-FL transfer membrane (Millipore, Billerica, MA, USA). After transfer, membranes were blocked in 5% milk for 1 h at RT. Membranes were incubated in primary antibodies (Table 4; each in 5% BSA) overnight at 4 °C, followed by incubation with secondary antibody (Table 4) at 1:5,000 dilution for 1 h at RT. Fluorescent color was detected with Odyssey CLx Imager (LI-COR, Lincoln, NE, USA). Band signal was quantified using Image Studio v5.2 (LI-COR, Lincoln, NE, USA). Quantified data obtained from two different membranes were combined by using the strength of the signal of the ladder bands as a control for intensity differences between the two membranes.

**Table 4.**
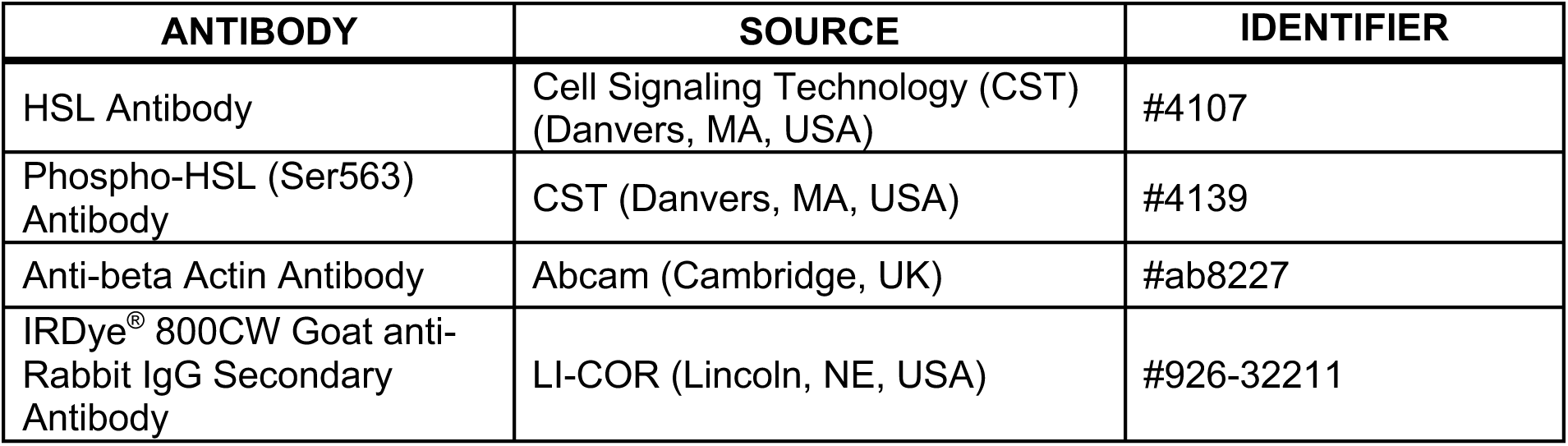
Antibodies used

| ANTIBODY | SOURCE | IDENTIFIER |
| --- | --- | --- |
| HSL Antibody | Cell Signaling Technology (CST)<br>(Danvers, MA, USA) | #4107 |
| Phospho-HSL (Ser563)<br>Antibody | CST (Danvers, MA, USA) | #4139 |
| Anti-beta Actin Antibody | Abcam (Cambridge, UK) | #ab8227 |
| IRDye® 800CW Goat anti-Rabbit IgG Secondary<br>Antibody | LI-COR (Lincoln, NE, USA) | #926-32211 |

### Liver ceramide quantification

Liver ceramide and dihydroceramide content was determined using LC-MS (Lipidomics Core Facility; University of the Highlands and the Islands, Inverness, UK). As per Folch *et al.* 1957 (125), 100 mg of frozen liver tissue was homogenised in ice-cold MeOH using a mechanical tissue homogeniser. Fifty microlitres of each sample was combined with 100 μL ceramide 17:0 (1 μM in MeOH) and 25 μL dihydroceramide 12:0 (2 μM in MeOH) (Avanti Polar Lipids, Alabaster, AL, USA) as internal standards. MeOH (1.875 mL) and CHCl_3_ (4 mL) were added. Samples were then stored on ice for 1 hour with intermittent vortex mixing, centrifuged at 2,000 rpm for 5 minutes at 4°C and the supernatant was collected. KCl (0.9%; 1.45 mL per sample) was added to the supernatant and the samples were centrifuged at 2,000 rpm for 5 minutes at 4°C. The lower phase was collected and evaporated under nitrogen and reconstituted in 1 mL CHCl_3_. Solid phase extraction was performed via a 100 mg/3 mL Isolute silica column (Biotage, Uppsala, Sweden). The column was conditioned twice with 3 mL CHCl_3_ and then loaded with 1 mL of the sample. The bound sample was washed with 2 washes of 3 mL CHCl_3_ then eluted with 2 additions of ethylacetate CHCl_3_ (1:1 v/v). The collected eluate was evaporated and dried under vacuum and reconstituted in 200 μL MeOH. LC-MS/MS analyses were performed in positive ion mode on a Thermo TSQ Quantum Ultra triple quadrupole mass spectrometer equipped with a HESI probe and coupled to a Thermo Accela 1250 UHPLC system. The ceramides and dihydroceramides were separated on a Kinetex C8 column (1.7 μm; 100 × 2.1 mm) (Phenomenex, Macclesfield, UK). Mobile phase A was 90% H_2_O, 9.9% acetonitrile and 0.1% formic acid. Mobile phase B was 99.9% acetonitrile and 0.1% formic acid. The gradient was held at 80% mobile phase B for 1 minute then gradually increased to 100% mobile phase B over 15 minutes. Mobile phase B at 100% was maintained for 1 minute and then equilibrated to storage conditions. The flow rate used was 500 μL per minute at 40°C. Two µL of each sample was injected into the LC-MS/MS for ceramide analysis and 10 μL for dihydroceramide. Total ceramide and dihydroceramide concentrations were calculated from the summed concentrations of all the monitored molecular species. All values were normalised to wet weight of liver. Principal component analysis for ceramide and dihydroceramide data was done using RStudio (RStudio Team, USA), using the code provided (see ‘Data and code availability”, below).

### RNA-seq

RNA was isolated from tissues using Ribozol^TM^ solution (cat. No. N580, Amresco, USA,) according to the manufacturer’s protocol. RNA quantity and quality were quantified using a NanoDrop spectrophotometer (Thermo Scientific,USA) and RNA Integrity Number (RIN) was assessed using Agilent RNA 6000 Pico Kit and an Agilent 2100 bioanalyzer (Agilent, USA). Library preparation and sequencing was performed by BGI (Shenzhen, Guangdong, China); 100 bp pair-end sequencing was done using a DNBseq-G400 (MGI Tech). Calculations were done using Eddie Mark 3, the University of Edinburgh’s high-performance computing cluster. After the trimming by TrimGalore and quality control by FastQC, sequences were aligned to the mouse genome (mm10) with the annotation data from the website of University of California Santa Cruz (https://hgdownload.soe.ucsc.edu) using STAR (v2.7.10a) (113). Mapped reads were counted using featureCounts in Subread package (v1.5.2) (114) and subsequent analyses performed using R, RStudio, GSEA, Enricher, and Morpheus. Count data were normalised and analysed using DESeq2 (v1.36.0) to detect differentially expressed genes (115). Principal component analysis was performed using the prcomp function for 500 genes with the highest variation among samples after transforming the raw count data using the vst function from the DESeq2 package. R code used for the analysis are provided (see ‘Data and code availability”, below). For GSEA, a gene set collection including 8 gene sets was used for Fig. 7C; further details are in table 5. Heatmaps were drawn with Morpheus.

**Table 5:**
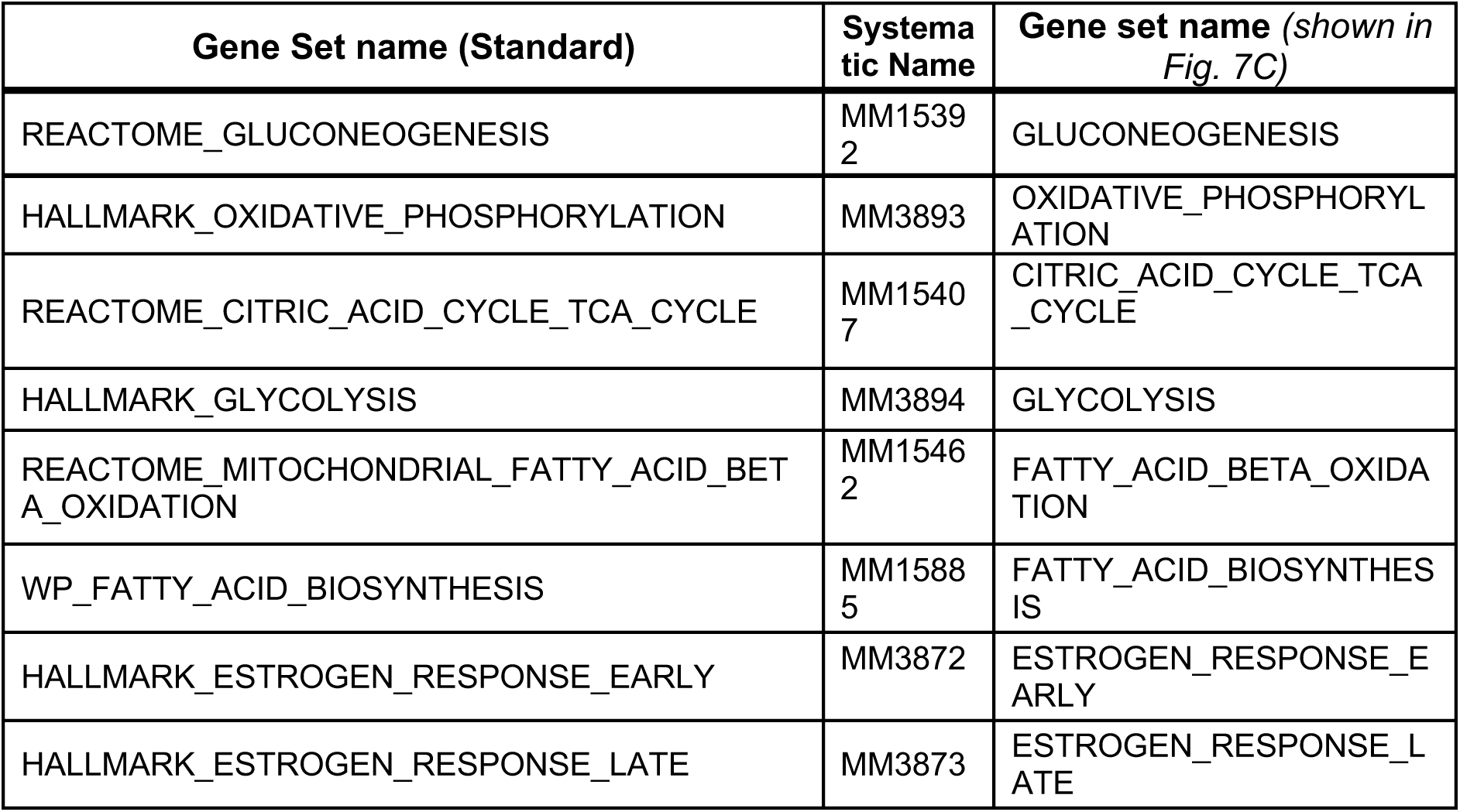
Gene sets used for GSEA. Further details can be found by searching for the Standard Gene Set name at http://www.gsea-msigdb.org/gsea/msigdb/mouse/genesets.jsp

### Statistical analysis

Data were analysed for normal distribution using the Shapiro-Wilk normality test. Normally distributed data were analysed by ANOVA, mixed models or t-tests, as appropriate; mixed models were used for repeated-measures analyses in which some data points were missing or had to be excluded for some mice. Where data were not normally distributed, non-parametric tests were used. When appropriate, *P* values were adjusted for multiple comparisons. Data are presented as mean ± SEM or as Violin plots, as indicated in the figure legends. All statistical analyses were performed using Prism software (GraphPad, USA). A *P*-value <0.05 (after adjustment for multiple comparisons) was considered statistically significant.

### Data and code availability

All source data and code from which the figures are based is available through University of Edinburgh DataShare at https://doi.org/10.7488/ds/3817. Raw RNAseq data will be deposited in a public repository. Any other relevant data are available from the authors upon reasonable request.

## Supporting information

Response to Review Commons reviewers

## ACKNOWLEDGEMENTS

This work was supported by grants from the Medical Research Council (MR/M021394/1 to W.P.C.), the British Heart Foundation (BHF) (RG/16/10/32375 and FS/19/34/34354 to A.A.S.T.; 4-year BHF PhD Studentship to B.J.T. and R.J.S.), the Takeda Science Foundation (Fellowship for Young Japanese MDs & PhDs Studying Abroad, to Y.M.I.), the Japan Society for the Promotion of Science (JSPS Overseas Research Fellowship, to Y.M.I.), the Japan Foundation for Applied Enzymology (to Y.M.I.), the University of Edinburgh (Chancellor’s Fellowship to W.P.C.; PhD Studentship to A.L.), the Carnegie Trust (RIG007416 to W.P.C.), the Wellcome Trust (Technology Development Award 221295/Z/20/Z to A.A.S.T.; Multi-user equipment grant 223818/Z/21/Z for the Promethion system), the Chief Scientist Office of the Scottish Government (SCAF/17/02 to R.H.S.), KAKENHI grants from MEXT/JSPS (21K08431 to H.K.), and a grant from the National Center for Global Health and Medicine (20A1010 to H.K.). C.F. and A.M.J. gratefully acknowledge financial support from the Scottish Government as part of the RESAS Strategic Research Programme at the Rowett Institute, University of Aberdeen. We thank Ami Onishi for assistance with indirect calorimetry studies using the Promethion system, and the BHF for supporting Ami Onishi’s salary through the Centre for Research Excellence Award III (RE/18/5/34216). The BHF also provided funding towards establishment of the Edinburgh Preclinical PET/CT laboratory (RE/13/3/30183) and support of the 2018 Very Important Project (VIP) PET prize, for which we are grateful. Finally, we thank Dr Matthew Bennett (University of Edinburgh) for advice analysing RNA-seq data; Prof Robert Semple (University of Edinburgh) and Prof Ormond MacDougald (University of Michigan) for helpful advice to interpret our findings; Stefanie Fung (University of Edinburgh) for assistance isolating RNA for RNA-seq; Dr Hai-Bin Ruan (University of Minnesota Medical School) and Dr Eleonora Pagano (Pontifical Catholic University of Argentina) for kindly providing a full text of Ballor and Poehlman’s 1994 study (78), and all staff at Edinburgh Bioresearch & Veterinary Services for their superb technical support.

## Rights Retention Statement

For the purpose of open access, the author has applied a Creative Commons Attribution (CC-BY) licence to any Author Accepted Manuscript version arising from this submission.

## AUTHOR CONTRIBUTIONS

Contributions are based on the CRediT (Contributor Roles Taxonomy) and are as follows: ***Conceptualisation***, K.J.S., B.J.T., G.G., R.J.S. A.M.J. and W.P.C.; ***Data curation,*** K.J.S., B.J.T., Y.M.I., K.C.C., C.F., R.J.S., A.L., A.M.J. and W.P.C.; ***Formal Analysis***, K.J.S., B.J.T., Y.M.I., K.C.C., C.F., R.J.S., A.L., H.K., P.D.W., N.M.M. and W.P.C.; ***Funding Acquisition***, K.J.S, A.A.S.T., G.A.G., N.M.M., R.H.S., A.M.J. and W.P.C.; ***Investigation***, K.J.S., B.J.T., Y.M.I., K.C.C., C.F., R.J.S., A.L., H.J.W., X.H., D.M., E.J.B., C.J.A-C., P.D.W., A.M.J. and W.P.C.; ***Methodology***, K.J.S., B.J.T., Y.M.I., K.C.C., A.A.S.T., H.K., P.D.W., N.M.M., A.M.J. and W.P.C.; ***Project administration,*** G.A.G., N.M.M., A.M.J. and W.P.C.; ***Resources***, A.A.S.T., G.A.G., N.M.M., R.H.S., A.M.J. and W.P.C.; ***Supervision,*** G.A.G. N.M.M., R.H.S., A.M.J. and W.P.C.; ***Visualisation***, K.J.S., B.J.T., Y.M.I., K.C.C., and W.P.C.; ***Writing – Original Draft***, K.J.S., B.J.T., Y.M.I. and W.P.C.; ***Writing – Review & Editing***, K.J.S., B.J.T., Y.M.I., G.A.G., R.H.S., N.M.M., A.M.J. and W.P.C.

## SUPPLEMENTARY FIGURE LEGENDS

**Supplementary Figure 1, Related to Figure 1 – Methods used and additional outcomes relating to the comprehensive review of the CR literature.**

The NIH PubMed database was searched using MeSH terms to identify research articles that studied CR *in vivo* in mice or humans, published between 2003 and 2021. **(A)** Method of database search presented as a flow diagram, including search terms, reasons for exclusion, and number of results in each category. **(B)** Percentage of mouse or human publications for each year, with publications classified into each group as described for Figure 1. Source data for (B) are provided as a Source Data file.

**Supplementary Figure 2, Related to Figure 2 – The effect of CR on percent fat and lean mass differs between male and female mice.**

Male and female mice were fed AL or CR diets and body composition was assessed weekly, as described for Figure 2. **(A,B)** Fat mass (A) and lean mass (B) were determined by TD-NMR and calculated as % body mass. **(C)** Masses of iWAT (inguinal WAT), mWAT (mesenteric WAT), pWAT (perirenal WAT), pcWAT (pericardial WAT) and BAT (brown adipose tissue) were recorded at necropsy. Data in (A-B) are shown as mean ±SEM of the following numbers of mice per group: *male AL*, n=42; *female AL,* n=43; *male CR,* n=44; *female CR,* n=52. Data in (C) are shown as violin plots of the following numbers of mice per group and tissue type: *male AL,* n=29 (iWAT, mWAT, BAT), 15 (pWAT) or 14 (pcWAT); *female AL,* n=28 (iWAT, mWAT, BAT), 14 (pWAT) or 13 (pcWAT); *male CR,* n=33 (iWAT, mWAT, BAT), 16 (pWAT) or 15 (pcWAT); *female CR,* n=34 (iWAT, mWAT, BAT), 15 (pWAT) or 14 (pcWAT). For (A-B), significant effects of diet, sex and/or duration were determined by 3-way ANOVA (A) or a mixed-effects model (B). For (C), significant effects of diet and/or sex within each tissue were determined by 2-way ANOVA with Tukey’s multiple comparisons tests; overall ANOVA *P* values are shown beneath each graph. Source data are provided as a Source Data file.

**Supplementary Figure 3, Related to Figure 2 – CR and sex effects on the masses of adipose and non-adipose tissues.**

Male and female mice were fed AL or CR diets, as described for Figure 2. After six weeks of CR (15 weeks of age) tissues were sampled and weighed. **(A-C)** Violin plots, overlaid with individual data points, showing the % body mass (A,C) or absolute mass (B) of each tissue; Gastroc = gastrocnemius muscle. For (A), the numbers of mice per group and tissue type are as reported for Figures 2G and S2C. For (B) and (C), the following numbers of mice were used for each group and tissue type: *male AL,* n=29 (liver, spleen), 23 (pancreas, kidneys, heart, thymus), 22 (adrenals) or 16 (gastroc.); *female AL,* n=33 (liver, spleen), 27 (pancreas, kidneys, heart, thymus, adrenals) or 16 (gastroc.); *male CR,* n=28 (liver, spleen), 22 (pancreas, kidneys, heart, thymus, adrenals) or 14 (gastroc.); *female CR,* n=34 (liver, spleen), 28 (pancreas, kidneys, heart, thymus, adrenals) or 15 (gastroc.). For each tissue, significant effects of diet and/or sex were determined by 2-way ANOVA with Tukey’s multiple comparisons tests and are reported as described for Figure 2. Source data are provided as a Source Data file.

**Supplementary Figure 4, Related to Figure 2 – CR decreases adipocyte size and stimulates lipolysis in male but not in female mice.**

Male and female mice were fed AL or CR diets as described for Figure 2. **(A,B)** After six weeks of CR (15 weeks of age) mWAT was sampled and processed for histological analysis. Micrographs of H&E-stained sections of mWAT (A) were used for histomorphometric analysis of adipocyte area (B); in (A), scale bar = 100 µm. **(C)** At 10 weeks of age, during the period of maximal weight and fat loss, a separate cohort of mice were culled and iWAT was collected for Western Blot analysis, as described for Figure 2. Expression of phosphorylated HSL (P-HSL) and total HSL was quantified using LICOR software and are shown normalised to β-actin expression. Data in (A) show representative micrographs. Data in (B) and (C) are shown as violin plots from the following number of mice per group: *male AL,* n=5 (B) or 11 (C); *female AL,* n=6 (B) or 10 (C); *male CR*, n=6 (B) or 11 (C); *female CR,* n=6 (B) or 13 (C). For (B), significant effects of diet and/or sex on adipocyte area (Ad.Ar) were determined by 3-way ANOVA, with *P* values shown beneath the graph. Significant differences in median Ad.Ar between AL and CR mice were further determined 2-way ANOVA with Šídák’s multiple comparisons test; * = *P*<0.05. For (C), significant differences were determined by 2-way ANOVA with Šídák’s multiple comparisons test and are reported as described for Figure 2. Source data are provided as a Source Data file.

**Supplementary Figure 5, Related to Figure 3 – Effects of CR on energy expenditure and physical activity.**

Male and female AL and CR mice underwent indirect calorimetry as described for Figure 3. **(A)** Overall energy expenditure relative to lean mass (kcal/g) during the day, night, or day + night (Total) for Week 1 and Week 3. **(B)** Linear regression of lean mass vs Total energy expenditure (kcal/24 h) during Week 3. **(C)** Distance moved by each mouse during the day, night, or day + night (total) for Week 1 and Week 3. Data presentation and statistical analysis are as described for Figure 3. Source data are provided as a Source Data file.

**Supplementary Figure 6, Related to Figure 3 – Effects of CR on Respiratory exchange ratio and FA oxidation**

Male and female AL and CR mice underwent indirect calorimetry as described for Figure 3. **(A)** Average RER during the day or night for Weeks 1 and 3. **(B)** Average RER during the fasted period, from 04.00-09.00, for Weeks 1 and 3. **(C)** Absolute FA oxidation was determined from energy expenditure and RER (26) and is shown as kcal/h over the 24 h day night periods, based on the average for days 2-4 of Promethion housing, for Week 1 (left) and Week 3 (right). Data presentation and statistical analysis are as described for Figure 3. Source data are provided as a Source Data file.

**Supplementary Figure 7, Related to Figure 2 – Sex differences in the effects of CR on body mass and composition persist when CR mice are fed in the evening.**

Male and female C57BL6/NCrl mice were single housed and fed AL or CR diet (70% of daily AL intake) from 9-13 weeks of age, with CR mice receiving their daily diet ration between 18:00-19:00. **(A-F)** Each week mice were weighed (A,B) and body composition was determined by TD-NMR (C-F). Body mass, fat mass and lean mass are shown as absolute masses (A,C,E) or fold-change relative to baseline (B,D,F). **(G,H)** Masses of the indicated tissues were recorded at necropsy and are shown as % body mass. Data in (A-H) represent 5 (*male AL, female AL*) or 6 (*male CR, female CR*) mice per group and are shown as mean ±SEM (A-F) or as truncated volcano plots overlaid with individual data points (G,H). For (A-F), significant effects of diet, sex or time, and interactions thereof, were determined by 3-way ANOVA. For (G-H), significant effects of sex, diet, and sex-diet interaction were assessed using 2-way ANOVA with Tukey’s multiple comparisons test. Overall *P* values for each variable, and their interactions, are shown beneath each graph. For (G-H), statistically significant differences between comparable groups are indicated by ** (*P*<0.01). Source data are provided as a Source Data file.

**Supplementary Figure 8, Related to Figure 4 – Sex differences in the effects of CR on glucose homeostasis persist when CR mice are fed in the evening.**

Male and female C57BL6/NCrl mice were fed AL or CR diet from 9-15 weeks of age, as described for Supplementary Figure 5. **(A,B)** At 13 weeks of age mice underwent OGTTs. Blood glucose was recorded at each timepoint (A) and the tAUC and iAUC was determined (B). Data represent 5 (*male AL, female AL*) or 6 (*male CR, female CR*) mice per group and are shown as mean ±SEM (A) or as truncated volcano plots overlaid with individual data points (B). In (A), significant effects of diet, sex or time, and interactions thereof, were determined by 3-way ANOVA. In (B), significant effects of sex, diet, and sex-diet interaction were assessed using 2-way ANOVA with Tukey’s multiple comparisons test. Overall *P* values for each variable, and their interactions, are shown beneath each graph. For (B), statistically significant differences between comparable groups are indicated by ** (*P*<0.01). Source data are provided as a Source Data file.

**Supplementary Figure 9, Related to Figure 5 – The effects of CR on glucose uptake in peripheral tissues.**

Male and female C57BL6/NCrl mice were fed AL or CR diet and glucose uptake was assessed by ^18^F-FDG PET/CT, as described for Figure 5. **(A,B)** ^18^F-FDG uptake in the indicated tissues was determined by PMOD analysis of PET/CT scans (A) or gamma counting of dissected whole tissues (B). Data in (A) and (B) are presented as truncated violin plots, overlaid with individual data points, from 7 (*male AL, female AL, male CR*) or 6 (*female CR*) mice per group; in (B), data for spleen are from 6 mice per group only. Statistical significance was determined as described for Figure 5. Source data are provided as a Source Data file.

**Supplementary Figure 10, Related to Figure 6 – CR exerts sexually dimorphic effects on hepatic sphingolipid content.**

Male and female mice were fed AL or CR diets as described for Figure 2. **(A-C)** At necropsy, livers were sampled for analysis of ceramide and dihydroceramide content by LC-MS. Data show the concentration of each species of ceramide (A), dihydroceramide (B), or dihydroceramide:ceramide ratio (C) and are presented as truncated violin plots showing all points. The following numbers of mice were used per group and species: *male AL*, n=6 (all except: DHC 20:1 & 26:0, Cer:DHC 20:1 & 26:0), 1 (DHC & Cer:DHC 20:1), or 0 (DHC & Cer:DHC 26:0); *female AL,* n=6; *male CR,* n=6 (all except DHC & Cer:DHC 26:0), or 1 (DHC & Cer:DHC 26:0); *female CR,* n=6. For each species, significant effects of sex, diet, and sex-diet interaction were assessed using 2-way ANOVA; overall *P* values for each variable, and their interactions, are shown beneath each graph. Significant differences between comparable groups were assess using Tukey’s multiple comparison test and are indicated by * (*P*<0.05), ** (*P*<0.01) or *** (*P*<0.001). Source data are provided as a Source Data file.

**Supplementary Figure 11, Related to Figure 7 – Effects of sex and diet on the hepatic transcriptome.**

Male and female C57BL6/NCrl mice were fed AL or CR diet from 9-13 weeks of age, as described for Figure 2. Livers were sampled at necropsy (13 weeks) and analysed by RNA-seq. **(A)** Volcano plots showing differentially expressed genes between AL and CR mice of both sexes (left), AL males vs CR males (middle), and AL females vs CR females (right). Transcripts encoding key proteins in gluconeogenesis, glycolysis, or the TCA cycle are shown on each graph. The names of genes with positive fold-change values are shown on the right side and those with negative fold-change values are shown on the left side of the volcano plots. Genes with absolute fold-change > 1.5 and *P*adj < 0.05 are marked with an asterisk next to the gene name. **(B)** List of genes not shown in the volcano plots of Fig. 7B and S11A because their higher values of absolute fold-change and/or *P*adj do not fit on the scale of the axes. **(C)** The number of genes highly expressed in CR male compared to AL male (*P*adj<0.01, left circle) and CR female compared to AL female (*P*adj<0.01, right circle), with the overlap representing the 316 CR-induced genes common to both sexes. GO analysis (KEGG_2019_Mouse) were conducted with the overlapping gene list and the top 5 GO pathways are shown in the table. **(D)** A heatmap showing log_2_ Z-scores for genes linked to the GO terms of ‘biosynthesis of unsaturated fatty acids’ or ‘fatty acid biosynthesis’ among the 316 genes whose expression was higher in CR than in AL in both males and females (*P*adj<0.05). Data represent the following numbers of mice per group: *male AL,* n=6; *female AL,* n=5; *male CR,* n=5; *female CR,* n=6. For (A), R scripts are provided as a Source Data file.

**Supplementary Figure 12, Related to Figure 8 – CR and sex effects on the masses of adipose and non-adipose tissues in aged mice.**

Male and female mice were fed AL or CR diets from 78-84 weeks of age, as described for Figure 7. After six weeks of CR (84 weeks of age) tissues were sampled and weighed. **(A-C)** Violin plots, overlaid with individual data points, showing the absolute mass (A,B) or % body mass (C) of each tissue; Gastroc = gastrocnemius muscle. The numbers of mice per group and tissue type are as reported for Figure 7G. For each tissue, significant effects of diet and/or sex were determined by 2-way ANOVA with Tukey’s multiple comparisons test and are reported as described for Figure 8. Source data are provided as a Source Data file.

**Supplementary Figure 13, Related to Figure 10 – Baseline fat mass or BMI do not influence sex differences in the effects of CR on body mass or body composition.**

Twenty male and twenty-two female volunteers participated in a weight loss study involving a 4-week dietary intervention, as described for Figure 10. **(A-F)** Simple linear regression of baseline fat mass (A-C) or baseline BMI (D-F) vs fold-change (*week 4 vs week 0)* in body mass (A,D), fat mass (B,E) and fat-free mass (C,F). For each sex, significant associations between age and outcome (fold-change) are indicated beneath each graph as ‘*P,* Slope’. ANCOVA was further used to test if the age-outcome relationship differs significantly between males and females. ANCOVA results are reported beneath each graph as ‘*P,* Slope’ and ‘*P,* Intercept’ for males vs females (M vs F). Similar slopes but different intercepts show that sex significantly influences weight loss, but the influence of baseline fat mass or BMI does not differ between the sexes. See also Table 3. Source data are provided as a Source Data file.

**Supplementary Figure 14, Related to Figures 2, 8 and 10 – CR-induced fat loss in mice is age- and sex-dependent.**

Male and female mice were fed AL or CR diets from 9-15 or 78-84 weeks of age and fat mass was determined each week, as described for Figures 2 and 7. For each mouse (CR and AL diets) the ratio of absolute fat masses (g) was calculated for week 4 vs week 0 (i.e. 13 vs 9 weeks of age for young mice; 82 vs 78 weeks for aged mice); week 4 was selected for consistency with the 4-week duration of our human CR study (Fig. 9). For the CR mice, the fold-change in fat mass, relative to AL mice, was then determined as follows: i) the mean week 4:week 0 ratio was calculated for AL mice in each age group; ii) within each sex and age group, the week 4:week 0 ratio for each CR mouse was calculated relative to the mean week 4:week 0 ratio for the corresponding AL group. The rationale for showing these “fold changes of fold changes” is that the AL mice gain fat mass over the course of the study and therefore we must determine how much the fat mass of each CR mouse has changed compared to where we would expect it to be if it had continued on AL diet. Data in are shown as violin plots of the following numbers of mice per group: *young female,* n=31; *young male,* n=26; *old female,* n=11; *old male,* n=9. Significant effects of diet and/or sex were determined by 2-way ANOVA with Tukey’s multiple comparisons test and are reported as described for Figure S3. Source data are provided as a Source Data file.

**Supplementary Figure 15, Related to Figure 10 – Body fat percentage in human CR participants.**

Twenty male and twenty-two female volunteers participated in a weight loss study involving a 4-week dietary intervention, as described for Figure 10. Total % fat mass at week 0 is shown for males vs females separated into younger (<45 years) and older (>45 years) groups. Data are presented as violin plots overlaid with individual data points. Significant effects of age, sex, and age*sex interaction were assessed using 2-way ANOVA with Tukey’s multiple comparisons test. Overall *P* values for each variable, and their interactions, are shown beneath the graph. Significant differences between comparable groups are indicated by ** (*P*<0.01) or *** (*P*<0.001). See also Table 3. Source data are provided as a Source Data file.

## REFERENCES

1. Lin SJ, Kaeberlein M, Andalis AA, Sturtz LA, Defossez PA, Culotta VC, et al. Calorie restriction extends Saccharomyces cerevisiae lifespan by increasing respiration. Nature. 2002;418(6895):344–8.

2. Weindruch R, Walford RL, Fligiel S, and Guthrie D. The retardation of aging in mice by dietary restriction: longevity, cancer, immunity and lifetime energy intake. J Nutr. 1986;116(4):641–54.

3. Mattison JA, Colman RJ, Beasley TM, Allison DB, Kemnitz JW, Roth GS, et al. Caloric restriction improves health and survival of rhesus monkeys. Nat Commun. 2017;8:14063.

4. Villareal DT, Fontana L, Das SK, Redman L, Smith SR, Saltzman E, et al. Effect of Two-Year Caloric Restriction on Bone Metabolism and Bone Mineral Density in Non-Obese Younger Adults: A Randomized Clinical Trial. J Bone Miner Res. 2015;doi: 10.1002/jbmr.2701. [Epub ahead of print].

5. Speakman JR, and Mitchell SE. Caloric restriction. Mol Aspects Med. 2011;32(3):159–221.

6. Most J, Tosti V, Redman LM, and Fontana L. Calorie restriction in humans: An update. Ageing Res Rev. 2017;39:36–45.

7. Barzilai N, Banerjee S, Hawkins M, Chen W, and Rossetti L. Caloric restriction reverses hepatic insulin resistance in aging rats by decreasing visceral fat. J Clin Invest. 1998;101(7):1353–61.

8. López-Otín C, Galluzzi L, Freije JMP, Madeo F, and Kroemer G. Metabolic Control of Longevity. Cell. 2016;166(4):802–21.

9. Mancuso P, and Bouchard B. The Impact of Aging on Adipose Function and Adipokine Synthesis. Front Endocrinol (Lausanne). 2019;10:137-.

10. Gabriely I, Ma XH, Yang XM, Atzmon G, Rajala MW, Berg AH, et al. Removal of visceral fat prevents insulin resistance and glucose intolerance of aging: an adipokine-mediated process? Diabetes. 2002;51(10):2951–8.

11. Muzumdar R, Allison DB, Huffman DM, Ma X, Atzmon G, Einstein FH, et al. Visceral adipose tissue modulates mammalian longevity. Aging cell. 2008;7(3):438–40.

12. Tran TT, Yamamoto Y, Gesta S, and Kahn CR. Beneficial effects of subcutaneous fat transplantation on metabolism. Cell Metab. 2008;7(5):410–20.

13. Larson-Meyer DE, Heilbronn LK, Redman LM, Newcomer BR, Frisard MI, Anton S, et al. Effect of calorie restriction with or without exercise on insulin sensitivity, beta-cell function, fat cell size, and ectopic lipid in overweight subjects. Diabetes Care. 2006;29(6):1337–44.

14. Mauvais-Jarvis F. Gender differences in glucose homeostasis and diabetes. Physiol Behav. 2017.

15. Oliva M, Muñoz-Aguirre M, Kim-Hellmuth S, Wucher V, Gewirtz ADH, Cotter DJ, et al. The impact of sex on gene expression across human tissues. Science. 2020;369(6509):eaba3066.

16. Valencak TG, Osterrieder A, and Schulz TJ. Sex matters: The effects of biological sex on adipose tissue biology and energy metabolism. Redox Biology. 2017;12:806–13.

17. Maggi A, and Della Torre S. Sex, metabolism and health. Mol Metabol. 2018;15:3–7.

18. Prendergast BJ, Onishi KG, and Zucker I. Female mice liberated for inclusion in neuroscience and biomedical research. Neurosci Biobehav Rev. 2014;40:1–5.

19. Kane AE, Sinclair DA, Mitchell JR, and Mitchell SJ. Sex differences in the response to dietary restriction in rodents. Current Opinion in Physiology. 2018;6:28–34.

20. Redman LM, Heilbronn LK, Martin CK, Alfonso A, Smith SR, Ravussin E, et al. Effect of calorie restriction with or without exercise on body composition and fat distribution. J Clin Endocrinol Metab. 2007;92(3):865–72.

21. Shi H, Strader AD, Woods SC, and Seeley RJ. Sexually dimorphic responses to fat loss after caloric restriction or surgical lipectomy. Am J Physiol Endocrinol Metab. 2007;293(1):E316–26.

22. Cawthorn WP, Scheller EL, Parlee SD, Pham HA, Learman BS, Redshaw CM, et al. Expansion of Bone Marrow Adipose Tissue During Caloric Restriction Is Associated With Increased Circulating Glucocorticoids and Not With Hypoleptinemia. Endocrinology. 2016;157(2):508–21.

23. Della Torre S, Mitro N, Meda C, Lolli F, Pedretti S, Barcella M, et al. Short-Term Fasting Reveals Amino Acid Metabolism as a Major Sex-Discriminating Factor in the Liver. Cell Metab. 2018;28(2):256–67 e5.

24. Lee SK. Sex as an important biological variable in biomedical research. BMB reports. 2018;51(4):167–73.

25. Johnson J, Sharman Z, Vissandjée B, and Stewart DE. Does a change in health research funding policy related to the integration of sex and gender have an impact? PLoS One. 2014;9(6):e99900-e.

26. Bruss MD, Khambatta CF, Ruby MA, Aggarwal I, and Hellerstein MK. Calorie restriction increases fatty acid synthesis and whole body fat oxidation rates. Am J Physiol Endocrinol Metab. 2010;298(1):E108–16.

27. Matthews DR, Hosker JP, Rudenski AS, Naylor BA, Treacher DF, and Turner RC. Homeostasis model assessment: insulin resistance and beta-cell function from fasting plasma glucose and insulin concentrations in man. Diabetologia. 1985;28(7):412–9.

28. Matsuda M, and DeFronzo RA. Insulin sensitivity indices obtained from oral glucose tolerance testing: comparison with the euglycemic insulin clamp. Diabetes Care. 1999;22(9):1462.

29. Zhang L, Xue X, Zhai R, Yang X, Li H, Zhao L, et al. Timing of Calorie Restriction in Mice Impacts Host Metabolic Phenotype with Correlative Changes in Gut Microbiota. mSystems. 2019;4(6):e00348–19.

30. Bartelt A, Koehne T, Todter K, Reimer R, Muller B, Behler-Janbeck F, et al. Quantification of Bone Fatty Acid Metabolism and Its Regulation by Adipocyte Lipoprotein Lipase. Int J Mol Sci. 2017;18(6).

31. Suchacki KJ, Tavares AAS, Mattiucci D, Scheller EL, Papanastasiou G, Gray C, et al. Bone marrow adipose tissue is a unique adipose subtype with distinct roles in glucose homeostasis. Nat Commun. 2020;11(1):3097.

32. Petersen MC, and Shulman GI. Roles of Diacylglycerols and Ceramides in Hepatic Insulin Resistance. Trends Pharmacol Sci. 2017;38(7):649–65.

33. Chaurasia B, Tippetts TS, Mayoral Monibas R, Liu J, Li Y, Wang L, et al. Targeting a ceramide double bond improves insulin resistance and hepatic steatosis. Science. 2019;365(6451):386–92.

34. Apostolopoulou M, Gordillo R, Gancheva S, Strassburger K, Herder C, Esposito I, et al. Role of ceramide-to-dihydroceramide ratios for insulin resistance and non-alcoholic fatty liver disease in humans. BMJ Open Diabetes Research & Care. 2020;8(2):e001860.

35. Perry RJ, Peng L, Cline GW, Petersen KF, and Shulman GI. A Non-invasive Method to Assess Hepatic Acetyl-CoA In Vivo. Cell Metab. 2017;25(3):749–56.

36. Forney LA, Stone KP, Gibson AN, Vick AM, Sims LC, Fang H, et al. Sexually Dimorphic Effects of Dietary Methionine Restriction are Dependent on Age when the Diet is Introduced. Obesity. 2020;28(3):581–9.

37. Chaix A, Deota S, Bhardwaj R, Lin T, and Panda S. Sex- and age-dependent outcomes of 9-hour time-restricted feeding of a Western high-fat high-sucrose diet in C57BL/6J mice. Cell reports. 2021;36(7).

38. Gee DM, Flurkey K, and Finch CE. Aging and the Regulation of Luteinizing Hormone in C57BL/6J Mice: Impaired Elevations after Ovariectomy and Spontaneous Elevations at Advanced Ages1. Biol Reprod. 1983;28(3):598–607.

39. Diaz Brinton R. Minireview: translational animal models of human menopause: challenges and emerging opportunities. Endocrinology. 2012;153(8):3571–8.

40. Piotrowska K, Tarnowski M, Zgutka K, and Pawlik A. Gender Differences in Response to Prolonged Every-Other-Day Feeding on the Proliferation and Apoptosis of Hepatocytes in Mice. Nutrients. 2016;8(3):176-.

41. Musante GJ. The dietary rehabilitation clinic: Evaluative report of a behavioral and dietary treatment of obesity. Behav Ther. 1976;7(2):198–204.

42. O’Neil PM, Currey HS, Hirsch AA, Riddle FE, Taylor CI, Malcolm RJ, et al. Effects of sex of subject and spouse involvement on weight loss in a behavioral treatment program: a retrospective investigation. Addict Behav. 1979;4(2):167–77.

43. Mauriege P, Imbeault P, Langin D, Lacaille M, Almeras N, Tremblay A, et al. Regional and gender variations in adipose tissue lipolysis in response to weight loss. J Lipid Res. 1999;40(9):1559–71.

44. Wong MHT, Holst C, Astrup A, Handjieva-Darlenska T, Jebb SA, Kafatos A, et al. Caloric restriction induces changes in insulin and body weight measurements that are inversely associated with subsequent weight regain. PLoS One. 2012;7(8):e42858-e.

45. Azar M, Nikpay M, Harper M-E, McPherson R, and Dent R. Can response to dietary restriction predict weight loss after Roux-en-Y gastroplasty? Obesity. 2016;24(4):805–11.

46. Das SK, Roberts SB, Bhapkar MV, Villareal DT, Fontana L, Martin CK, et al. Body-composition changes in the Comprehensive Assessment of Long-term Effects of Reducing Intake of Energy (CALERIE)-2 study: a 2-y randomized controlled trial of calorie restriction in nonobese humans. The American Journal of Clinical Nutrition. 2017;105(4):913–27.

47. Cawthorn WP, Scheller EL, Learman BS, Parlee SD, Simon BR, Mori H, et al. Bone Marrow Adipose Tissue Is an Endocrine Organ that Contributes to Increased Circulating Adiponectin during Caloric Restriction. Cell Metab. 2014;20(2):368–75.

48. Dionne DA, Skovso S, Templeman NM, Clee SM, and Johnson JD. Caloric Restriction Paradoxically Increases Adiposity in Mice With Genetically Reduced Insulin. Endocrinology. 2016;157(7):2724–34.

49. Li X, Cope MB, Johnson MS, Smith DL, Jr., and Nagy TR. Mild calorie restriction induces fat accumulation in female C57BL/6J mice. Obesity. 2010;18(3):456–62.

50. Leenen R, van der Kooy K, Droop A, Seidell JC, Deurenberg P, Weststrate JA, et al. Visceral fat loss measured by magnetic resonance imaging in relation to changes in serum lipid levels of obese men and women. Arterioscler Thromb. 1993;13(4):487–94.

51. Fontana L, Klein S, and Holloszy JO. Effects of long-term calorie restriction and endurance exercise on glucose tolerance, insulin action, and adipokine production. Age. 2010;32(1):97–108.

52. Evans EM, Mojtahedi MC, Thorpe MP, Valentine RJ, Kris-Etherton PM, and Layman DK. Effects of protein intake and gender on body composition changes: a randomized clinical weight loss trial. Nutr Metab (Lond). 2012;9(1):55.

53. Kraus WE, Bhapkar M, Huffman KM, Pieper CF, Krupa Das S, Redman LM, et al. 2 years of calorie restriction and cardiometabolic risk (CALERIE): exploratory outcomes of a multicentre, phase 2, randomised controlled trial. The Lancet Diabetes & Endocrinology. 2019;7(9):673–83.

54. Dhahbi JM, Kim HJ, Mote PL, Beaver RJ, and Spindler SR. Temporal linkage between the phenotypic and genomic responses to caloric restriction. Proc Natl Acad Sci U S A. 2004;101(15):5524–9.

55. Sheng Y, Xia F, Chen L, Lv Y, Lv S, Yu J, et al. Differential Responses of White Adipose Tissue and Brown Adipose Tissue to Calorie Restriction During Aging. The Journals of Gerontology: Series A. 2020;76(3):393–9.

56. Rohrbach S, Aurich AC, Li L, and Niemann B. Age-associated loss in adiponectin-activation by caloric restriction: lack of compensation by enhanced inducibility of adiponectin paralogs CTRP2 and CTRP7. Mol Cell Endocrinol. 2007;277(1-2):26–34.

57. Mitchell SJ, Madrigal-Matute J, Scheibye-Knudsen M, Fang E, Aon M, González-Reyes JA, et al. Effects of Sex, Strain, and Energy Intake on Hallmarks of Aging in Mice. Cell Metab. 2016;23(6):1093–112.

58. Brandhorst S, Choi In Y, Wei M, Cheng Chia W, Sedrakyan S, Navarrete G, et al. A Periodic Diet that Mimics Fasting Promotes Multi-System Regeneration, Enhanced Cognitive Performance, and Healthspan. Cell Metab. 2015;22(1):86–99.

59. Porter MH, Fine JB, Cutchins AG, Bai Y, and DiGirolamo M. Sexual dimorphism in the response of adipose mass and cellularity to graded caloric restriction. Obes Res. 2004;12(1):131–40.

60. Valle A, Catala-Niell A, Colom B, Garcia-Palmer FJ, Oliver J, and Roca P. Sex-related differences in energy balance in response to caloric restriction. Am J Physiol Endocrinol Metab. 2005;289(1):E15–22.

61. Valle A, Garcia-Palmer FJ, Oliver J, and Roca P. Sex differences in brown adipose tissue thermogenic features during caloric restriction. Cellular physiology and biochemistry : international journal of experimental cellular physiology, biochemistry, and pharmacology. 2007;19(1-4):195–204.

62. Hill JO, Talano CM, Nickel M, and DiGirolamo M. Energy utilization in food-restricted female rats. J Nutr. 1986;116(10):2000–12.

63. Hill JO, Latiff A, and DiGirolamo M. Effects of variable caloric restriction on utilization of ingested energy in rats. Am J Physiol. 1985;248(5 Pt 2):R549–59.

64. Fenton JI, Nunez NP, Yakar S, Perkins SN, Hord NG, and Hursting SD. Diet-induced adiposity alters the serum profile of inflammation in C57BL/6N mice as measured by antibody array. Diabetes, obesity & metabolism. 2009;11(4):343–54.

65. Varady KA, Allister CA, Roohk DJ, and Hellerstein MK. Improvements in body fat distribution and circulating adiponectin by alternate-day fasting versus calorie restriction. The Journal of Nutritional Biochemistry. 2010;21(3):188–95.

66. Harvey AE, Lashinger LM, Otto G, Nunez NP, and Hursting SD. Decreased systemic IGF-1 in response to calorie restriction modulates murine tumor cell growth, nuclear factor-kappaB activation, and inflammation-related gene expression. Mol Carcinog. 2013;52(12):997–1006.

67. Zgheib S, Mequinion M, Lucas S, Leterme D, Ghali O, Tolle V, et al. Long-term physiological alterations and recovery in a mouse model of separation associated with time-restricted feeding: a tool to study anorexia nervosa related consequences. PLoS One. 2014;9(8):e103775.

68. Janssen I, and Ross R. Effects of sex on the change in visceral, subcutaneous adipose tissue and skeletal muscle in response to weight loss. Int J Obes Relat Metab Disord. 1999;23(10):1035–46.

69. Wirth A, and Steinmetz B. Gender differences in changes in subcutaneous and intra-abdominal fat during weight reduction: an ultrasound study. Obes Res. 1998;6(6):393–9.

70. Parker B, Noakes M, Luscombe N, and Clifton P. Effect of a high-protein, high-monounsaturated fat weight loss diet on glycemic control and lipid levels in type 2 diabetes. Diabetes Care. 2002;25(3):425–30.

71. Weiss EP, Racette SB, Villareal DT, Fontana L, Steger-May K, Schechtman KB, et al. Improvements in glucose tolerance and insulin action induced by increasing energy expenditure or decreasing energy intake: a randomized controlled trial. Am J Clin Nutr. 2006;84(5):1033–42.

72. Chen JH, Ouyang C, Ding Q, Song J, Cao W, and Mao L. A Moderate Low-Carbohydrate Low-Calorie Diet Improves Lipid Profile, Insulin Sensitivity and Adiponectin Expression in Rats. Nutrients. 2015;7(6):4724–38.

73. Zhu M, Miura J, Lu LX, Bernier M, DeCabo R, Lane MA, et al. Circulating adiponectin levels increase in rats on caloric restriction: the potential for insulin sensitization. Exp Gerontol. 2004;39(7):1049–59.

74. Mitchell SE, Tang Z, Kerbois C, Delville C, Konstantopedos P, Bruel A, et al. The effects of graded levels of calorie restriction: I. impact of short term calorie and protein restriction on body composition in the C57BL/6 mouse. Oncotarget. 2015;6(18):15902–30.

75. Mitchell SE, Delville C, Konstantopedos P, Hurst J, Derous D, Green C, et al. The effects of graded levels of calorie restriction: II. Impact of short term calorie and protein restriction on circulating hormone levels, glucose homeostasis and oxidative stress in male C57BL/6 mice. Oncotarget. 2015;6(27):23213–37.

76. Mitchell SE, Tang Z, Kerbois C, Delville C, Derous D, Green CL, et al. The effects of graded levels of calorie restriction: VIII. Impact of short term calorie and protein restriction on basal metabolic rate in the C57BL/6 mouse. Oncotarget. 2017;8(11):17453–74.

77. Derous D, Mitchell SE, Green CL, Wang Y, Han JDJ, Chen L, et al. The Effects of Graded Levels of Calorie Restriction: X. Transcriptomic Responses of Epididymal Adipose Tissue. J Gerontol A Biol Sci Med Sci. 2017.

78. Ballor DL, and Poehlman ET. Exercise-training enhances fat-free mass preservation during diet-induced weight loss: a meta-analytical finding. Int J Obes Relat Metab Disord. 1994;18(1):35–40.

79. Suchacki KJ, Thomas BJ, Fyfe C, Tavares AAS, Sulston RJ, Lovdel A, et al. The effects of caloric restriction on adipose tissue and metabolic health are sex- and age-dependent. bioRxiv. 2022:2022.02.20.481222.

80. Arner P, Andersson DP, Backdahl J, Dahlman I, and Ryden M. Weight Gain and Impaired Glucose Metabolism in Women Are Predicted by Inefficient Subcutaneous Fat Cell Lipolysis. Cell Metab. 2018;28(1):45–54 e3.

81. Gazdag AC, Dumke CL, Kahn CR, and Cartee GD. Calorie restriction increases insulin-stimulated glucose transport in skeletal muscle from IRS-1 knockout mice. Diabetes. 1999;48(10):1930–6.

82. Johnson ML, Distelmaier K, Lanza IR, Irving BA, Robinson MM, Konopka AR, et al. Mechanism by Which Caloric Restriction Improves Insulin Sensitivity in Sedentary Obese Adults. Diabetes. 2016;65(1):74–84.

83. Sequea DA, Sharma N, Arias EB, and Cartee GD. Calorie restriction enhances insulin-stimulated glucose uptake and Akt phosphorylation in both fast-twitch and slow-twitch skeletal muscle of 24-month-old rats. J Gerontol A Biol Sci Med Sci. 2012;67(12):1279–85.

84. Liu Y, Basty N, Whitcher B, Bell JD, Sorokin EP, van Bruggen N, et al. Genetic architecture of 11 organ traits derived from abdominal MRI using deep learning. eLife. 2021;10:e65554.

85. Perry RJ, Peng L, Cline GW, Wang Y, Rabin-Court A, Song JD, et al. Mechanisms by which a Very-Low-Calorie Diet Reverses Hyperglycemia in a Rat Model of Type 2 Diabetes. Cell Metab. 2018;27(1):210–7.e3.

86. Taylor R, Al-Mrabeh A, Zhyzhneuskaya S, Peters C, Barnes AC, Aribisala BS, et al. Remission of Human Type 2 Diabetes Requires Decrease in Liver and Pancreas Fat Content but Is Dependent upon Capacity for β Cell Recovery. Cell Metab. 2018;28(4):547–56.e3.

87. Norheim F, Bjellaas T, Hui ST, Chella Krishnan K, Lee J, Gupta S, et al. Genetic, dietary, and sex-specific regulation of hepatic ceramides and the relationship between hepatic ceramides and IR [S]. J Lipid Res. 2018;59(7):1164–74.

88. Turpin SM, Nicholls HT, Willmes DM, Mourier A, Brodesser S, Wunderlich CM, et al. Obesity-induced CerS6-dependent C16:0 ceramide production promotes weight gain and glucose intolerance. Cell Metab. 2014;20(4):678–86.

89. Raichur S, Wang ST, Chan PW, Li Y, Ching J, Chaurasia B, et al. CerS2 haploinsufficiency inhibits β-oxidation and confers susceptibility to diet-induced steatohepatitis and insulin resistance. Cell Metab. 2014;20(4):687–95.

90. Raichur S, Brunner B, Bielohuby M, Hansen G, Pfenninger A, Wang B, et al. The role of C16:0 ceramide in the development of obesity and type 2 diabetes: CerS6 inhibition as a novel therapeutic approach. Molecular metabolism. 2019;21:36–50.

91. Holland WL, Brozinick JT, Wang LP, Hawkins ED, Sargent KM, Liu Y, et al. Inhibition of ceramide synthesis ameliorates glucocorticoid-, saturated-fat-, and obesity-induced insulin resistance. Cell Metab. 2007;5(3):167–79.

92. Zhang Q-J, Holland WL, Wilson L, Tanner JM, Kearns D, Cahoon JM, et al. Ceramide Mediates Vascular Dysfunction in Diet-Induced Obesity by PP2A-Mediated Dephosphorylation of the eNOS-Akt Complex. Diabetes. 2012;61(7):1848–59.

93. Mauvais-Jarvis F, Arnold AP, and Reue K. A Guide for the Design of Pre-clinical Studies on Sex Differences in Metabolism. Cell Metab. 2017;25(6):1216–30.

94. Link JC, Wiese CB, Chen X, Avetisyan R, Ronquillo E, Ma F, et al. X chromosome dosage of histone demethylase KDM5C determines sex differences in adiposity. The Journal of Clinical Investigation. 2020;130(11):5688–702.

95. Dearden L, Bouret SG, and Ozanne SE. Sex and gender differences in developmental programming of metabolism. Mol Metabol. 2018;15:8–19.

96. Shen T, Ding L, Ruan Y, Qin W, Lin Y, Xi C, et al. SIRT1 Functions as an Important Regulator of Estrogen-Mediated Cardiomyocyte Protection in Angiotensin II-Induced Heart Hypertrophy. Oxid Med Cell Longev. 2014;2014:713894.

97. Wei Y, and Huang J. Role of estrogen and its receptors mediated-autophagy in cell fate and human diseases. J Steroid Biochem Mol Biol. 2019;191:105380.

98. Gupte AA, Pownall HJ, and Hamilton DJ. Estrogen: An Emerging Regulator of Insulin Action and Mitochondrial Function. J Diabetes Res. 2015;2015:916585.

99. Klinge CM. Estrogenic control of mitochondrial function. Redox Biol. 2020;31:101435.

100. Wheatley KE, Nogueira LM, Perkins SN, and Hursting SD. Differential effects of calorie restriction and exercise on the adipose transcriptome in diet-induced obese mice. J Obes. 2011;2011:265417.

101. Dakin RS, Walker BR, Seckl JR, Hadoke PW, and Drake AJ. Estrogens protect male mice from obesity complications and influence glucocorticoid metabolism. Int J Obes (Lond). 2015;39(10):1539–47.

102. Zhu L, Martinez MN, Emfinger CH, Palmisano BT, and Stafford JM. Estrogen signaling prevents diet-induced hepatic insulin resistance in male mice with obesity. Am J Physiol Endocrinol Metab. 2014;306(10):E1188–E97.

103. Acosta-Rodríguez V, Rijo-Ferreira F, Izumo M, Xu P, Wight-Carter M, Green CB, et al. Circadian alignment of early onset caloric restriction promotes longevity in male C57BL/6J mice. Science. 2022;376(6598):1192–202.

104. Acosta-Rodriguez VA, de Groot MHM, Rijo-Ferreira F, Green CB, and Takahashi JS. Mice under Caloric Restriction Self-Impose a Temporal Restriction of Food Intake as Revealed by an Automated Feeder System. Cell Metab. 2017;26(1):267–77.e2.

105. Pak HH, Haws SA, Green CL, Koller M, Lavarias MT, Richardson NE, et al. Fasting drives the metabolic, molecular and geroprotective effects of a calorie-restricted diet in mice. Nature Metabolism. 2021;3(10):1327–41.

106. Mitchell SJ, Bernier M, Mattison JA, Aon MA, Kaiser TA, Anson RM, et al. Daily Fasting Improves Health and Survival in Male Mice Independent of Diet Composition and Calories. Cell Metab. 2019;29(1):221–8 e3.

107. Handelsman DJ, and Wartofsky L. Requirement for Mass Spectrometry Sex Steroid Assays in the Journal of Clinical Endocrinology and Metabolism. The Journal of Clinical Endocrinology & Metabolism. 2013;98(10):3971–3.

108. Kane AE, Hilmer SN, Boyer D, Gavin K, Nines D, Howlett SE, et al. Impact of Longevity Interventions on a Validated Mouse Clinical Frailty Index. The Journals of Gerontology: Series A. 2016;71(3):333–9.

109. Ferland G, Doucet I, and Mainville D. Phylloquinone and Menaquinone-4 Tissue Distribution at Different Life Stages in Male and Female Sprague-Dawley Rats Fed Different VK Levels Since Weaning or Subjected to a 40% Calorie Restriction since Adulthood. Nutrients. 2016;8(3):141.

110. Austad SN, and Bartke A. Sex Differences in Longevity and in Responses to Anti-Aging Interventions: A Mini-Review. Gerontology. 2016;62(1):40–6.

111. Bohler H, Jr., Mokshagundam S, and Winters SJ. Adipose tissue and reproduction in women. Fertil Steril. 2010;94(3):795–825.

112. Misra M, and Klibanski A. Endocrine consequences of anorexia nervosa. The lancet Diabetes & endocrinology. 2014;2(7):581–92.

113. Dobin A, Davis CA, Schlesinger F, Drenkow J, Zaleski C, Jha S, et al. STAR: ultrafast universal RNA-seq aligner. Bioinformatics. 2013;29(1):15–21.

114. Liao Y, Smyth GK, and Shi W. featureCounts: an efficient general purpose program for assigning sequence reads to genomic features. Bioinformatics. 2013;30(7):923–30.

115. Love MI, Huber W, and Anders S. Moderated estimation of fold change and dispersion for RNA-seq data with DESeq2. Genome Biol. 2014;15(12):550.

116. Mootha VK, Lindgren CM, Eriksson K-F, Subramanian A, Sihag S, Lehar J, et al. PGC-1α-responsive genes involved in oxidative phosphorylation are coordinately downregulated in human diabetes. Nat Genet. 2003;34(3):267–73.

117. Subramanian A, Tamayo P, Mootha VK, Mukherjee S, Ebert BL, Gillette MA, et al. Gene set enrichment analysis: a knowledge-based approach for interpreting genome-wide expression profiles. Proc Natl Acad Sci U S A. 2005;102(43):15545–50.

118. Xie Z, Bailey A, Kuleshov MV, Clarke DJB, Evangelista JE, Jenkins SL, et al. Gene Set Knowledge Discovery with Enrichr. Current Protocols. 2021;1(3):e90.

119. Kuleshov MV, Jones MR, Rouillard AD, Fernandez NF, Duan Q, Wang Z, et al. Enrichr: a comprehensive gene set enrichment analysis web server 2016 update. Nucleic Acids Res. 2016;44(W1):W90–7.

120. Chen EY, Tan CM, Kou Y, Duan Q, Wang Z, Meirelles GV, et al. Enrichr: interactive and collaborative HTML5 gene list enrichment analysis tool. BMC Bioinformatics. 2013;14:128.

121. Clarke DJB, Jeon M, Stein DJ, Moiseyev N, Kropiwnicki E, Dai C, et al. Appyters: Turning Jupyter Notebooks into data-driven web apps. Patterns. 2021;2(3).

122. Johnstone AM, Murison SD, Duncan JS, Rance KA, and Speakman JR. Factors influencing variation in basal metabolic rate include fat-free mass, fat mass, age, and circulating thyroxine but not sex, circulating leptin, or triiodothyronine. Am J Clin Nutr. 2005;82(5):941–8.

123. McCrory MA, Gomez TD, Bernauer EM, and Molé PA. Evaluation of a new air displacement plethysmograph for measuring human body composition. Med Sci Sports Exerc. 1995;27(12):1686–91.

124. Galarraga M, Campión J, Muñoz-Barrutia A, Boqué N, Moreno H, Martínez JA, et al. Adiposoft: automated software for the analysis of white adipose tissue cellularity in histological sections. J Lipid Res. 2012;53(12):2791–6.

125. Folch J, Lees M, and Sloane Stanley GH. A simple method for the isolation and purification of total lipides from animal tissues. J Biol Chem. 1957;226(1):497–509.

