## Supplementary_Figures for "The effects of caloric restriction on adipose tissue and metabolic health are sex- and age-dependent"

### Supplementary Figure 1

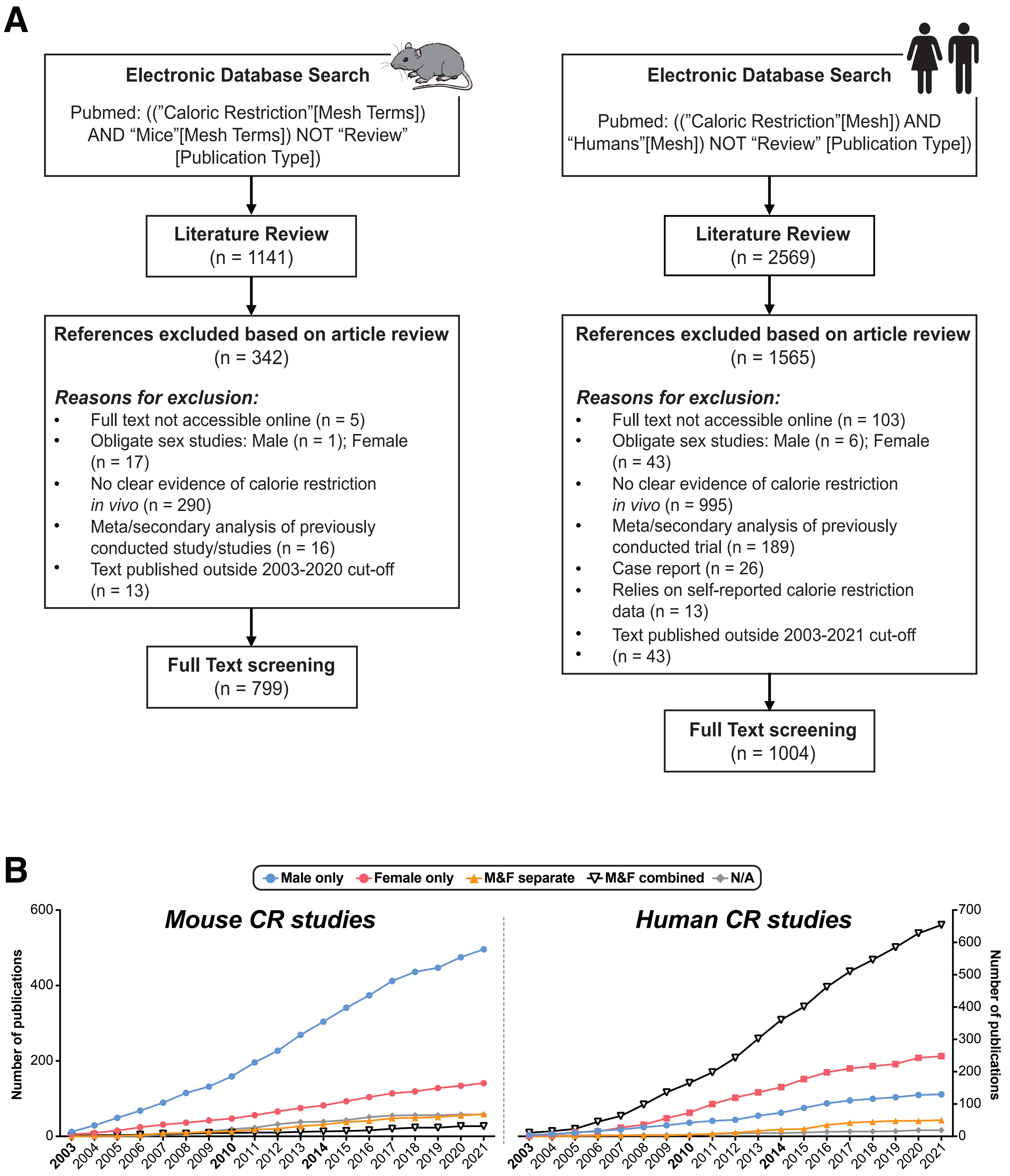

### Supplementary Figure 2

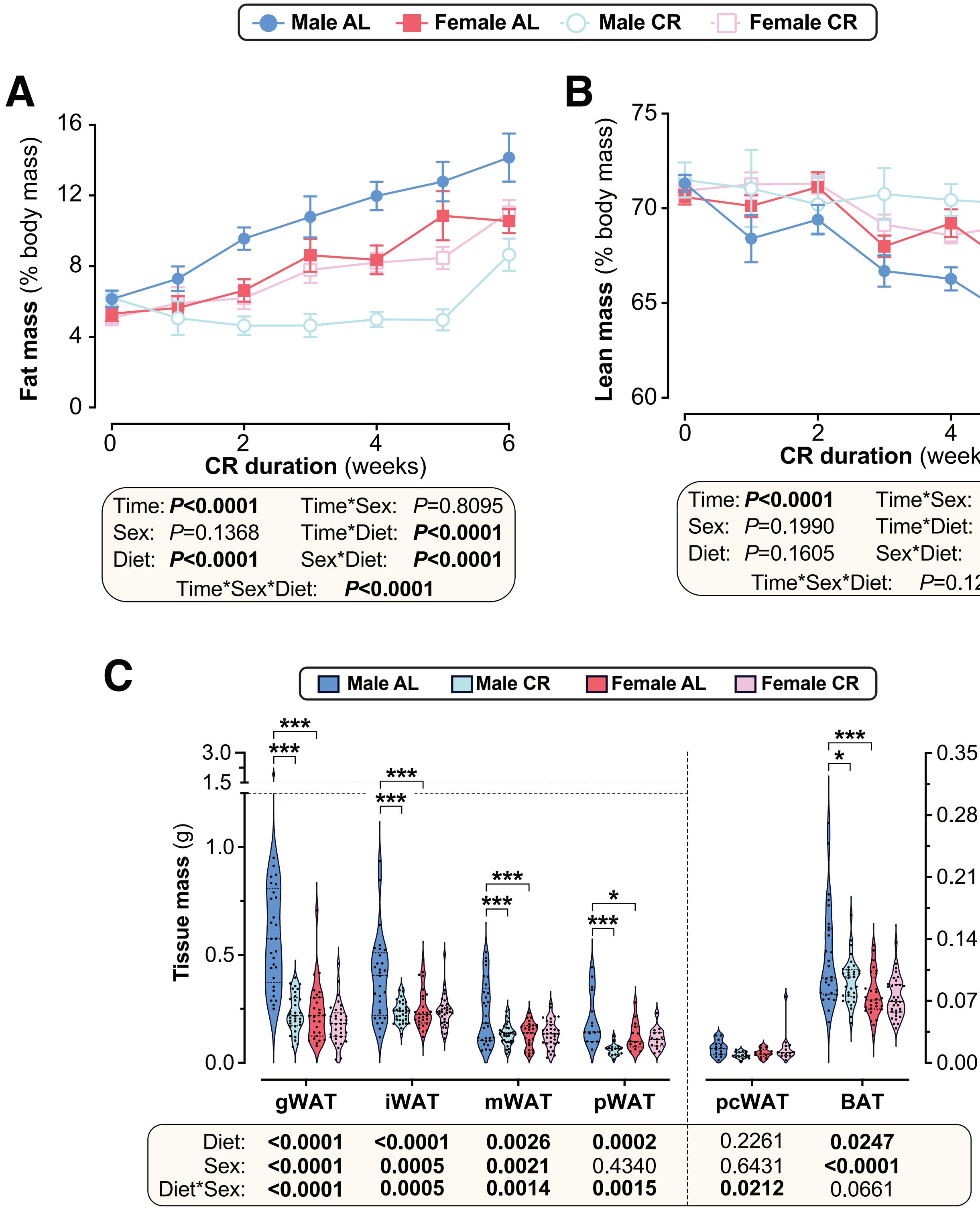

### Supplementary Figure 3

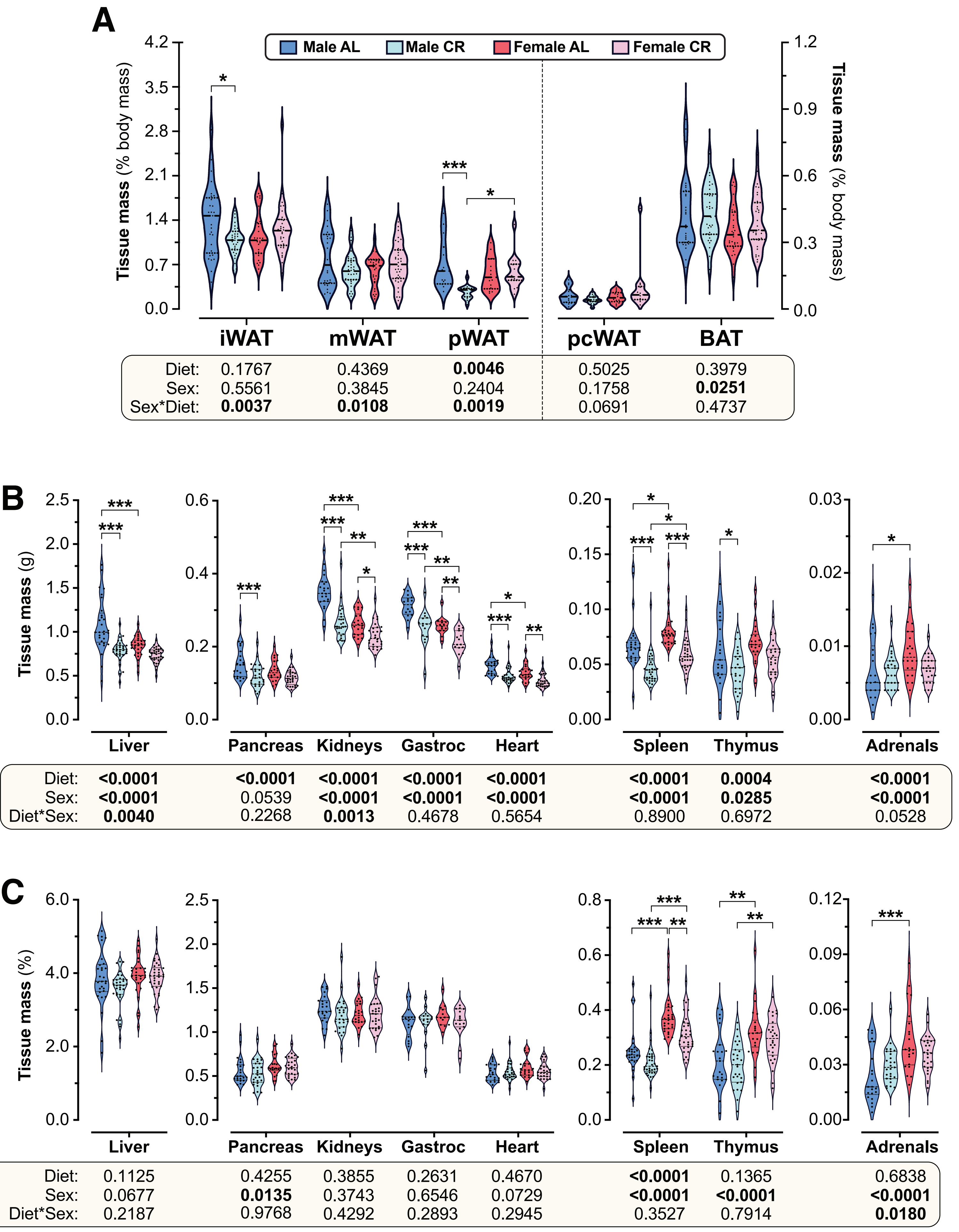


| | iWAT | mWAT | pWAT | pcWAT | BAT |
|---|---|---|---|---|---|
| Diet: | 0.1767 | 0.4369 | **0.0046** | 0.5025 | 0.3979 |
| Sex: | 0.5561 | 0.3845 | 0.2404 | 0.1758 | **0.0251** |
| Sex*Diet: | **0.0037** | **0.0108** | **0.0019** | 0.0691 | 0.4737 |

| | Liver | Pancreas | Kidneys | Gastroc | Heart | Spleen | Thymus | Adrenals |
|---|---|---|---|---|---|---|---|---|
| Diet: | **<0.0001** | **<0.0001** | **<0.0001** | **<0.0001** | **<0.0001** | **<0.0001** | **0.0004** | **<0.0001** |
| Sex: | **<0.0001** | 0.0539 | **<0.0001** | **<0.0001** | **<0.0001** | **<0.0001** | **0.0285** | **<0.0001** |
| Diet*Sex: | **0.0040** | 0.2268 | **0.0013** | 0.4678 | 0.5654 | 0.8900 | 0.6972 | 0.0528 |

| | Liver | Pancreas | Kidneys | Gastroc | Heart | Spleen | Thymus | Adrenals |
|---|---|---|---|---|---|---|---|---|
| Diet: | 0.1125 | 0.4255 | 0.3855 | 0.2631 | 0.4670 | **<0.0001** | 0.1365 | 0.6838 |
| Sex: | 0.0677 | **0.0135** | 0.3743 | 0.6546 | 0.0729 | **<0.0001** | **<0.0001** | **<0.0001** |
| Diet*Sex: | 0.2187 | 0.9768 | 0.4292 | 0.2893 | 0.2945 | 0.3527 | 0.7914 | **0.0180** |

### Supplementary Figure 4

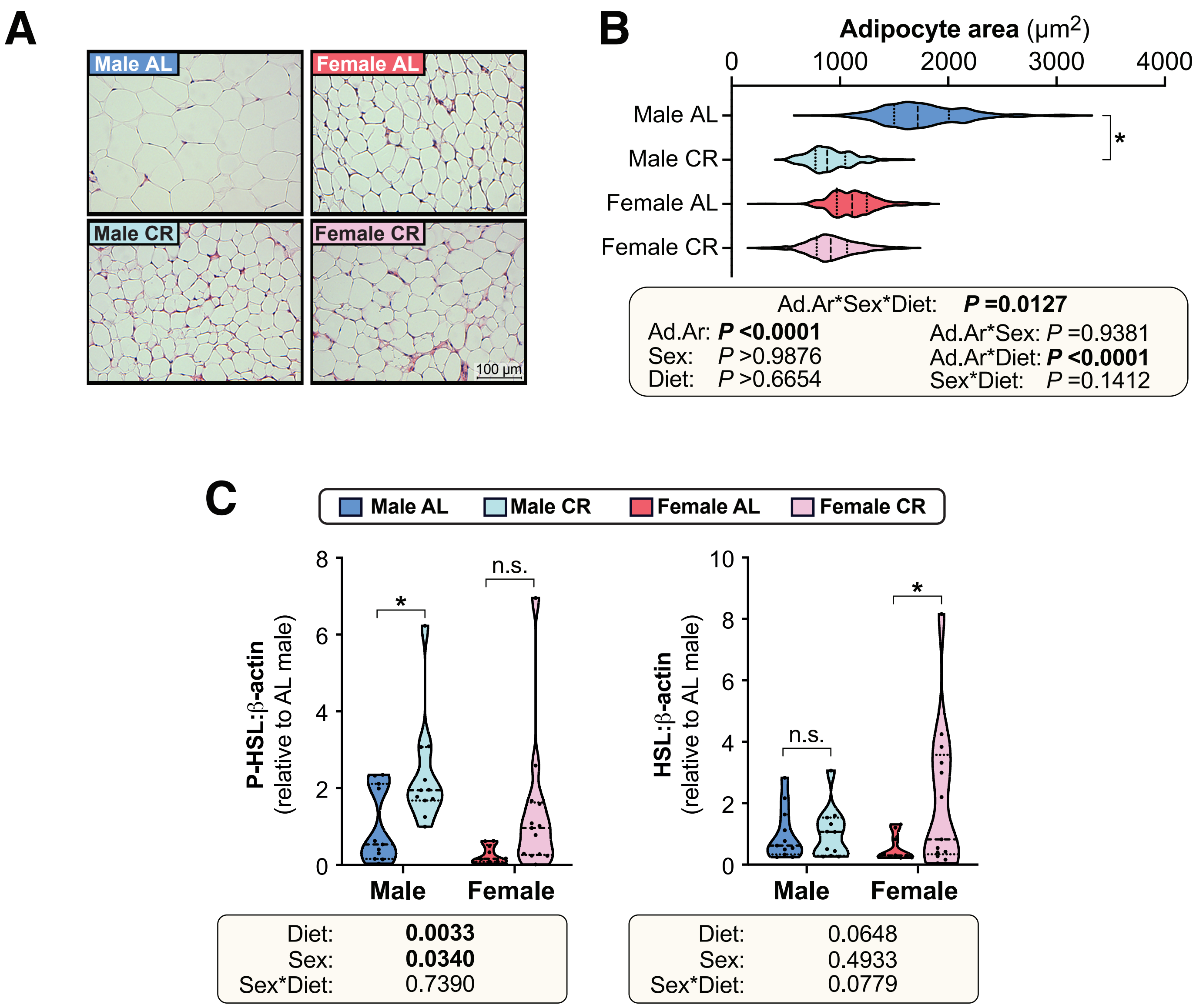

### Supplementary Figure 5

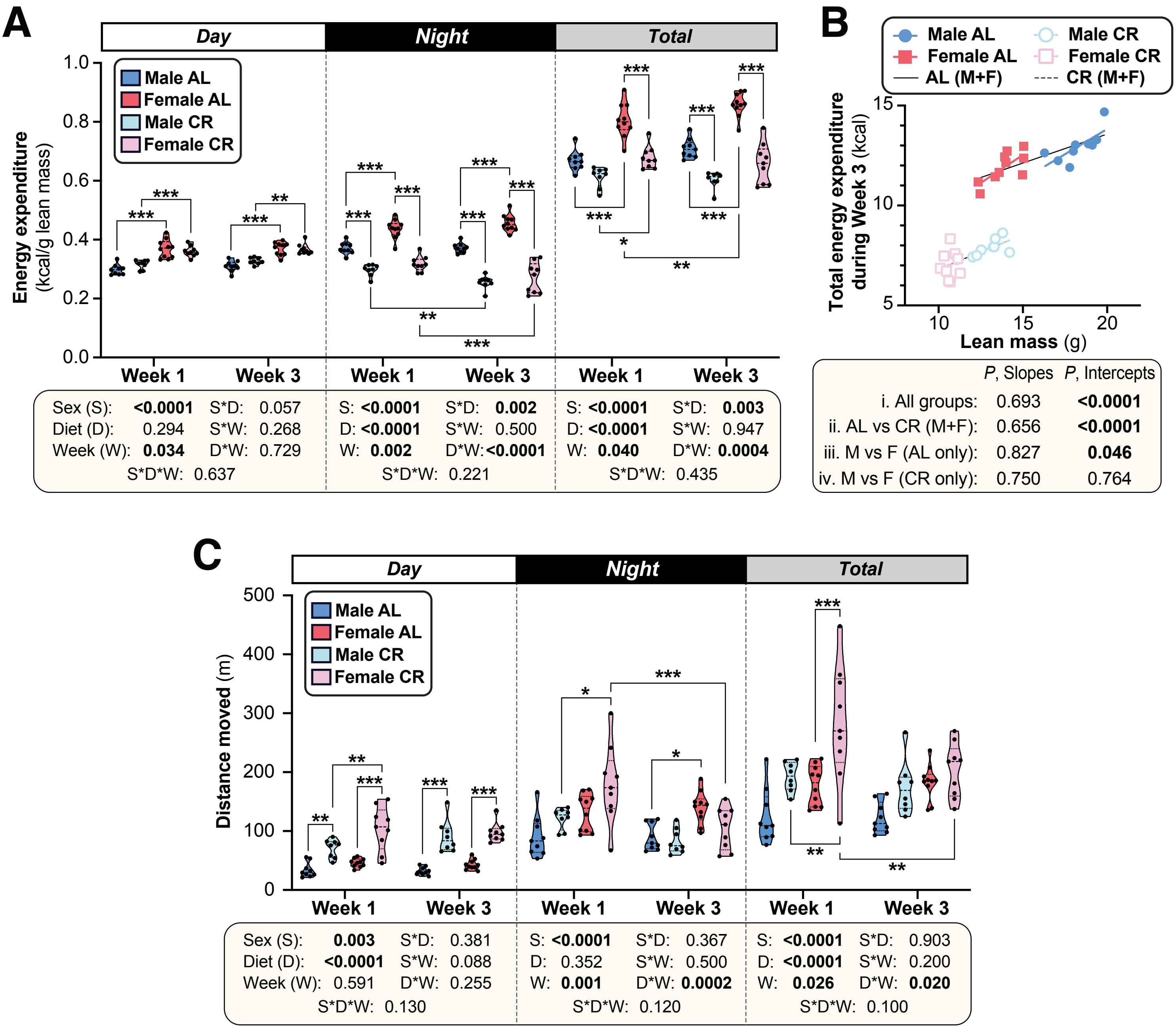

### Supplementary Figure 6

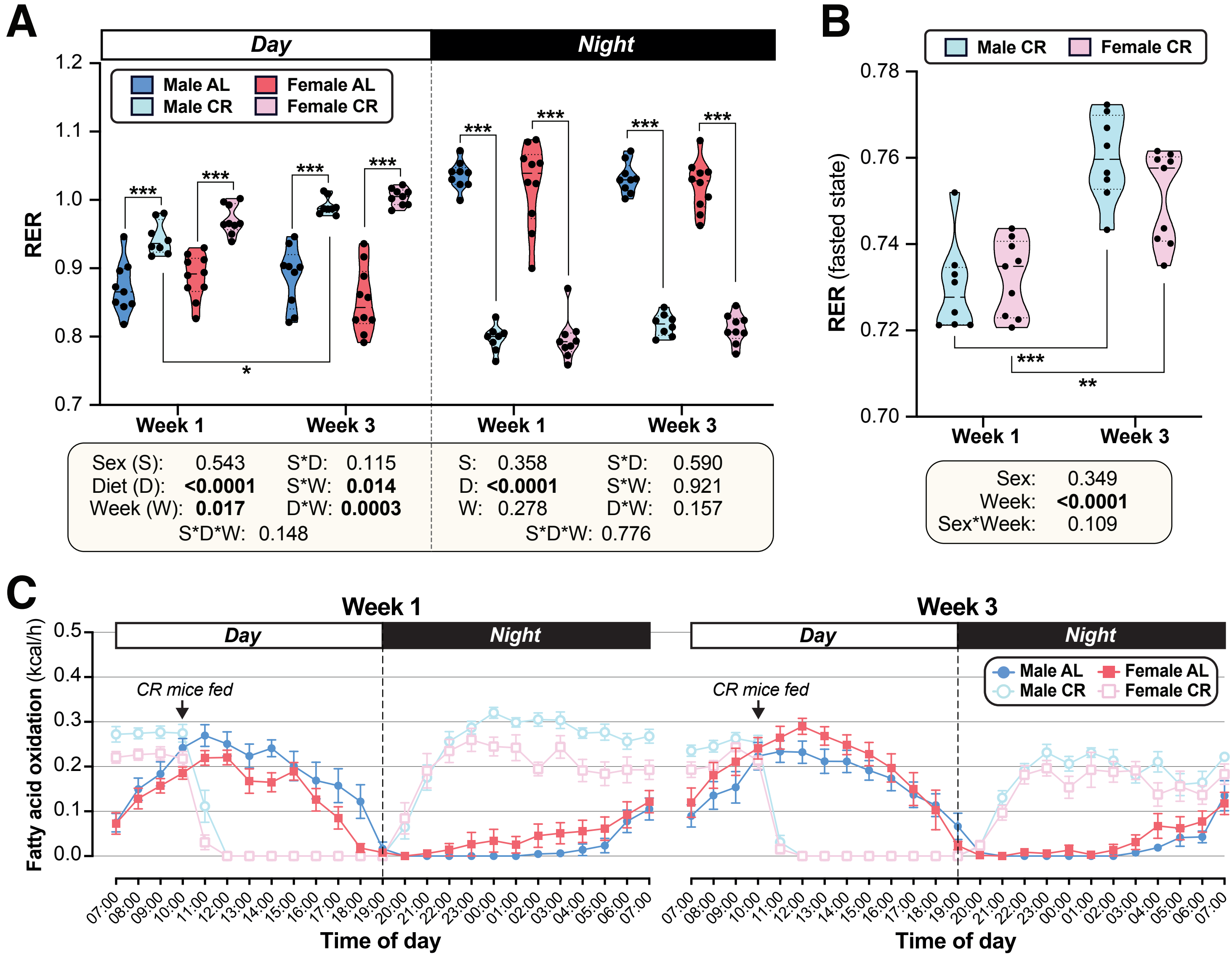

### Supplementary Figure 7

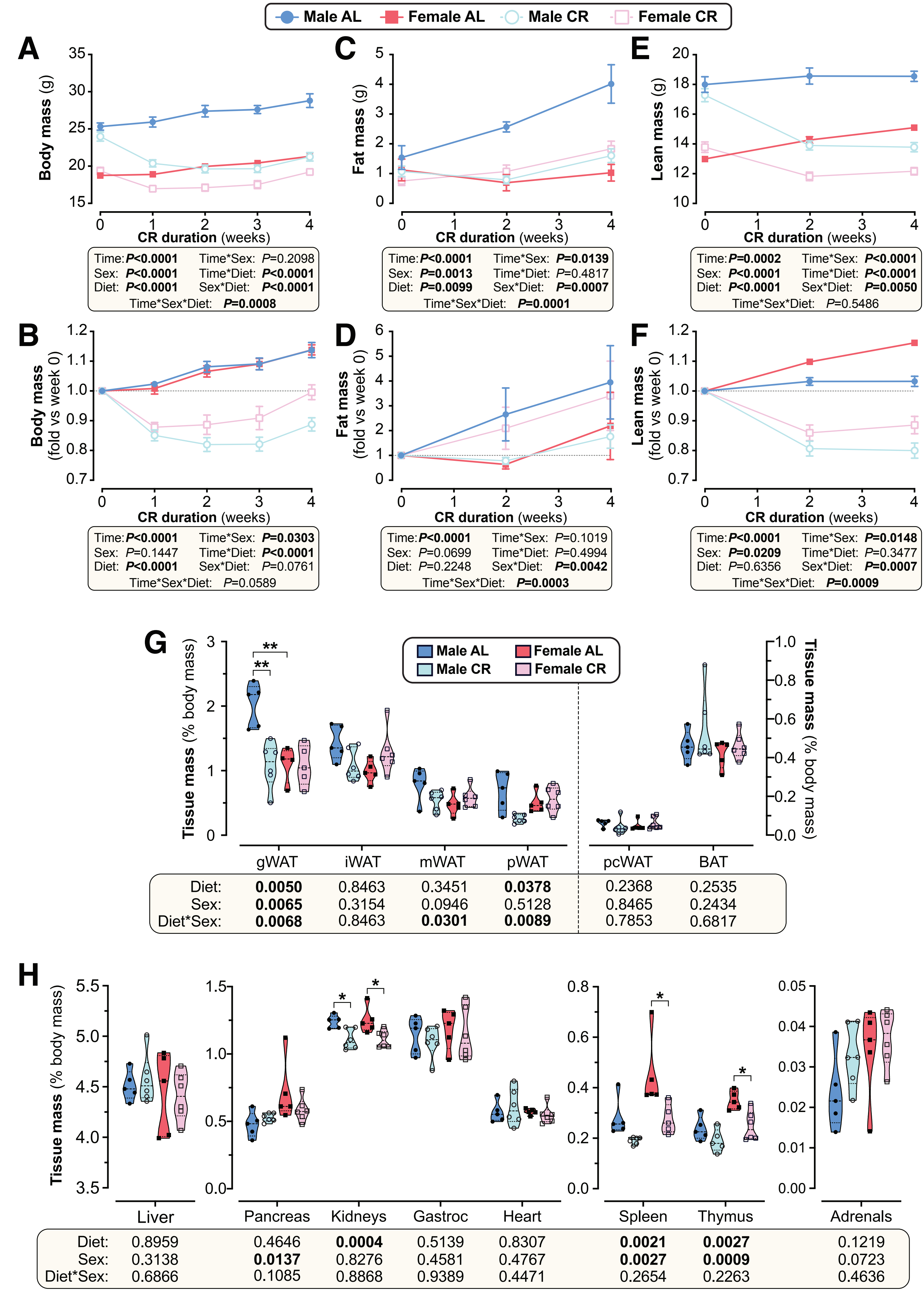

### Supplementary Figure 8

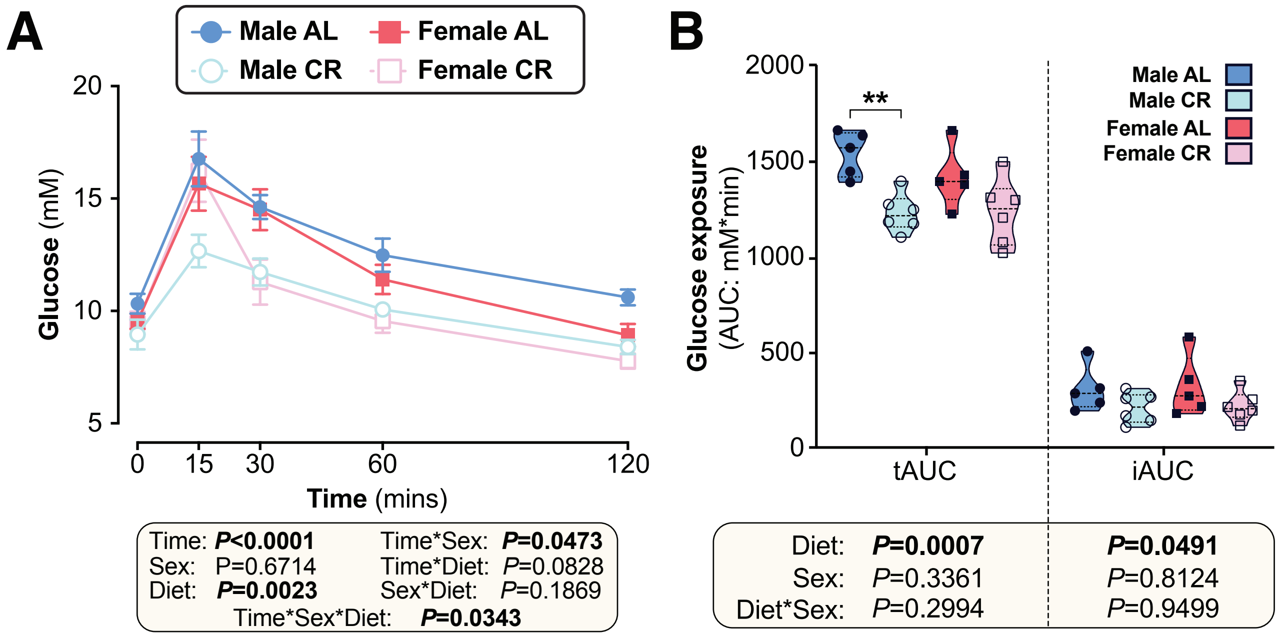

### Supplementary Figure 9

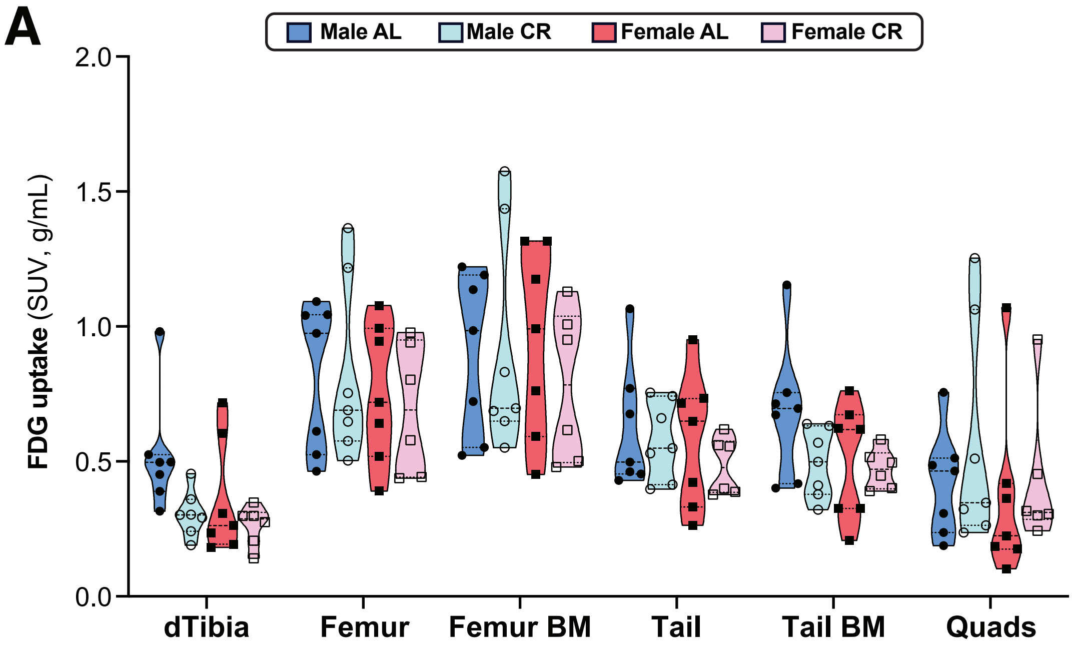


| | dTibia | Femur | Femur BM | Tail | Tail BM | Quads |
|---|---|---|---|---|---|---|
| Diet: | **0.0218** | 0.7910 | 0.5784 | 0.3608 | 0.1219 | 0.3838 |
| Sex: | 0.1120 | 0.3859 | 0.7138 | 0.3767 | 0.1639 | 0.4146 |
| Diet*Sex: | 0.3524 | 0.7916 | 0.5096 | 0.7259 | 0.2648 | 0.7337 |

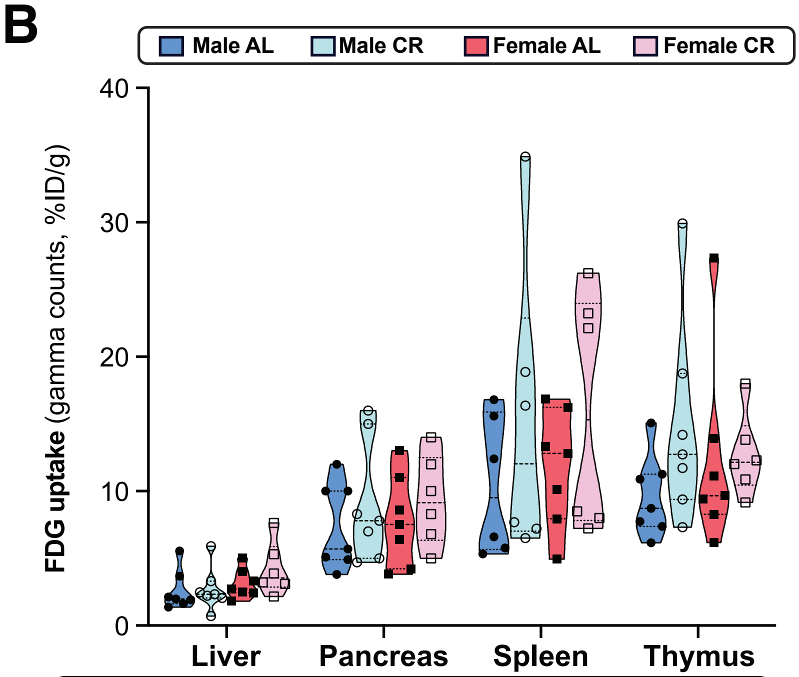


| | Liver | Pancreas | Spleen | Thymus |
|---|---|---|---|---|
| Diet: | 0.3175 | 0.2701 | 0.1583 | 0.2069 |
| Sex: | 0.1097 | 0.8173 | 0.7562 | 0.9136 |
| Diet*Sex: | 0.4065 | 0.9993 | 0.9128 | 0.2833 |

### Supplementary Figure 10

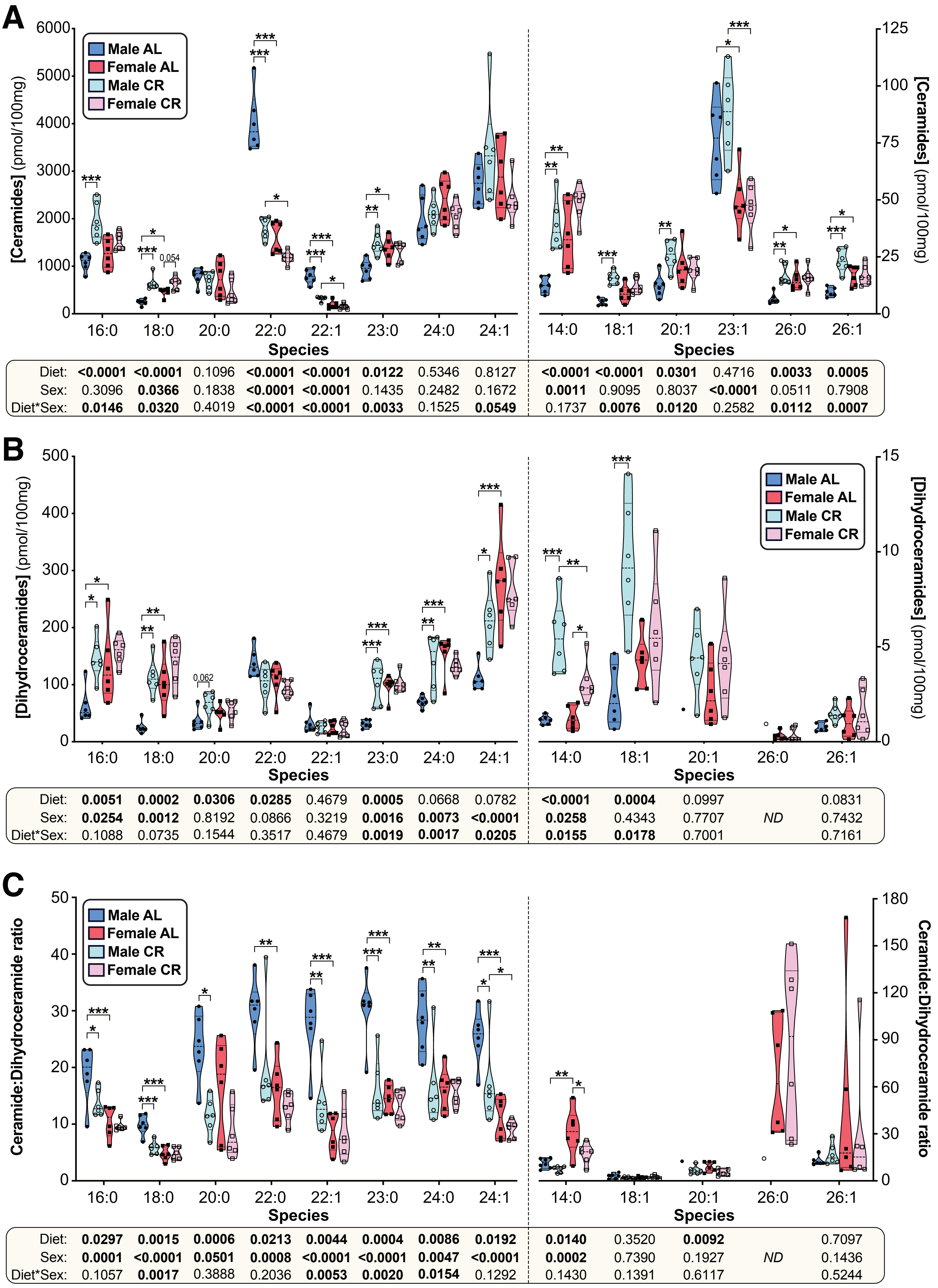


| | 16:0 | 18:0 | 20:0 | 22:0 | 22:1 | 23:0 | 24:0 | 24:1 | 14:0 | 18:1 | 20:1 | 23:1 | 26:0 | 26:1 |
|---|---|---|---|---|---|---|---|---|---|---|---|---|---|---|
| Diet: | **<0.0001** | **<0.0001** | 0.1096 | **<0.0001** | **<0.0001** | **0.0122** | 0.5346 | 0.8127 | **<0.0001** | **<0.0001** | **0.0301** | 0.4716 | **0.0033** | **0.0005** |
| Sex: | 0.3096 | **0.0366** | 0.1838 | **<0.0001** | **<0.0001** | 0.1435 | 0.2482 | 0.1672 | **0.0011** | 0.9095 | 0.8037 | **<0.0001** | 0.0511 | 0.7908 |
| Diet*Sex: | **0.0146** | **0.0320** | 0.4019 | **<0.0001** | **<0.0001** | **0.0033** | 0.1525 | **0.0549** | 0.1737 | **0.0076** | **0.0120** | 0.2582 | **0.0112** | **0.0007** |

| | 16:0 | 18:0 | 20:0 | 22:0 | 22:1 | 23:0 | 24:0 | 24:1 | 14:0 | 18:1 | 20:1 | 26:0 | 26:1 |
|---|---|---|---|---|---|---|---|---|---|---|---|---|---|
| Diet: | **0.0051** | **0.0002** | **0.0306** | **0.0285** | 0.4679 | **0.0005** | 0.0668 | 0.0782 | **<0.0001** | **0.0004** | 0.0997 | | 0.0831 |
| Sex: | **0.0254** | **0.0012** | 0.8192 | 0.0866 | 0.3219 | **0.0016** | **0.0073** | **<0.0001** | **0.0258** | 0.4343 | 0.7707 | *ND* | 0.7432 |
| Diet*Sex: | 0.1088 | 0.0735 | 0.1544 | 0.3517 | 0.4679 | **0.0019** | **0.0017** | **0.0205** | **0.0155** | **0.0178** | 0.7001 | | 0.7161 |

| | 16:0 | 18:0 | 20:0 | 22:0 | 22:1 | 23:0 | 24:0 | 24:1 | 14:0 | 18:1 | 20:1 | 26:0 | 26:1 |
|---|---|---|---|---|---|---|---|---|---|---|---|---|---|
| Diet: | **0.0297** | **0.0015** | **0.0006** | **0.0213** | **0.0044** | **0.0004** | **0.0086** | **0.0192** | **0.0140** | 0.3520 | **0.0092** | | 0.7097 |
| Sex: | **0.0001** | **<0.0001** | **0.0501** | **0.0008** | **<0.0001** | **<0.0001** | **0.0047** | **<0.0001** | **0.0002** | 0.7390 | 0.1927 | *ND* | 0.1436 |
| Diet*Sex: | 0.1057 | **0.0017** | 0.3888 | 0.2036 | **0.0053** | **0.0020** | **0.0154** | 0.1292 | 0.1430 | 0.1391 | 0.6117 | | 0.5244 |

### Supplemental Figure 11

**A**

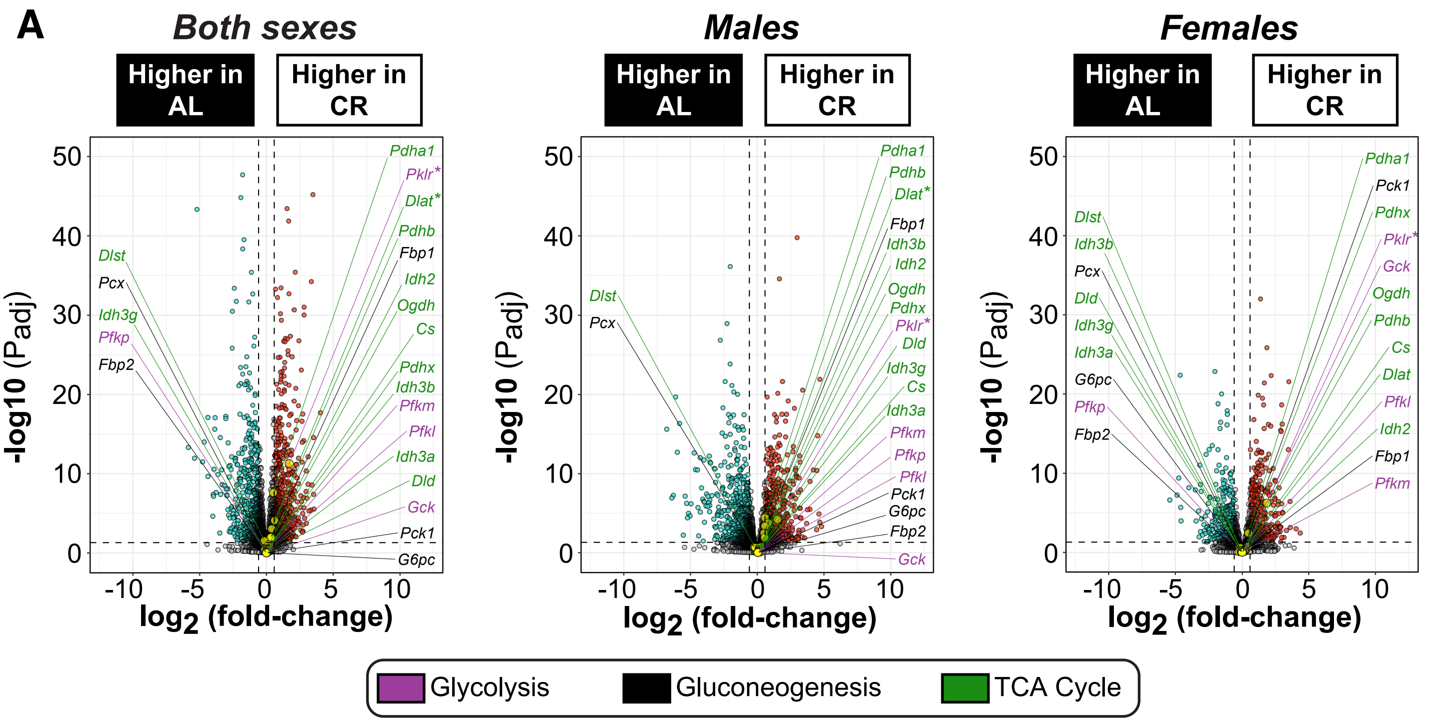


**B**

| Comparison | Gene | $Log_2$ (fold-change) | Padj |
|---|---|---|---|
| AL_F vs AL_M | *Cyp2b9* | 7.01 | 9.68E-95 |
| | *Slc22a26* | 6.92 | 8.20E-64 |
| | *Cyp2c40* | 3.85 | 6.12E-61 |
| | *Cyp2d9* | -4.51 | 7.98E-56 |
| | *Cyp2c69* | 5.67 | 8.22E-56 |
| | *Fmo2* | 3.08 | 8.02E-55 |
| | *Cyp7b1* | -3.41 | 1.65E-52 |
| CR_F vs CR_M | *Sult2a1* | 12.39 | 1.94E-19 |
| CR vs AL *(both sexes)* | *Ak4* | 1.86 | 6.71E-77 |
| | *Gm36041* | -2.15 | 5.08E-55 |
| | *Abcb1a* | 2.57 | 5.46E-51 |
| CR_M vs AL_M | *Cyp2b9* | 6.26 | 4.19E-75 |
| | *Fmo2* | 3.28 | 7.32E-62 |
| CR_F vs AL_F | *Ak4* | 2.05 | 5.60E-52 |

**C**

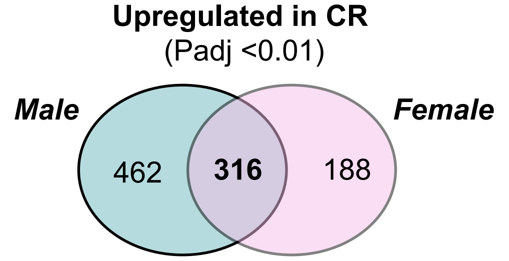


| Term | P-value | Adjusted P-value |
|---|---|---|
| Steroid biosynthesis | 2.31E-10 | 5.45E-08 |
| Biosynthesis of unsaturated fatty acids | 2.69E-08 | 3.15E-06 |
| Fatty acid biosynthesis | 4.00E-08 | 3.15E-06 |
| Terpenoid backbone biosynthesis | 1.19E-06 | 7.04E-05 |
| Circadian rhythm | 6.40E-06 | 3.02E-04 |

**D**

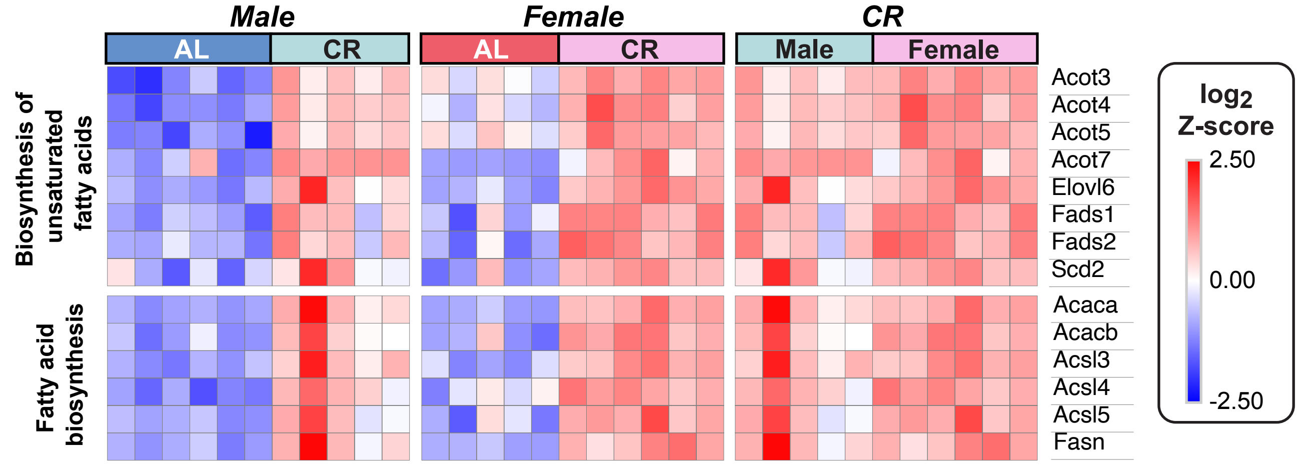

### Supplementary Figure 12

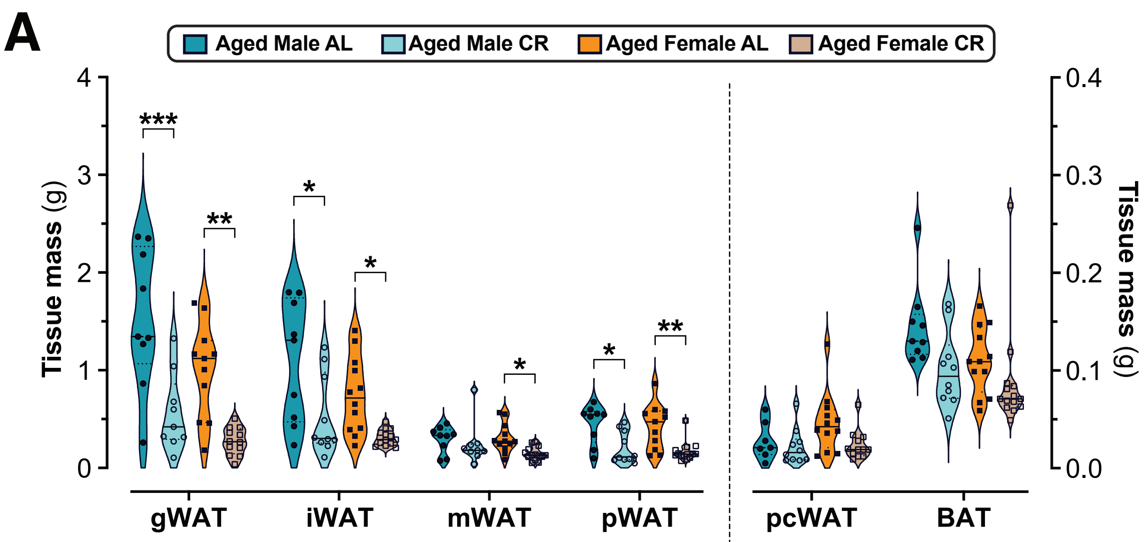


| | gWAT | iWAT | mWAT | pWAT | pcWAT | BAT |
|---|---|---|---|---|---|---|
| Diet: | **<0.0001** | **0.0002** | **0.0228** | **<0.0001** | **0.0483** | **0.0138** |
| Sex: | **0.0076** | **0.0245** | 0.3540 | 0.5178 | 0.1590 | 0.0726 |
| Diet*Sex: | 0.4167 | 0.5757 | 0.2339 | 0.8911 | 0.2095 | 0.4197 |

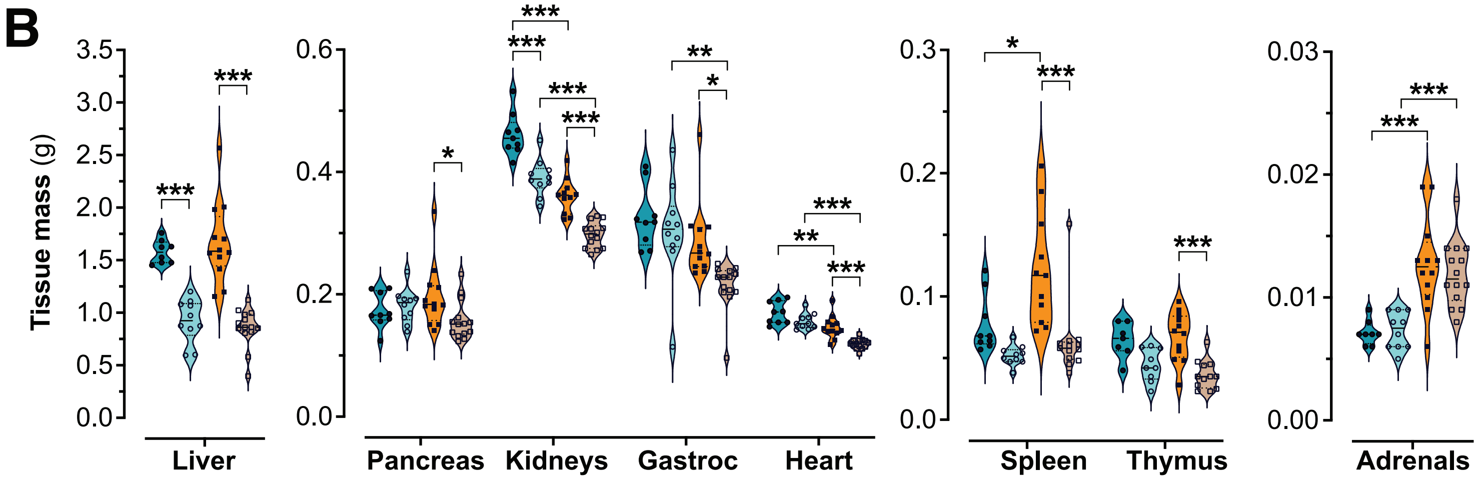


| | Liver | Pancreas | Kidneys | Gastroc | Heart | Spleen | Thymus | Adrenals |
|---|---|---|---|---|---|---|---|---|
| Diet: | **<0.0001** | 0.1460 | **<0.0001** | **0.0111** | **<0.0001** | **<0.0001** | **<0.0001** | 0.7281 |
| Sex: | 0.8879 | 0.9182 | **<0.0001** | **0.0010** | **<0.0001** | **0.0050** | 0.6235 | **<0.0001** |
| Diet*Sex: | 0.3167 | **0.0459** | 0.6647 | 0.1696 | 0.1365 | 0.0851 | 0.2733 | 0.5735 |

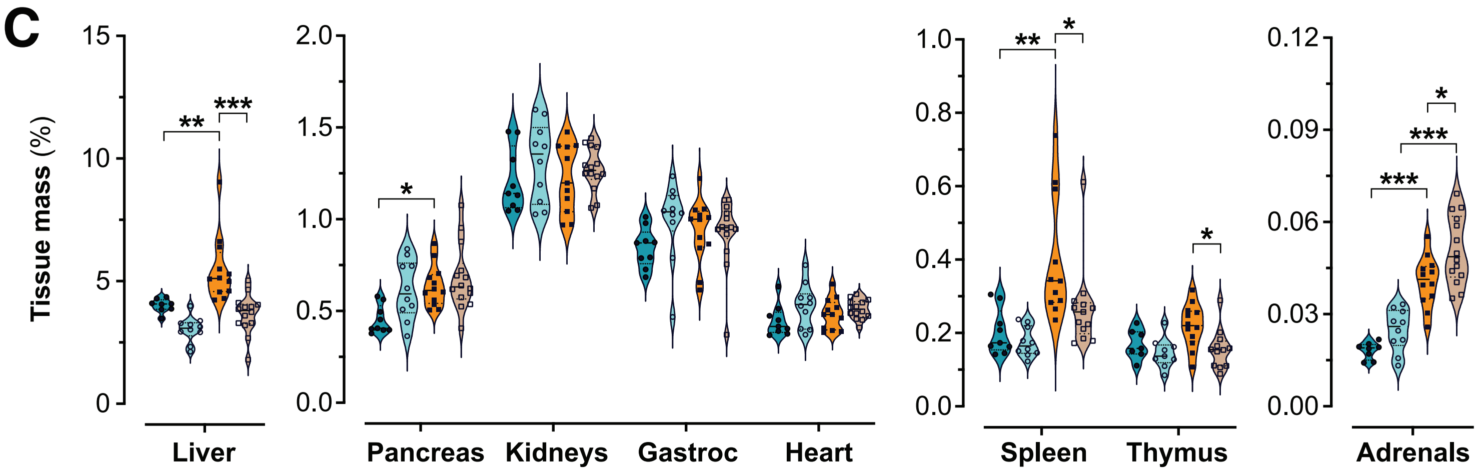


| | Liver | Pancreas | Kidneys | Gastroc | Heart | Spleen | Thymus | Adrenals |
|---|---|---|---|---|---|---|---|---|
| Diet: | **<0.0001** | **0.0213** | 0.1572 | 0.2234 | **0.0422** | **0.0240** | **0.0115** | **0.0021** |
| Sex: | **0.0004** | **0.0065** | 0.7951 | 0.8229 | 0.7450 | **0.0001** | 0.0744 | **<0.0001** |
| Diet*Sex: | 0.1267 | 0.1758 | 0.6350 | 0.1498 | 0.2881 | 0.1278 | 0.2482 | 0.4753 |

### Supplementary Figure 13

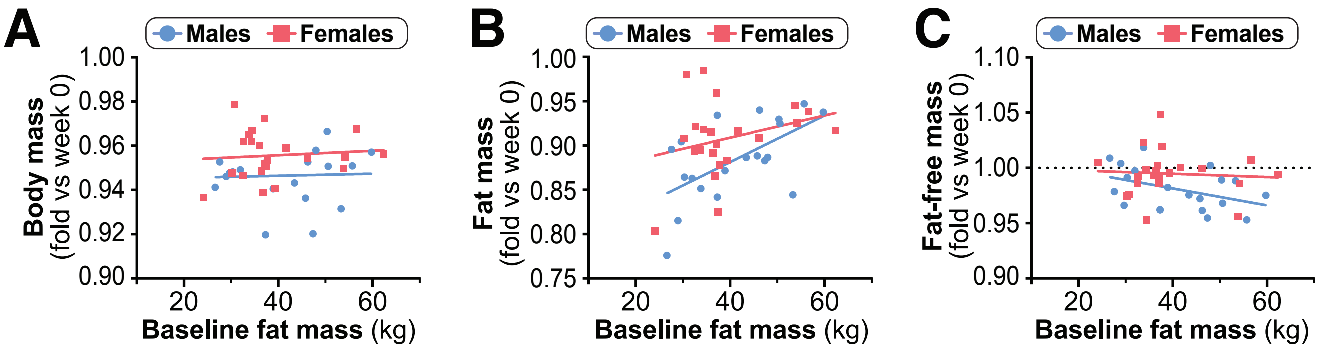


A

| | Male | Female | M vs F |
|---|---|---|---|
| *P*, Slope: | 0.872 | 0.690 | 0.887 |
| *P*, Intercept: | - | - | **0.015** |

B

| | Male | Female | M vs F |
|---|---|---|---|
| *P*, Slope: | **0.006** | 0.208 | 0.289 |
| *P*, Intercept: | - | - | **0.034** |

C

| | Male | Female | M vs F |
|---|---|---|---|
| *P*, Slope: | 0.067 | 0.759 | 0.329 |
| *P*, Intercept: | - | - | **0.032** |

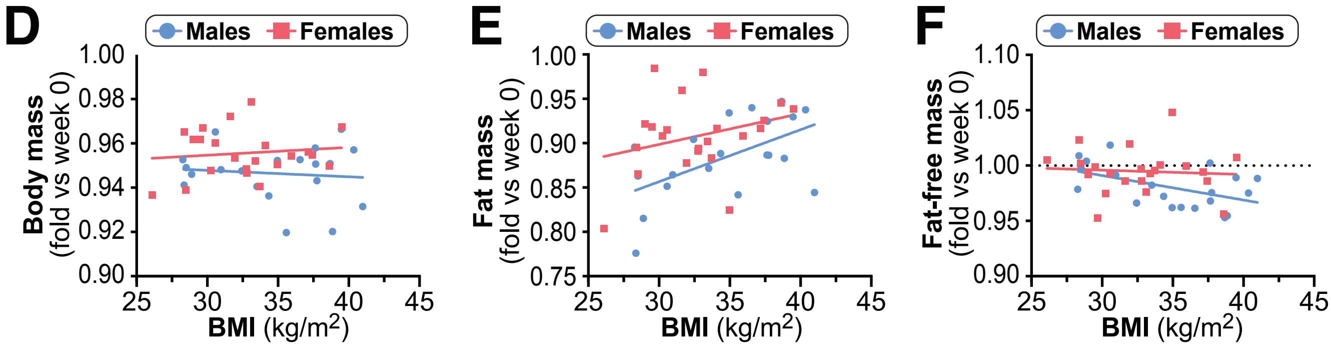


D

| | Male | Female | M vs F |
|---|---|---|---|
| *P*, Slope: | 0.676 | 0.600 | 0.509 |
| *P*, Intercept: | - | - | **0.019** |

E

| | Male | Female | M vs F |
|---|---|---|---|
| *P*, Slope: | **0.011** | 0.189 | 0.472 |
| *P*, Intercept: | - | - | **0.013** |

F

| | Male | Female | M vs F |
|---|---|---|---|
| *P*, Slope: | **0.024** | 0.762 | 0.250 |
| *P*, Intercept: | - | - | 0.072 |

### Supplementary Figure 14

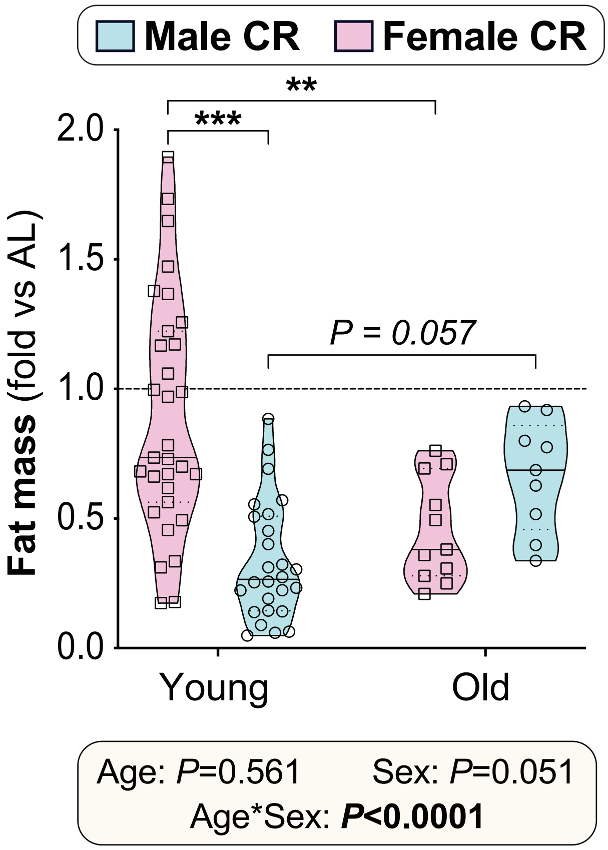

### Supplementary Figure 15

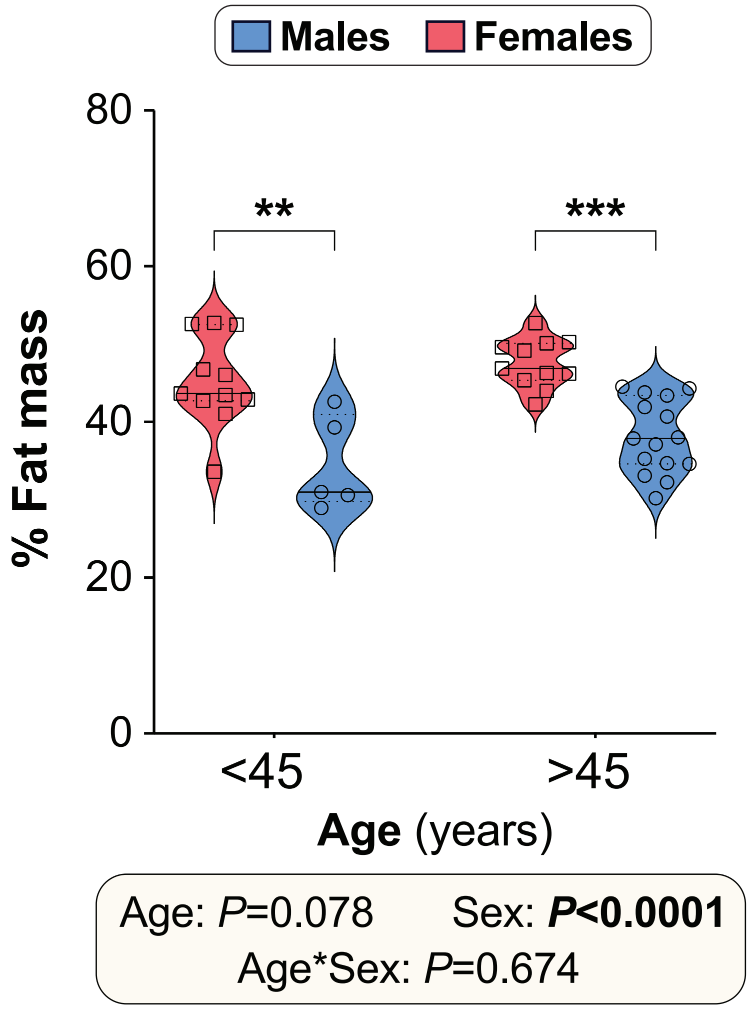
